# A meta-analysis of genome-wide association studies identifies new genetic loci associated with all-cause and vascular dementia

**DOI:** 10.1101/2022.10.11.509802

**Authors:** Bernard Fongang, Muralidharan Sargurupremraj, Xueqiu Jian, Aniket Mishra, Vincent Damotte, Joshua C Bis, Fan Kang-Hsien, Gloria Li, Jingyun Yang, Saima Hilal, M.J. Knol, Maria Pina Concas, Girotto Giorgia, Moeen Riaz, Alexander Guðjónsson, Paul Lacaze, Adam C Naj, Sven J. van der Lee, Monica Goss, Yannick W. Ngouongo, Olivia Skrobot, Vilmundur Guðnason, Lenore Launer, Oscar Lopez, Mary Haan, Ingunn Bosnes, Carole Dufouil, Mary Ganguli, Ching-Lung Cheung, David A Bennett, Christopher Chen, Kamboh M. Ilyas, Claudia Satizabal, Arfan M. Ikram, Stephanie Debette, Myriam Fornage, Yang Qiong, Gerard D. Schellenberg, Bendik Winsvold, Patrick G. Kehoe, Agustin Ruiz, Jean-Charles Lambert, Galit Weinstein, Sudha Seshadri, the Cohorts for Heart and Aging Research in Genomic Epidemiology (CHARGE)

## Abstract

Dementia is multifactorial with Alzheimer (AD) and vascular (VaD) pathologies making the largest contributions. Genome-wide association studies (GWAS) have identified over 70 genetic risk loci for AD but the genomic determinants of other dementias, including VaD remain understudied. We hypothesize that common forms of dementia will share genetic risk factors and conducted the largest GWAS to date of “all-cause dementia” (ACD) and examined the genetic overlap with VaD. Our dataset includes 809,299 individuals from European, African, Asian, and Hispanic ancestries with 46,902 and 8,702 cases of ACD and VaD, respectively. We replicated known AD loci at genome-wide significance for both ACD and VaD and conducted bioinformatic analyses to prioritize genes that are likely functionally relevant, and shared with closely related traits and risk factors. For ACD, novel loci identified were associated with energy transport (*SEMA4D*), neuronal excitability (*ANO3*), amyloid deposition in the brain (*RBFOX1*), and MRI markers of small vessel disease (*HBEGF*). Novel VaD loci were associated with hypertension, diabetes, and neuron maintenance (*SPRY2, FOXA2, AJAP1*, and *PSMA3*). Our study identified genetic risks underlying all-cause dementia, demonstrating overlap with neurodegenerative processes, vascular risk factors (Type-II diabetes, blood pressure, lipid) and cerebral small vessel disease. These novel insights could lead to new prevention and treatment strategies for all dementias.

## Introduction

Dementia is a clinical diagnosis typically occurring in late-life, characterized by a decline in cognitive performance that impairs activities of daily living and social function. Although incidence rates are stable and even declining possibly through improvement in risk control, the already substantial burden of dementia is expected to increase due to the aging of the population.^1,2^ Traditionally, Alzheimer’s disease was considered the most common dementia subtype followed by vascular dementia, and the two conditions were considered clinically distinct. Vascular dementia was diagnosed based on the presence of stroke or extensive cerebral vascular disease, with atherosclerosis and arteriolosclerosis considered the underlying pathologies.^3^ However, a wealth of evidence from recent years has emphasized a broad role for brain vascular damage, beyond that of lacunar and larger cerebral infarcts, as a major mechanism for cognitive impairment.^4-6^ It is now increasingly recognized that a component of vascular pathology is prominent in all major dementias and acts synergistically with amyloid β, tau and other neurodegenerative pathologies to affect dementia risk.^6-8^ Moreover, a new hypothetical model of dementia dynamics suggests that damage to brain vasculature, manifested as impairment in cerebral blood flow and breakdown of the blood-brain-barrier, is an early process in the dementia continuum that precedes brain atrophy, neurodegeneration, and emergence of amyloid and tau biomarker abnormalities.^4^ Recent genetic studies using methods that are relatively immune to reverse causation also suggest a putative causal relation of brain imaging markers of cerebral small vessel disease with AD.^9^

In light of the accumulating evidence stressing the importance of vascular pathology in dementia and cognitive impairment, researchers are encouraged to explore the ‘vascular contributions to cognitive impairment and dementia’ (VCID), a term that allows the inclusion of a broad range of vascular mechanisms and phenotypes and represents the multifactorial nature of dementia and related disorders as a pathway for reducing dementia burden.^10^ In particular, genetic exploration of VCID may highlight important mechanisms across the wide spectrum of pathologies, including vascular pathways, which, in turn, are considered to be the a major and modifiable target for prevention of dementia, including the Alzheimer’s type.^11^

Emerging evidence suggests that VCID is highly heritable. Monogenic conditions characterized by cerebral small vessel disease and early cognitive impairment such as Cerebral Autosomal Dominant Arteriopathy with Subcortical Infarcts and Leukoencephalopathy (CADASIL), caused by mutations in the *NOTCH3* gene,^12^ also influence later onset polygenic manifestations of VCID acting through common, less pathogenic variations in the same genes. Other examples are several point mutations in the amyloid precursor protein (APP) gene that lead to cerebral amyloid angiopathy (CAA)^13^ as well as mutations in HtrA Serine Peptidase 1(*HTRA1*) and Collagen Type IV Alpha 1 Chain (COL4A1) or COL4A2 genes.^14^ Further support for the strong genetic basis of VCID stems from heritability and genome-wide association studies (GWAS) of cerebral small vessel disease endophenotypes that are closely related with VCID, including stroke and its subtypes^15^, white matter hyperintensities (WMH),^16-19,20^ and cerebral microbleeds.^21^ In contrast to the over 70 loci identified as associated with AD genetic variance, the genetic architecture of ‘sporadic’ VCID is largely unknown. Most genetic studies of VCID have utilized a candidate gene approach, which did not yield consistent and replicable findings.^22^

Genome-wide association studies (GWAS) of VCID are sparse. In 2012, a GWAS of VaD conducted among the participants of the Rotterdam Study (N=67 cases and 5,700 controls) identified a novel locus associated with VaD, located near the androgen receptor on the X chromosome,^23^ however this finding could not be replicated.^24,25^ More recently, a GWAS of dementia and its clinical endophenotypes was conducted as part of the GR@ACE study.^26^ This study demonstrated the differential biological pathways associated with clinical AD subgroups based on the degree of vascular burden. It identified a variant near *CNTNAP2* that was associated with probable or possible VCID (N=373), however this finding did not reach genome-wide significance.

The multifactorial nature of VaD and the heterogeneity of the clinicopathological criteria used to define this entity have hampered the identification of genetic polymorphisms underlying VCID. To overcome these limitations, large-scale studies with sufficient power are needed.^27^ In the current study, we aimed to investigate the genetic predisposition of VCID. Hence, we explored the genetic variability associated with all-cause dementia as a broad phenotype, as well as vascular dementia, as an extreme phenotype of the dementia continuum characterized by increased vascular burden. Our findings from genetic exploration of VCID were then analyzed in light of the knowledge already gained from previous large-scale genome-wide and sequencing studies on the genetic determinants of Alzheimer’s disease, stroke and additional phenotypes along the VCID spectrum.^12-15^

## Results

Our analysis included 809,299 individuals comprising 46,902 and 8,702 cases of ACD and VaD, respectively. They were recruited from 19 CHARGE cohorts, the ADGC, the EADB, and the UK Biobank, encompassing four different reported ancestries: European (98.5%), African (1.0%), Asian (0.4%), and Hispanics/Latino (0.1%). Association analyses were performed in each cohort following a pre-defined analysis plan, using logistic regression and Cox proportional hazards models for prevalent and incident cases, respectively. We performed study-specific quality control (QC) of the summary statistic data, followed by ancestry-specific meta-analyses and trans-ethnic meta-analyses of ACD and VaD as described in the Methods section. For each cohort description, association analysis method, QC parameters, and cutoffs are provided in the **Supplementary file SF3**.

### Meta-analyses of ACD and VaD GWAS in European ancestry populations replicated known AD loci

We conducted fixed-effects inverse variant-weighted meta-analyses of the 14 European ancestry cohorts (N = 466,606, N_ACD_ = 44,009, N_VaD_ = 3,892) from the Cohorts for Heart and Aging Research in Genomic Epidemiology (*CHARGE*) consortium, the Alzheimer’s Disease Genetics Consortium (*ADGC*).^28^, and the UK Biobank (UKBB)^29^. Furthermore, we replicated significant and suggestive signals from our VaD GWAS in the European Alzheimer Disease Biobank (EADB) consortium VCID data (N = 275,745, N_VaD_ = 4,564). For UKBB, we utilized the proxy-AD phenotype defined by Marioni et al.^30^ for ACD and defined VaD using the ICD10 codes, as explained in the methods section. Briefly, the proxy-AD phenotype identified 35,214 individuals aged at least 65 years, reporting a history of dementia in one or both of their parents as ‘cases’ and 180,791 individuals with >= 65 with no history of dementia in either biological parents serving as controls. The complete list of cohorts included in this study is provided in **Supplementary File SF1 [Table ST1]**.

A total of 11,596,629 and 9,878,961 SNPs passed the study-level QC criteria and were tested for association with ACD and VaD, respectively. After post meta-analysis quality control, we identified 10 genome-wide significant loci associated with ACD (GWS, P<5 × 10^−8^), all of which have been previously associated with AD (**Table 2**, extended results in **Supplementary File SF1 [Table ST2]**). Significant loci associated with ACD included signals in or around known AD gene such as *APOE, BIN1, MS4A6A, PICALM, CR1, CD2AP, ABCA7, PILRB, SLC24A4*, and *ACE*. For VaD, only one variant located near the *APOE* gene reached genome-wide significance (**Table 3**, extended results in **Supplementary File SF1 [Table ST3]**). The genomic inflation coefficients (lambda) were 1.05 and 1.07 for ACD and VaD respectively, the lambda intercept computed with the LDSC software was 1.01 for both analyses, suggesting no systematic inflations of associations statistics. The Manhattan and quantile-quantile (QQ) plots for both analyses are provided in **Figures 1-3** (Forest and locusZoom plots for prominent signals are provided in **Supplementary File SF2 [Figure 7-24])**.

**Table 1:**
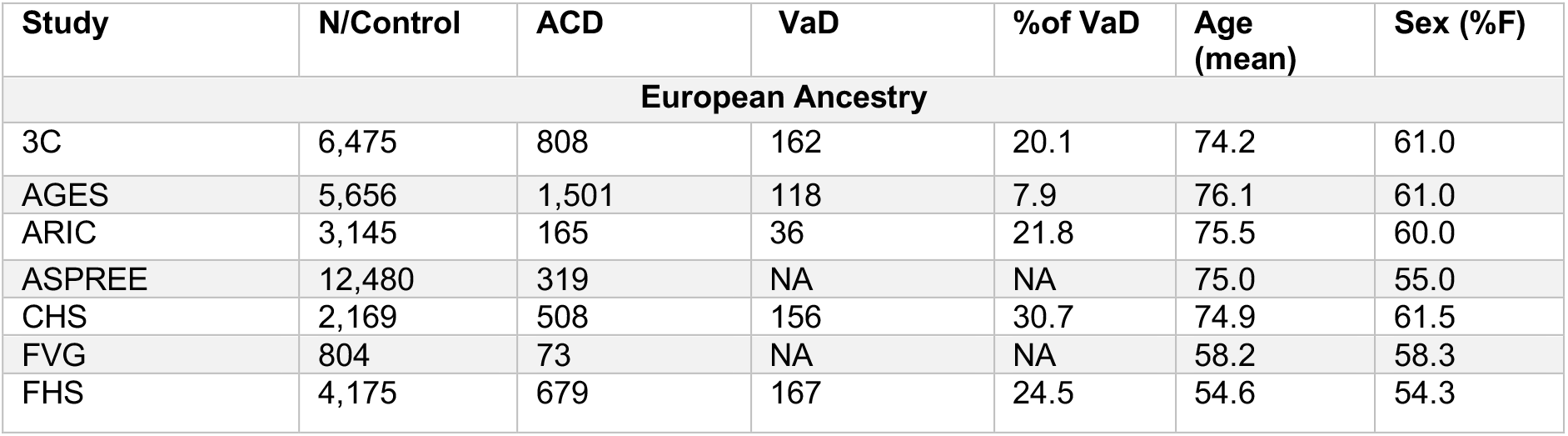
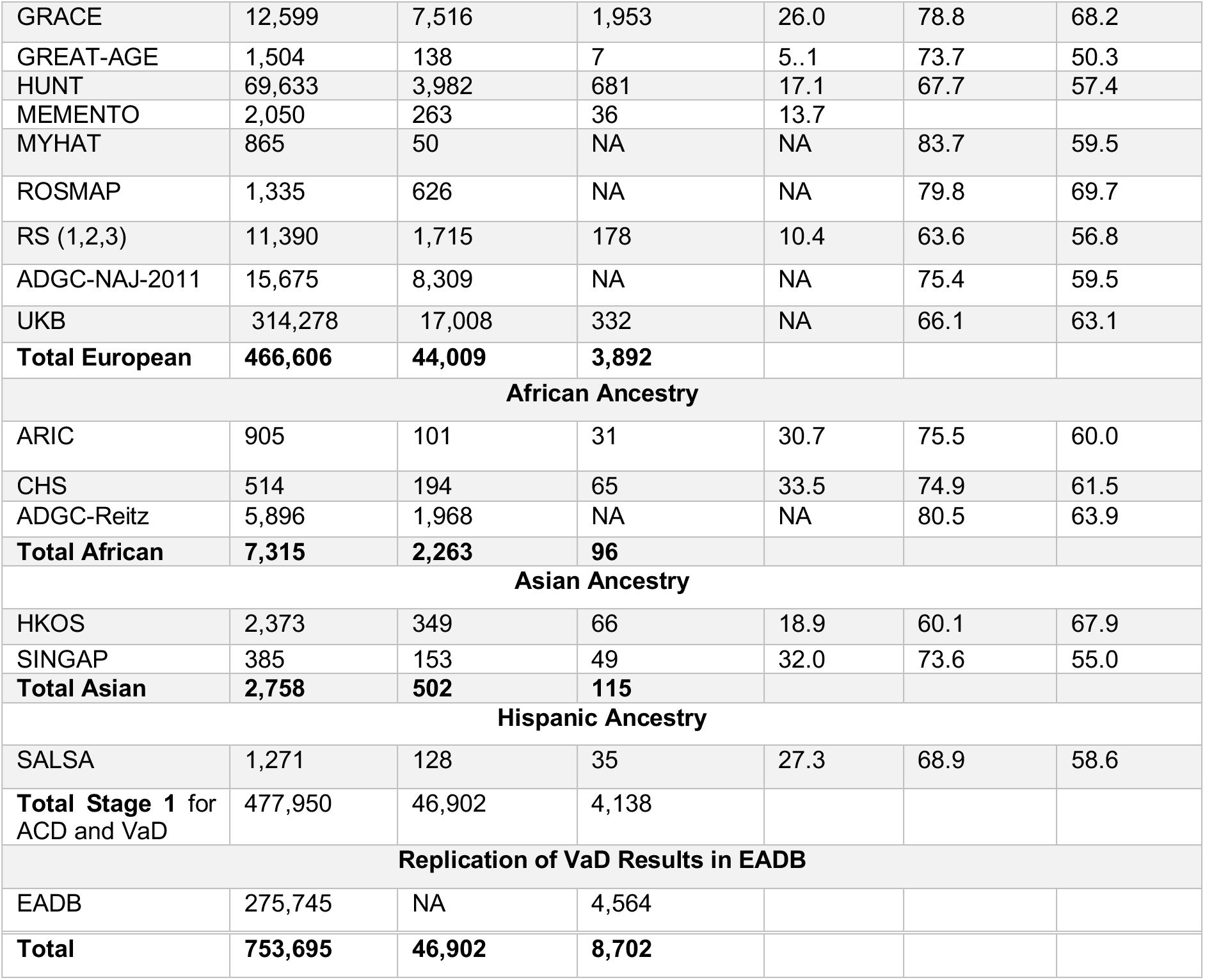
Demographics: Data from 17 CHARGE cohorts were included in our meta-analysis as well as the UKB, ADGC, and EADB for the replication of our VaD results in European ancestry. Overall, 809,299 individuals were included the meta-analysis accounting for 46,902 and 8,702 cases of ACD and VaD respectively. For UKB, we used the proxy-AD (familiar AD) for ACD analysis and assess VaD cases using ICD10 codes (see Methods). We also used ADGC-NAJ-2011 and ADGC-Reitz for ACD in European and African ancestry, respectively to avoid overlap with CHARGE samples. We subsequently replicated our VaD results in EADB.

| Study | N/Control | ACD | VaD | %of VaD | Age (mean) | Sex (%F) |
| --- | --- | --- | --- | --- | --- | --- |
| **European Ancestry** |  |  |  |  |  |  |
| 3C | 6,475 | 808 | 162 | 20.1 | 74.2 | 61.0 |
| AGES | 5,656 | 1,501 | 118 | 7.9 | 76.1 | 61.0 |
| ARIC | 3,145 | 165 | 36 | 21.8 | 75.5 | 60.0 |
| ASPREE | 12,480 | 319 | NA | NA | 75.0 | 55.0 |
| CHS | 2,169 | 508 | 156 | 30.7 | 74.9 | 61.5 |
| FVG | 804 | 73 | NA | NA | 58.2 | 58.3 |
| FHS | 4,175 | 679 | 167 | 24.5 | 54.6 | 54.3 |
| GRACE | 12,599 | 7,516 | 1,953 | 26.0 | 78.8 | 68.2 |
| GREAT-AGE | 1,504 | 138 | 7 | 5.1 | 73.7 | 50.3 |
| HUNT | 69,633 | 3,982 | 681 | 17.1 | 67.7 | 57.4 |
| MEMENTO | 2,050 | 263 | 36 | 13.7 |  |  |
| MYHAT | 865 | 50 | NA | NA | 83.7 | 59.5 |
| ROSMAP | 1,335 | 626 | NA | NA | 79.8 | 69.7 |
| RS (1,2,3) | 11,390 | 1,715 | 178 | 10.4 | 63.6 | 56.8 |
| ADGC-NAJ-2011 | 15,675 | 8,309 | NA | NA | 75.4 | 59.5 |
| UKB | 314,278 | 17,008 | 332 | NA | 66.1 | 63.1 |
| **Total European** | **466,606** | **44,009** | **3,892** |  |  |  |
| **African Ancestry** |  |  |  |  |  |  |
| ARIC | 905 | 101 | 31 | 30.7 | 75.5 | 60.0 |
| CHS | 514 | 194 | 65 | 33.5 | 74.9 | 61.5 |
| ADGC-Reitz | 5,896 | 1,968 | NA | NA | 80.5 | 63.9 |
| **Total African** | **7,315** | **2,263** | **96** |  |  |  |
| **Asian Ancestry** |  |  |  |  |  |  |
| HKOS | 2,373 | 349 | 66 | 18.9 | 60.1 | 67.9 |
| SINGAP | 385 | 153 | 49 | 32.0 | 73.6 | 55.0 |
| **Total Asian** | **2,758** | **502** | **115** |  |  |  |
| **Hispanic Ancestry** |  |  |  |  |  |  |
| SALSA | 1,271 | 128 | 35 | 27.3 | 68.9 | 58.6 |
| **Total Stage 1 for ACD and VaD** | **477,950** | **46,902** | **4,138** |  |  |  |
| **Replication of VaD Results in EADB** |  |  |  |  |  |  |
| EADB | 275,745 | NA | 4,564 |  |  |  |
| **Total** | **753,695** | **46,902** | **8,702** |  |  |  |

**Table 2.**
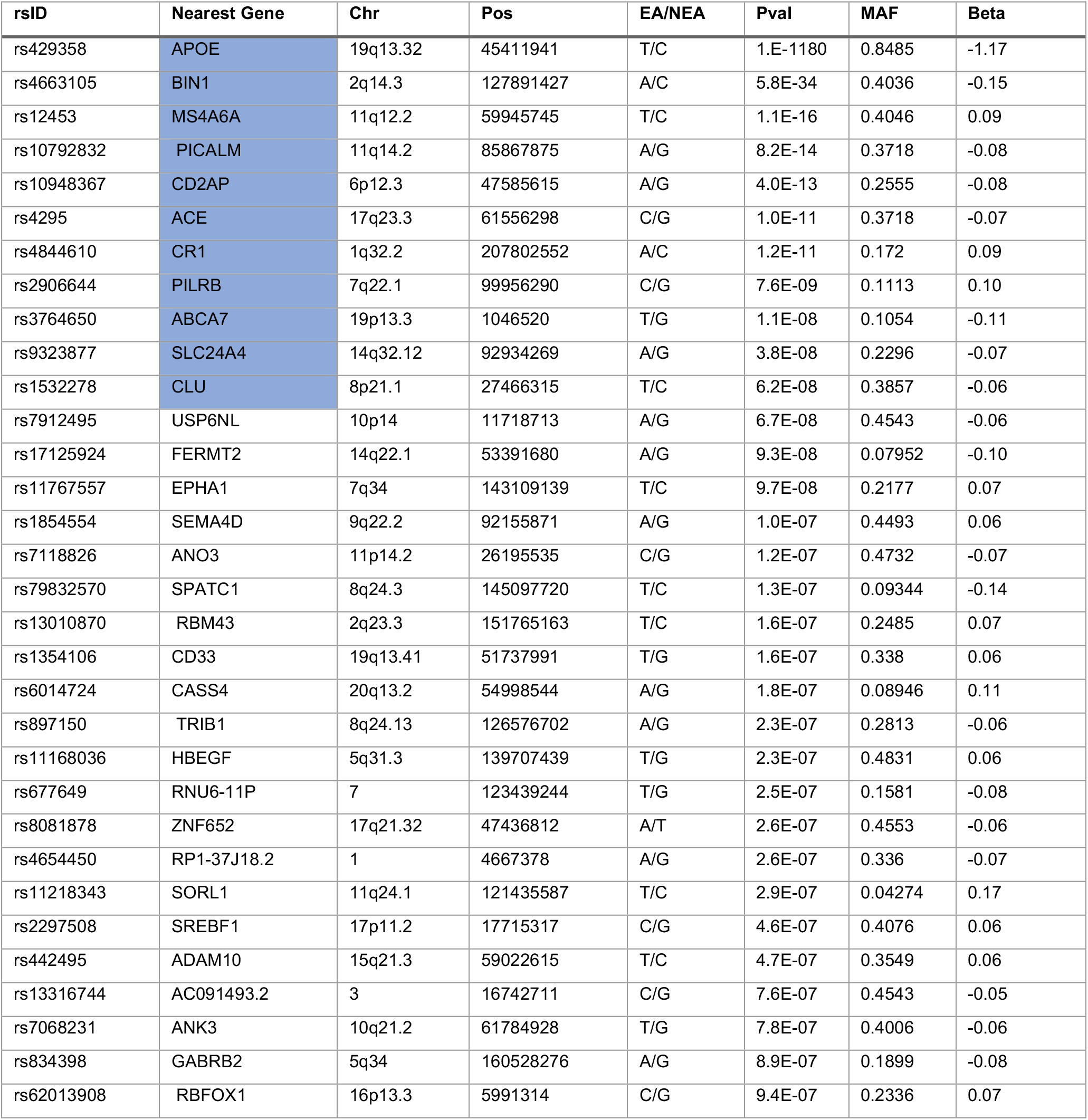
Genome-wide significant (P<5×10-8) and suggestive (P<1×10-6) variants associated with all-cause dementia in European population.

**Table 3.**
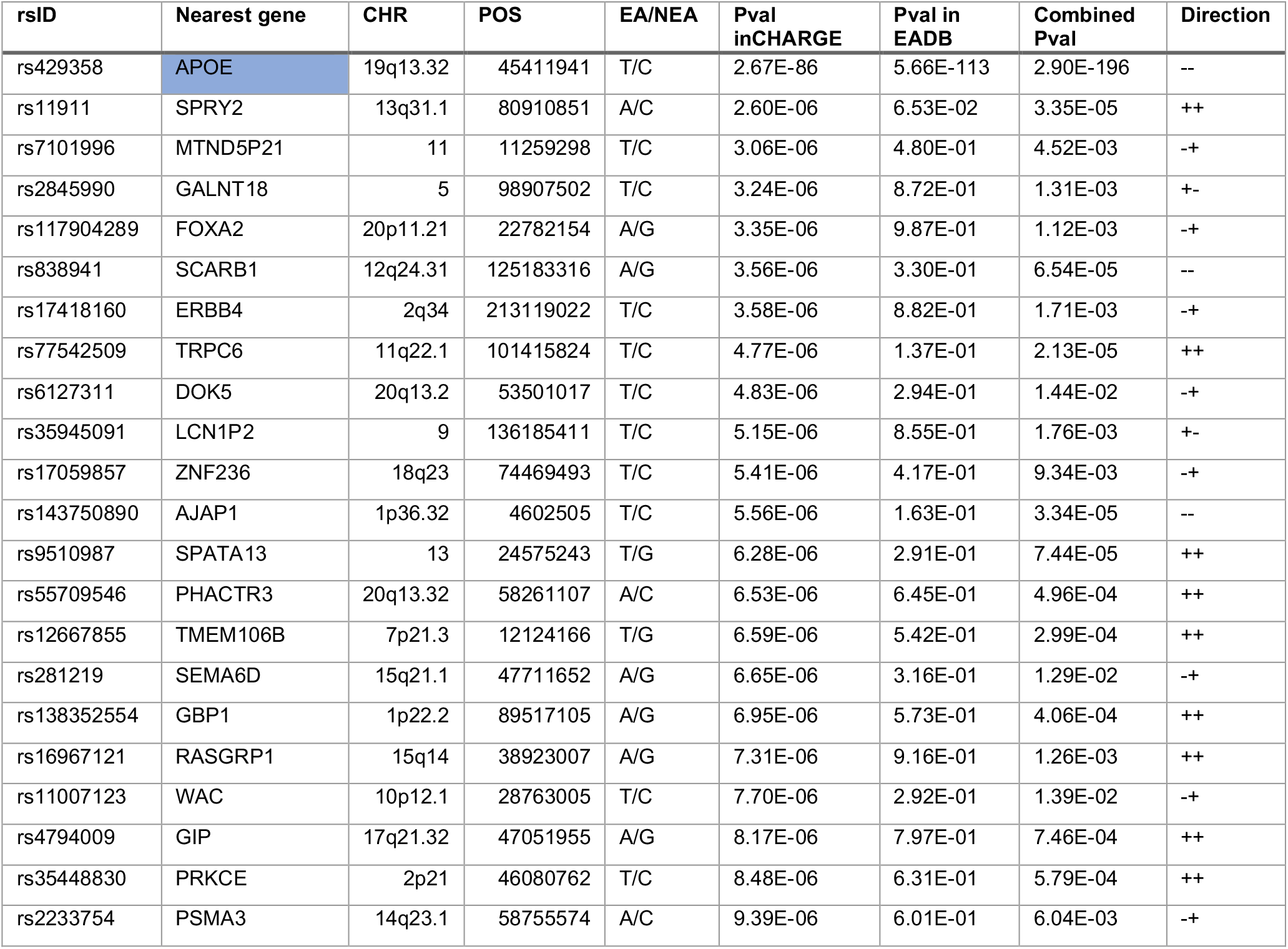
Genome-wide significant (P<5×10^−8^) and suggestive (P<1×10^−6^) variants associated with vascular dementia in European population. The meta-analysis includes 11 cohorts from the CHARGE consortium, and the UK Biobank (UKBB) GWAS. Direction denotes the direction of association in CHARGE and EADB

**Figure 1:**
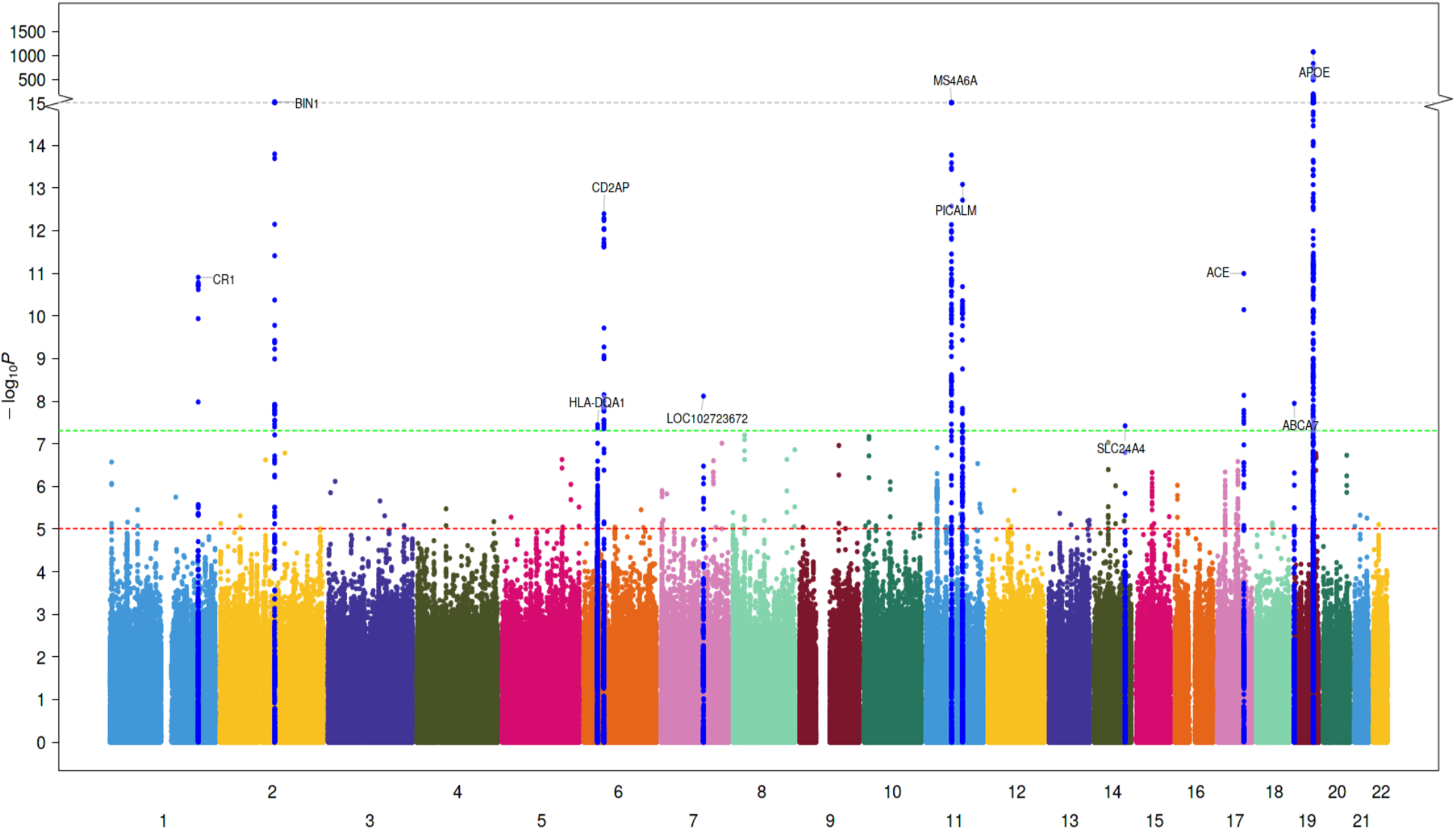
Manhattan plot of the ACD GWAS. In addition to variants in the APOE region, we identified five new genetic loci associated with VaD. Blue and red lines correspond to the p-value of 5e^-7^ and 5e^-8^ for genome-wide suggestive and significant SNPs, respectively. Manhattan plots for the cross-ancestry meta-analysis. Each dot represents a SNP, the X-axis shows the chromosomes where each SNP is located, and the Y-axis shows -log10 P-value of the association of each SNP with POAG in the cross-ancestry meta-analysis. The red horizontal line shows the genome-wide significant threshold (P-value=5e-8; -log10 P-value=7.30). The nearest gene to the most significant SNP in each locus has been labeled

**Figure 2:**
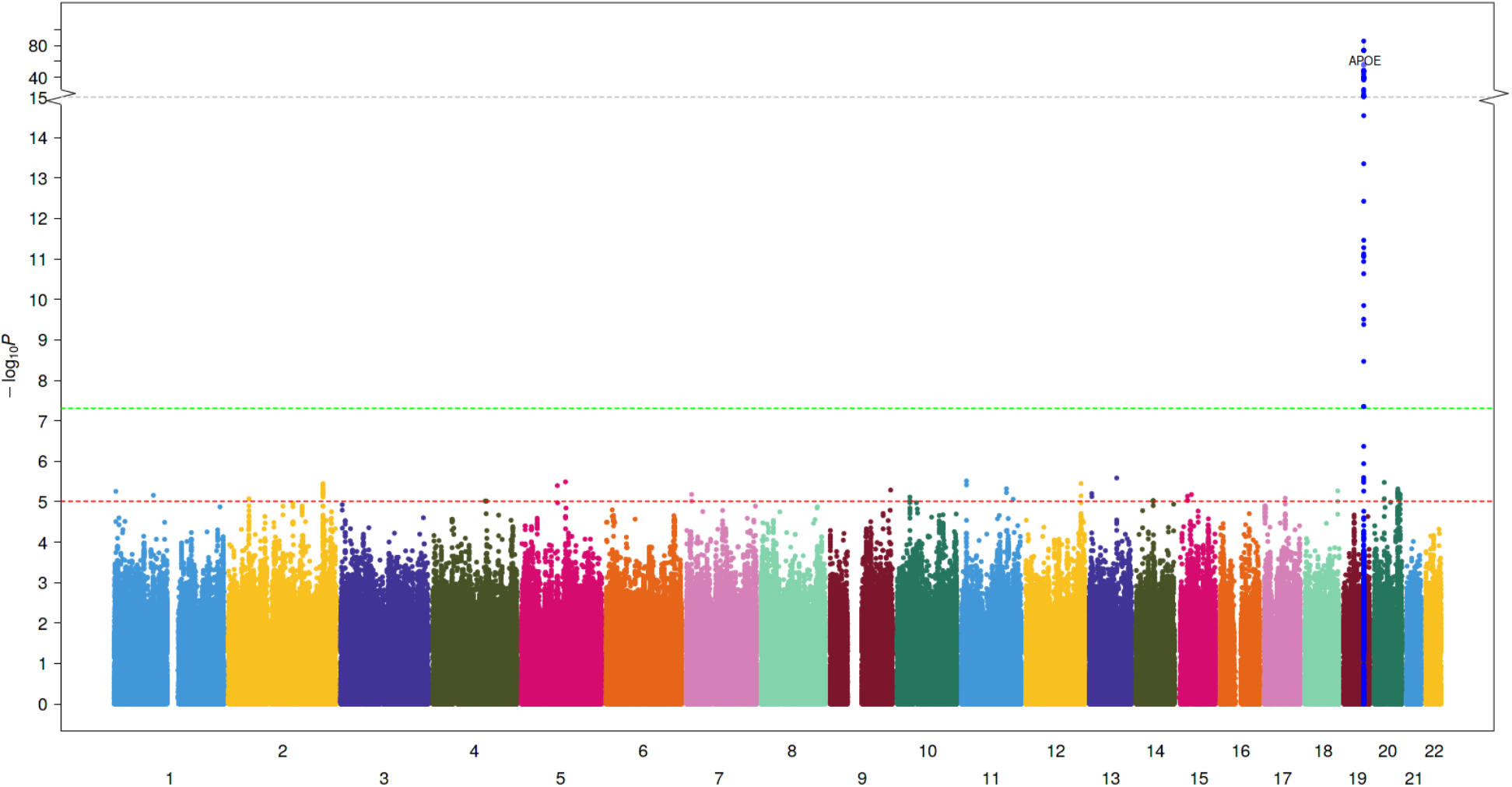
Manhattan plot of the VaD GWAS. In addition to variants in the APOE region, we identified five new genetic loci associated with VaD. Blue and red lines correspond to the p-value of 5e^-7^ and 5e^-8^ for genome-wide suggestive and significant SNPs, respectively. Manhattan plots for the cross-ancestry meta-analysis. Each dot represents a SNP, the X-axis shows the chromosomes where each SNP is located, and the Y-axis shows -log10 P-value of the association of each SNP with POAG in the cross-ancestry meta-analysis. The red horizontal line shows the genome-wide significant threshold (P-value=5e-8; -log10 P-value=7.30). The nearest gene to the most significant SNP in each locus has been labeled

**Figure 3:**
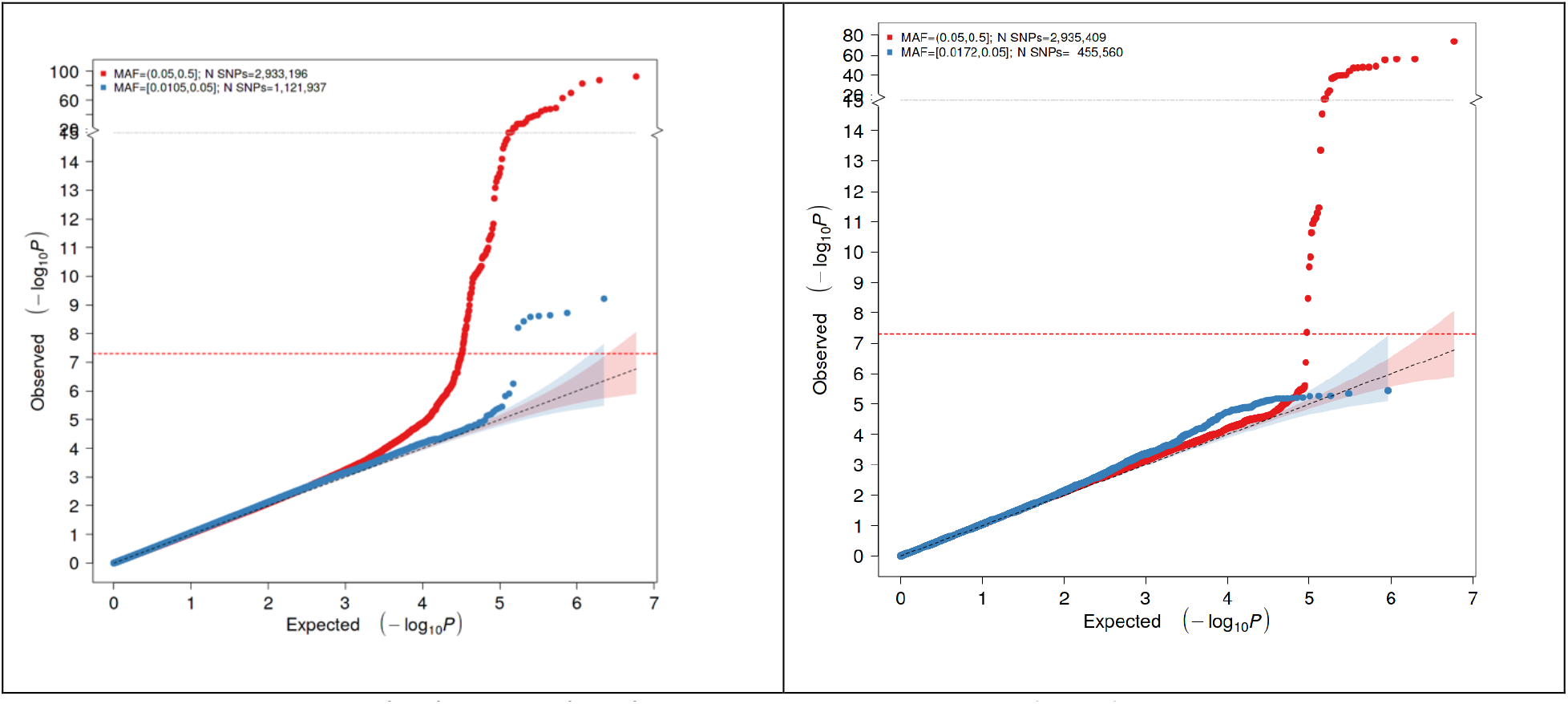
Q-Q plots of the ACD (left) and VaD (right) GWASs. The expected P values (X-axis) are plotted against the observed P-values (Y-axis). The units of the axes are the –log10 of the P-value. The red and blue curves represent the plots with MAF >= 0.05 and 0.01 respectively. The diagonal line of the null hypothesis and its 95% confidence interval were plotted in grey based on the P-values without the previously reported SNPs. The red dot line represents the cutoff for genome-wide significance

For the VaD trait, we selected all variants with p-value less than 1×10^−5^ and meta-analyzed with EADB summary results using the p-value based approach. Only one variant near the APOE gene was statistically significant and has the same direction of effect in both studies (**Table 3**).

The meta-analyses of ACD and VaD GWAS in African, Asian, and Hispanic/Latino ancestries did not provide new genome-wide significant variants.

### Trans-Ancestral meta-analysis of ACD and VaD GWAS

We next performed trans-ancestral meta-analysis, first to assess whether the increased sample size could lead to the identification of additional loci associated with ACD and VaD, and also to identify loci that are relevant in other ancestries. The trans-ancestral meta-analyses included individuals from European, African, Asian, and Hispanic/Latino ancestries. The total Number of variants included were 17,054,226 and 11,595,061 for ACD and VaD, respectively. The Manhattan plots of the SNP-wide meta-analyses for both traits are provided in **Supplementary File SF2 [Figures 4-5]**. Significant and suggestive signals for ACD and VaD are presented in **Supplementary File SF2 [Table 3-4] and the extended results in Supplementary File SF1 [Table ST4-ST5]**. We identified novel signals reaching genome-wide significance at 20q11.21 (*CHD6*, an oxidative DNA damage response factor previously associated with neurological phenotype),^31,32^ 2q14.1 (*DAW1*, involved in cerebrospinal fluid circulation and cilia motility during development),^33^ and 15q15.1 (*PWRN2*, previously associated with tauopathy and Prader-Willi syndrome)^34,35^ for ACD and 17q21.1 (*MARCHF10*) for VaD.

**Table 4.**
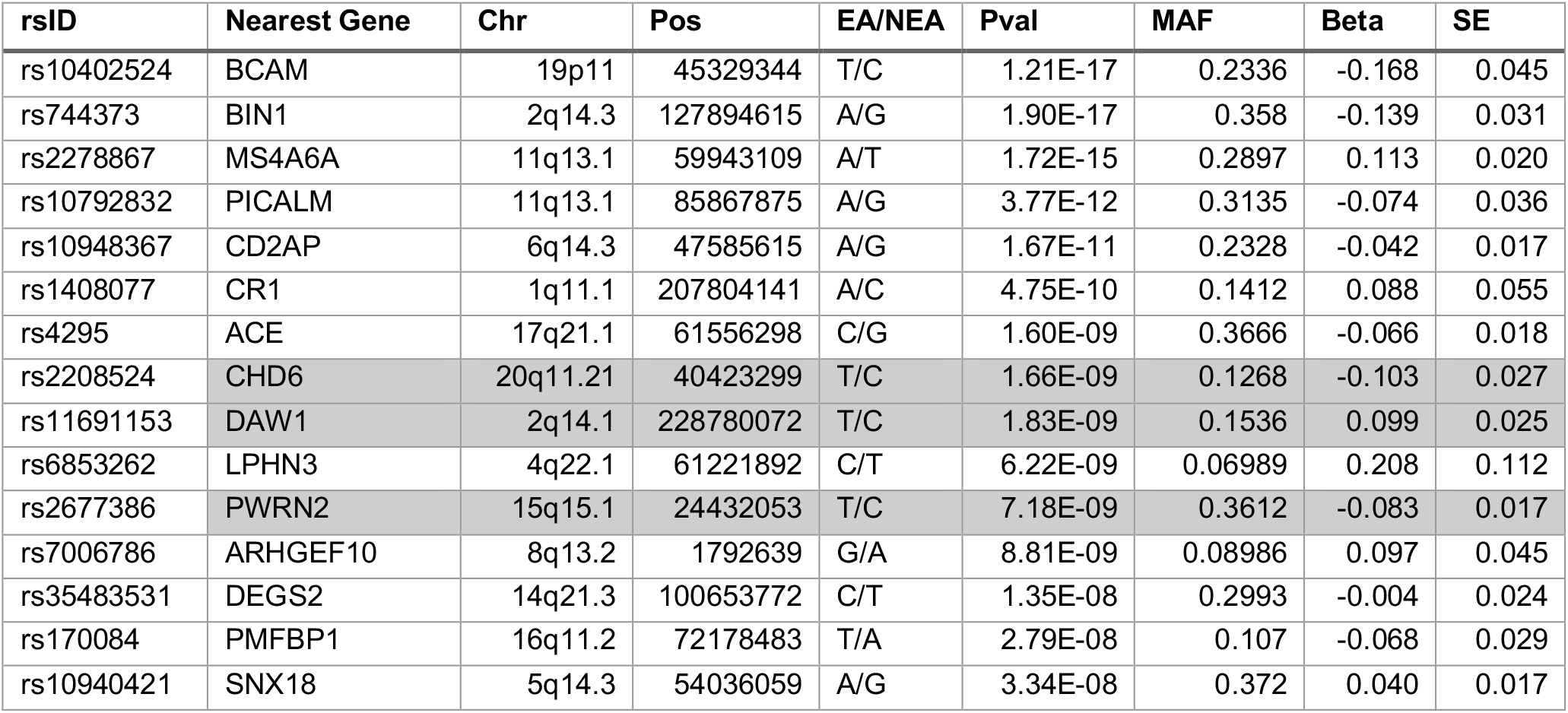
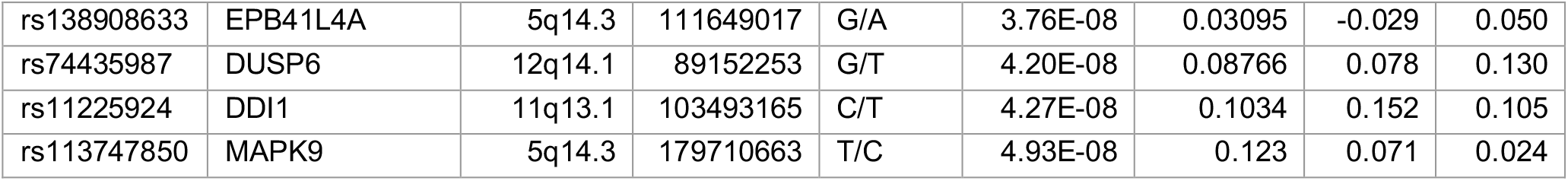
Genome-wide significant (P<5×10^−8^) and suggestive (P<1×10^−6^) variants associated with all-cause dementia in trans-ancestral meta-analysis. The meta-analysis includes European, African, Asian, and Hispanic/Latino ancestries. Three new variants at 20q11.21, 2q14.1, and 15q15.1 reached genome-wide significance.

**Table 5.**
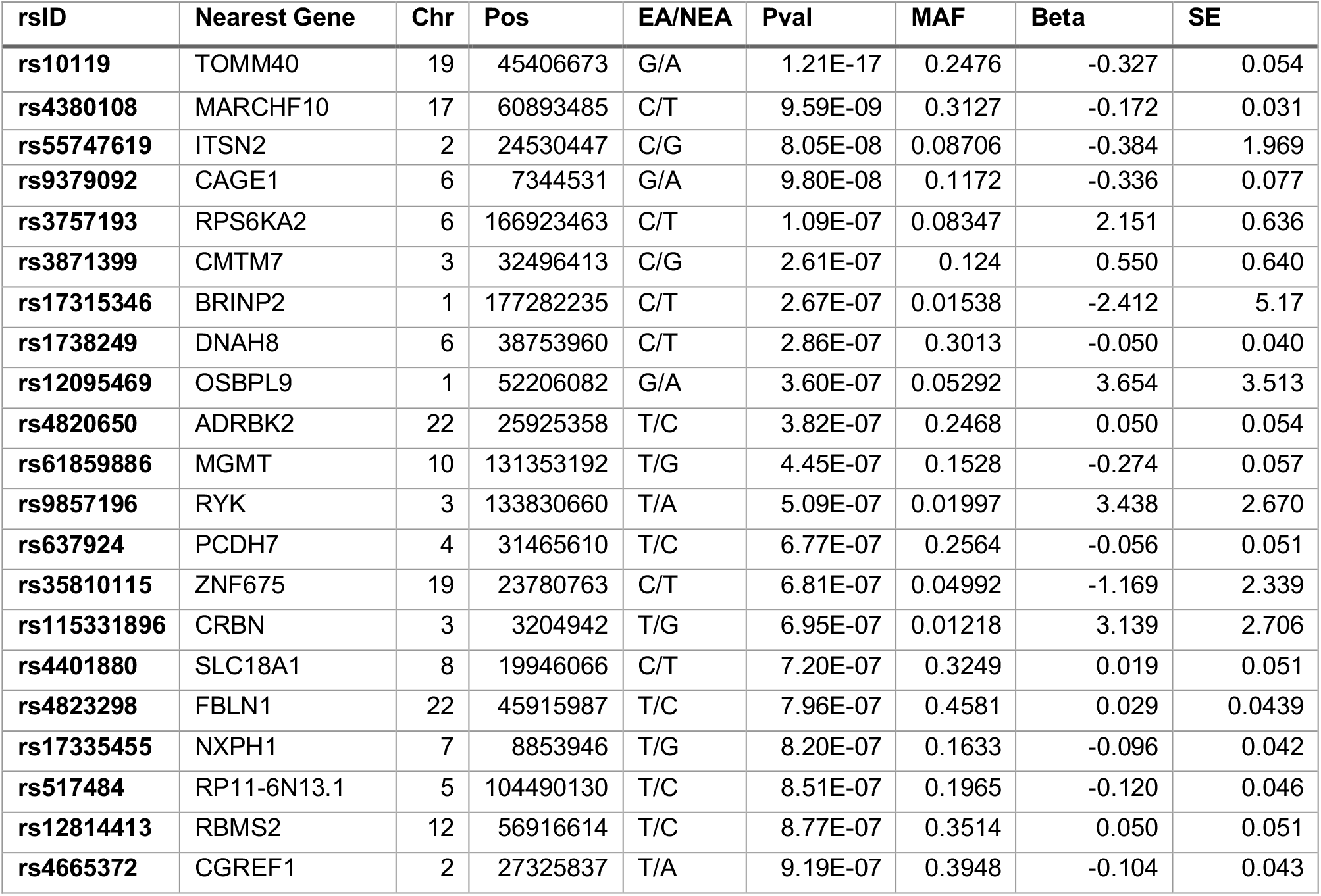
Genome-wide significant (P<5×10^−8^) and suggestive (P<1×10^−6^) variants associated with vascular dementia in trans-ancestral meta-analysis. The meta-analysis includes European, African, Asian, and Hispanic/Latino ancestries. One new variant at

## Functional characterization of GW suggestive signals for ACD and VaD meta-analyses

### Shared genetic susceptibility with complex disease traits

Substantial shared genetic susceptibility of ACD and VaD with risk factors and complex disease traits is evident across different genomic scales (single variant, regional and the global level). ACD exhibits genetic pleiotropy with vascular risk factors (hypertension, white-matter hyperintensity burden; WMH), hematological traits (neutrophil, lymphocyte count) and blood-based biomarkers indicative of inflammation (C-reactive protein levels), hemostasis (fibrinogen, factor-VII levels) and neurodegeneration (soluble *TREM2* levels) (**Supplementary File SF1 [Table ST6]**). This shared genetic susceptibility is primarily driven by the *MS4A* gene famil*y* (membrane-spanning 4A; *MS4A6A, MS4A4A*). The sharing of common genetic variation between ACD and vascular risk factors [blood pressure traits (DBP, SBP, PP), and T2D] at the *ACE* and *PILRB* locus (**Supplementary File SF2 [Figure 25], Supplementary File SF1 [Table ST6]**) is further supported by our regional Bayesian pairwise (GWAS-PW) analysis highlighting the high-probability (see methods) of harboring a shared causal variant (**Supplementary File SF1 [Table ST7]**). Interestingly, the GWAS-PW approach additionally reveals the shared genetic susceptibility of VaD with ischemic stroke (IS) and WMH at the *PRPF8* and *PRDM6* locus (**Supplementary File SF1 [Table ST7]**). In support, global level genetic correlation analysis (excluding *APOE* region) using LDSR confirmed the association of increased levels of WMH (borderline) with increased risk of ACD (**Supplementary File SF1 [Table ST8]**). Additionally, our LDSR analysis observed an inverse association of high levels of high-density lipoprotein (protective) and cardio-embolic stroke with the ACD risk. As expected, a strong genetic correlation of poorer cognitive performance (general cognitive function; GCF) with ACD was also observed. The impact of possible sample overlap was estimated to be negligible using LDSR (**Supplementary File SF1 [Table ST8]**).

### Polygenicity and multi-trait analysis

To identify additional SNPs conferring susceptibility to ACD and acting through related risk factors, we jointly studied the genome-wide distribution of genetic effects for ACD and its closely related traits. We first prioritized those traits which have a polygenic background similar to ACD using a Bayesian approach (*ashR*, see methods). The *ashR* analysis showed that certain traits (ACD, stroke and its subtypes, WMH, coronary artery disease [CAD]) had specific, possibly overlapping pathophysiological processes, compared to other ACD risk factors (SBP, smoking-SMK, BMI) that involved multiple biological pathways (**Supplementary File SF2 [Figure 26]**). Second, using multi-trait GWAS analysis (MTAG, see methods) on ACD and the prioritized traits (CAD, stroke, WMH), we identified intronic SNPs in *SMG6* and *ABCG8* to be genome-wide significant (pMTAG < 1.67E-08, for 3 phenotypes) for ACD and possibly acting through CAD (**Supplementary File SF1 [Table ST9]**). Interestingly, *SMG6* may also have a role in tau biology.^36^

### Functional prioritization using molecular profile (gene expression)

To functionally characterize and prioritize individual ACD and VaD genomic risk loci, we performed transcriptome-wide association studies (TWAS) using TWAS-Fusion, ACD and VaD association statistics and weights from 23 gene-expression reference panels from blood, arterial, and brain tissues (see methods). We identified 29 trait-associated (ACD/VaD) SNPs functioning as expression quantitative trait loci (eQTL), regulating the expression of 22 genes (eGenes) in disease-relevant tissue types (**Supplementary File SF1 [Table ST10]**). To explore whether the observed associations are real or merely reflect the random overlap between eQTLs and non-causal risk variants for the dementia traits, a colocalization analysis was performed at each significant locus estimating the posterior probability of a shared causal variant (PP4 ≥ 75%) between the gene expression and trait association. Overall, 30% of the eQTL-eGene satisfied the colocalization threshold for a causal variant being shared between the ACD or VaD and gene expression. In addition to fine mapping functional genes (*RP11-385F7*.*1, CR1, MS4A6A, ACE, APOC4*) in the loci exhibiting genome-wide association with ACD/VaD, TWAS identified putative novel (*CLU-*ACD, *PIKFYVE-*VaD, *SH3D21-*ACD) genes satisfying transcriptome-wide significance threshold (pTWAS < 1.18E-05) and the colocalization probability threshold. The majority (91%) of the eGenes, are supported by the positional overlap of corresponding eQTLs with regulatory marks (enhancer and promoter binding sites) for active transcription in relevant tissue types.

### Identification of independent case-case loci with CC-GWAS

In additional to genetic overlap, we explored genetic difference between ACD/VaD and AD using case-case GWAS (see methods). We identified 2 loci associated with ACD-AD status, including the known APOE region, and the IQUB gene on chr7 (**Supplementary File SF1 [Table ST11]**), which is not significant in AD GWAS and only suggestively significant in our ACD GWAS. No genome-wide significant loci were associated with VaD-AD status, although we observed some suggestive association (**Supplementary File SF1 [Table ST12]**).

### Protein-Protein interaction (PPI) evidenced SEMA4D, RBFOX1, and SPRY2 as hub genes for ACD and VaD

To determine the functional interactome of genes near genome-wide significant (excluding *APOE* region) and suggestive loci (P<1×10^−6^) associated with ACD and VaD, we performed a PPI analysis using the STRING database. The analysis comprised 82 ACD and 21 VaD genome-wide suggestive genes that successfully mapped to the human genome. Evidence of interaction between proteins was based on “experiments”, “co-occurrence”, “database”, and “co-expression” with a minimum score of 0.15. Non-connected proteins were removed from the network. To further determine how suggestive genes will fit in the network of known AD genes, we used *kmeans* to cluster the proteins based on validated interaction. ACD genes formed two main clusters (**Supplementary File SF2 [Figure 27])**. The first cluster was enriched in known AD genes, including *BIN1, CLU, ABCA7, CR1*, but also suggestive genes including *SEMA4D, CHD18*, and *APH1B* with more than two evidences of connection. *RBFOX1* appears to be a major hub gene for the second cluster which includes other suggestive genes like *AJAP1, ANO3*, and *TRIB1. RBFOX1* and *SEMA4D* strongly (> 2 evidences of connection) interact with known AD genes, suggesting their potential role in ACD. PPI network of VaD (**Supplementary file SF2 [Figure 28])** genes highlights the potential role *SPRY2* as it functionally connects other genes including *ERBB4, RASGRP1*, and *FOXA2*.

### Pathways and Functional Enrichment Analyses

We conducted several analyses (pathways, gene ontology, disease enrichment) to gain functional and biological contexts of genes (near variants with P< 1e-5, excluding the APOE region) associated with ACD and VaD.

#### *Pathways analysis* (Supplementary File SF1 [Table ST13])

revealed enrichment in several pathways including, “SREBF and miR33 in cholesterol and lipid homeostasis”, “Hypertrophy model”, and “Cholesterol metabolism with Bloch and Kandutsch-Russell pathways” for ACD.

#### *Gene Ontology (GO) analysis* (Supplementary Files SF2 [Figure 29], Supplementary File SF1 [Table ST14])

focusing on the biological processes (GO-BP) were enriched in terms related to amyloid-beta, “amyloid-beta metabolic process”, “amyloid precursor protein catabolic process”, and “negative regulation of amyloid precursor protein catabolic process” for ACD. For VaD, GO-BP analysis was enriched in several terms, including “response to glucose”, “response to hexose”, “response to monosaccharide”, “mesenchymal cell differentiation”, and “response to carbohydrate” (**Supplementary Files SF2 [Figure 30], Supplementary File SF1 [Table ST15]**).

#### *Disease enrichment analysis* (Supplementary Files SF2 [Figure 31], Supplementary File SF1 [Table ST16])

revealed that ACD genes were previously connected to Alzheimer’s disease, tauopathy, nephritis, central nervous system disease. It also highlighted previous associations of *SEMA4D* and *RBFOX1* with diseases of the central nervous system. Besides the AD connection, VaD genes were previously related to cancer, diabetes, and colorectal carcinoma (**Supplementary Files SF2 [Figure 32], Supplementary File SF1, [Table ST17]**) Finally, we used Framingham Heart Study data to estimate the **heritability of VaD**. We found the heritability of VaD to be 6.1%, with 95% CI of [3.2%, 21%]

## Discussion

We conducted, to our knowledge, the largest genome-wide association analyses to date, investigating genetic variants associated with ACD/ and VaD, using data from 21 cohorts and consortia with a total of 809,299 individuals across 4 ancestries, including 46,902 and 8,702 cases of ACD and VaD, respectively. Additionally, we conducted downstream analyses to understand underlying biological implications. Our findings expand current knowledge by focusing on ACD and VaD, in line with the current acknowledgment that vascular pathologies in dementia are common and have a prominent role in the pathogenesis and clinical manifestations of dementia, although controversy remains about the additive versus synergistic interactions of neurodegenerative and vascular pathologies. Our GWAS of ACD replicated several genes previously associated with AD, and GWAS of VaD identified SNPs in the ApoE region. Using functional protein-protein interaction and transcriptome-wide analyses, we identified novel genes underlying ACD that have been previously implicated in recovery from vascular injury and in neurotrophin signaling. Using genetic risk scores as instrumental variables we suggest that certain vascular risk factors may have a causal role not in both ACD and VaD pathogenesis.

In our ACD analysis of European ancestry we identified 11 genome-wide significant loci, including *BIN1, MS4A6A, PICALM, CR1, CD2AP, ABCA7, PILRB, SLC24A4*, and ’

*ACE*, all of which have been linked with AD risk in prior studies.^37^ In addition, our analyses highlighted 24 suggestive risk loci, of which 13 are novel. Among them are variants located near *ANO3*, a gene that encodes anoctamin-3, a transmembrane protein that belongs to a family of calcium-activated chloride channels and is implicated in focal dystonia, particularly craniocervical.^38^ Another suggestive locus was located near *SEMA4D*, a gene that encodes Semaphorin 4D and is known to modulate a variety of processes related to neuroinflammation and neurodegeneration, including the initiation of inflammatory microglial activation.^39,40^ Indeed, SEMA4D plays a critical role in regulating the transition between homeostatic and reactive states of various types of glial cells. Antibody blockade of SEMA4D is being explored as a potential disease modifying strategy to slow cognitive decline in patients with early Huntington’s disease^41^ and may be beneficial in other ACD. We have also identified a prominent signal near *RBFOX1*, a gene that encodes the RNA binding fox-1 which has been shown to have a role in alternative splicing of the amyloid precursor protein.^42^ Genetic variation in this gene has been associated with brain amyloid burden in preclinical and early AD^43^ and with risk of clinical AD in African-Americans.^44^ The gene may also impact dementia risk through non-amyloidogenic pathways as it additionally regulates neuron development, neuronal excitability including BDNF dependent long-term potentiation in the hippocampus,^45^ and has been implicated in brain development, essential tremor^46^ and schizophrenia.^47-51^ Other suggestive loci are located in the *ZNF652* gene, a transcriptional repressor involved in nucleic acid binding, that has diverse effects including determining the risk of hypertension.^52^ Hypertension is the most important risk factor for stroke and white matter hyperintensity and may be the most important modifiable risk factor for population prevention of dementia.^53-57^ We additionally identified a variant near *HBEGF*, a growth factor implicated in the pathobiology of CADASIL (Cerebral Autosomal Dominant Arteriopathy with Sub-cortical Infarcts and Leukoencephalopathy),^58^ the major Mendelian prototype of vascular dementia. HBEGF also has an effect on angiogenesis, expression of VEGF-A,^59^ on inflammation and oxidative stress and has been implicated in hydrocephalus.^60^

*APOE* was strongly associated both with ACD and VaD in our meta-analysis. While the association of *APOE* with ACD could be driven by AD, the relationship with VaD is less established but has been demonstrated in some population studies^61^ and candidate-gene analyses ^62,63^ and in a recent GWAS among the GR@ACE project participants.^26^ The link of *APOE* with VaD is in line with recent literature suggesting that the pathogenesis of ApoE extends beyond amyloid-β peptide aggregation and clearance. Indeed, *APOE* also influences tau-induced neurodegeneration and atrophy, microglia and the blood brain barrier,^64^ and is associated with intracranial atherosclerosis^65^, white matter hyperintensity burden, presence of cerebral microbleeds^66^ as well as with cerebral hypertensive angiopathy, which is common in individuals with VaD.^67^

In addition to a significant association of VaD with ApoE in our sample, we identified several suggestive variants also associated with VaD. These include variants near the *SPPRY2* protein-coding gene as well as *GALNT8, FOXA1, ERBB4, PSMA3* and *SEMA6D* with consistency across samples in the direction of effect, and a large number of SNPs in LD with the lead SNP. Our downstream analyses added support for a highly plausible causal link between variants including *SEMA4D, HBEGF, PIKFYVE*, and *RBFOX1 with ACD, and SPRY2* with VaD. These genes collectively emphasize a possible role for novel pathological mechanisms in ACD and VaD. A key mechanism underlying ACD highlighted by our findings is recovery after vascular injury. For example, *SEMA4D*, a member of the semaphorin family, is upregulated in the neurovascular unit after ischemic stroke, where it exerts multiple neuroprotective effects.^68,69^ Moreover, this gene has been additionally highlighted in our protein-protein interaction analysis as strongly associated with known AD genes. Another example is *SPRY2*, which was highlighted in our analyses as a suggestive gene for VaD and had strong functional associations with known ADRD genes. In the replication analysis of VaD signals in the EADB dataset, *SPRY2* has the lowest p-value and a consistent direction of association. This gene has also been suggested as a possible pharmacological target for stroke patients^70^, as it promotes angiogenesis and glial scarring around the ischemic injury, and therefore prevents increase in lesion size and secondary damage to brain tissue.^71^ In addition, *SPRY2* may exert neuroprotective effects as its expression regulates BDNF-induced signaling pathways ^72^. Similarly, *PIKFYVE* is an important regulator of platelet lysosome homeostasis ^73^, which in turn may promote recovery after ischemic stroke ^74^. Another hub gene in our analyses is *RBFOX1*, which, in addition to having a role in amyloid accumulation as discussed earlier, mediates ischemic damage by enhancing neuronal survival and Blood Brain Barrier (BBB) integrity after stroke ^75^. This gene is a neuron-specific splicing factor that regulates multiple neuronal splicing networks in neurons, and is implicated in intellectual disability, epilepsy, autism and Parkinson’s disease. Its downregulation has been associated with destabilization of mRNAs encoding for synaptic transmission proteins, which may contribute to the loss of synaptic function in Alzheimer’s disease.^76^ Furthermore, *RBFOX1* upregulation was shown to influence neuronal expression levels of the BDNF receptor, TrkB^45^, which in turn, may influence risk for ACD.^77^

We found that the MS4A gene cluster drives genetic pleiotropy that involves vascular risk factors, inflammation, hemostasis and soluble TREM2 levels. These findings are in line with preclinical studies^78-80^ and emphasize the important role and multifactorial contribution of this gene cluster to ACD pathogenesis. Although previous literature points to an association of *ACE* with AD^81^ but not VaD^82,83^, we herein show that this gene underlies both ACD and vascular risk factors. A recent study supports this finding by showing that overexpression of *ACE* on macrophages lead to reduction in vascular amyloid and GFAP+ astroglial reactivation, which indicate its role in protection of the neurovascular unit^84^. Moreover, our pairwise analysis highlighted a locus at the *PRDM6* that explained shared genetic susceptibility of VaD with ischemic stroke (IS) and WMH. Low levels of leukocyte DNA methylation of the *PRDM6* gene have been previously associated with an increased risk of ischemic stroke and with worse outcomes at 3 months after an ischemic stroke^85^. Moreover, *PRDM6* acts as an epigenetic regulator of vascular smooth muscle cell plasticity^86^.

Despite some evidence showing an inverse relationship between plasma HDL levels and risk of incident AD^87^, results are conflicting, with some studies pointing to a higher dementia risk in individuals with high HDL levels^88^, as was also the case in our study. It should be acknowledged that HDL represents a class of lipoproteins that is heterogeneous in structure and function which is not reflected by a simple measurement of HDL plasma levels^89^. High HDL levels can be deleterious under certain conditions^90^. Vascular risk factors and the presence of cardiovascular disease can alter HDL functionality by changing the structure of HDLs and converting them into proinflammatory, pro-oxidant, prothrombotic, and proapoptotic compounds^89^. We see an association with *ABCG8* in our MTAG analyses of ACD and coronary artery disease (see results). *ABCG8* is required for normal sterol homeostasis, forming an obligate heterodimer that mediates Mg(2+)- and ATP-dependent sterol transport across the cell membrane. It plays an essential role in the selective transport of dietary cholesterol in and out of the enterocytes and in selective sterol excretion by the liver into bile, which is required for normal sterol homeostasis.^91-94^ Our observation emphasizes the need for a formal causal analysis using genetic instruments to rule out potential pleiotropy and establish the causal relation between dyslipidemia and dementia risk.

Our trans-ancestral analysis identified several additional novel genes for ACD (*CHD6, DAW1* and *PWRN2*) and for VaD (*MARCHF10*), mostly driven by associations in persons of Asian ancestry. These loci were nominally significant in European Ancestry meta-analysis but have conflicting direction of association across the EA cohorts. Due to limited statistical power in analyses among Africans, Asians and Hispanics, future studies are warranted to identify the genetic determinants of ACD and VaD is non-Europeans. Nevertheless, the genes identified in our trans-ancestral analyses are biologically plausible. CDH6 has been identified as related to AD in both an epigenome-wide analysis^95^ and in an agnostic plasma proteomic analysis.^96^ It plays a role in both hippocampal synaptic development and injury.^97^ Overexpression of PWRN2, a Prader-Willi region non-protein-coding RNA 2, has been associated with in age-related macular degeneration.^98^ MARCHF10 is a E3 ubiquitin-protein ligase, a class of molecules implicated in memory formation, recovery from stroke and several neurological diseases.^99^

We replicated our VaD signals using EADB data (**Table 3**). Overall, we replicated an association within the *APOE* region. The suggestive variant near *SPRY2* also has the lowest p-value in the EADB GWAS with the same direction of effect, suggesting it could become genome-wide significant with an increased sample size.

The following caveats should be considered when interpreting the results from this study. First, the multifactorial nature and heterogeneous clinical manifestations of ACD and VaD have led to various attempts to develop diagnostic criteria^100^ which were differentially applied across the participating cohorts. More specifically, it should be noted that ACD has been ascertained using the Diagnostic and Statistical Manual of Mental Disorders-Fourth Edition DSM-IV in some studies, while others have used ICD-9/10 codes alone or in combination with autopsy or death certificate information and this can result in a varying proportion of persons identified as having dementia.^101^ Different diagnostic criteria were also used by the various cohorts to define VaD. In all cohorts a key requirement for VaD diagnosis remains the demonstration of a cognitive deficit and the presence of cerebrovascular disease, consistent with the most recent consensus criteria for VCID.^102^ Whereas these criteria differ in sensitivity and specificity, thus introducing statistical noise, this heterogeneity does not diminish the importance of loci identified despite the constraints. A second limitation is the limited power to identify associations with VaD in ancestries other than European.

Our study identified several putative genetic variants and biological pathways associated with ACD and VaD, and added additional support for the involvement of vascular mechanisms in dementia pathogenesis emphasizing a role for larger studies that embrace the entire phenotypic spectrum of ACD and VaD with further increases in sample size, especially amongst historically underrepresented populations and using emerging computational tools in order to unravel the complexity of ADRD biology.

## Methods

### 1 Study population, genotype and phenotype definitions

#### 1 – 1 Study population

A total of 809,229 participants from 21 cohorts and consortia contributed for 46,902 and 8,702 cases of ACD and VaD, respectively. The overall sample included individuals from 4 different ethnicities (European, African, Asian, Hispanic) from North America, Europe, and Asia. The mean age ranged between 54 and 80 years with 54% to 68% females. The summary demographics is described in **Table 1 (**also detailed in **Supplementary file SF1 [Table ST1])**. Each study obtained written informed consent from participants or, for those with substantial cognitive impairment, from a caregiver, legal guardian, or other proxies. Study protocols for all cohorts were reviewed and approved by the appropriate institutional review boards.

#### 1 – 2 Phenotypes definition

The primary study outcomes are VaD and ACD, measured by each participating cohort as described in **Supplementary File SF3 (Supplementary File SF1 [Table ST18])**. Moreover, to increase the sensitivity, cohorts were asked to run separate association analyses for (a) incident all-cause dementia, (b) prevalent all-cause dementia, (c) incident vascular dementia, (d) prevalent vascular dementia, (e) incident probable and definite vascular dementia, and (f) prevalent probable whenever possible. To address the overlap with AD, we included all VCID (including persons with possible VCID) and separately analyze only cases of ‘pure’ (probable and autopsy proven definite) VCID and requested that each cohort provides the most accurate, detailed description of their diagnostic algorithm. Although VaD in UKB was defined based on ICD-10 codes, we used the family history of dementia GWAS (“*imputed dementia*”) recently published by Marioni et al.^30^ for ACD. Imputed dementia was defined as individuals >= 65 years reporting a history of dementia in one or both parents. As explained in Ghosh et al. ^103^ the effect sizes and standard errors of the imputed dementia GWAS were doubled 2 to analytically correct for the use of proxy phenotypes.

#### 1 – 3 Genotyping and imputation

Genotyping was performed using cohort-specific genotyping arrays as described in **Supplementary File SF3**. Genetic variants were imputed using 1000 Genomes Project (1KG), the Haplotype Reference Consortium (HRC) ^104^, and the NHLBI Trans-Omics for Precision Medicine (TOPMed). UK biobank imputed the genotypes to HRC, 1KG, and UK10K. Details on study-specific quality control filters and software used for phasing and imputation are provided in Supplementary materials. Briefly, rare variants (MAF <1%) and poorly imputed variants (Imputation quality, Rsq < 0.3) were excluded as well as variants mapping to sex chromosomes or mitochondria. Samples with poor genotyping call rate (<95%) and Hardy-Weinberg P-values < 1×10^−6^ were removed. All genetic positions are reported in genome build 37 (GRCh37, hg19). Moreover, we use HRC v.1.2 as the main reference panel and selected only variants in this panel were subsequently used in the association analyses. Additional details on the genotyping and imputation methods and quality control are provided in **Supplementary File SF3**.

### 2 Genome-wide association analyses, Quality control, and meta-analysis

#### 2 – 1 Study-level association analyses

We conducted study and ethnic-specific association analyses adjusting for age, sex, sites, and population structure to test the association of each variant with VaD and ACD. Cohorts were asked to run logistic regression and Cox proportional hazard models for prevalent and incident VaD/ACD respectively assuming additive allelic effects and imputed dosages. The UKB association analyses were performed with linear mixed models (LMM) using the BOLT-LMM software.^105^ BOLT-LMM has the advantage over other methods in that it accounts for cryptic relatedness and population structure and thus, allows the inclusion of related individuals in the models which increase the overall sample size. Details on the methods and software used for study-level association analyses are provided in **Supplementary File SF3**.

#### 2 – 2 Quality control of study-level summary statistics

We performed stringent quality-control (QC) check of the summary statistics from each cohort using EasyQC.^106^ We mapped each variant from the non-EA cohort to the appropriate 1KG project phase 3 reference panel and all European Ancestry to HRC (details in **Supplementary File SF1 [Table ST1]**). Then, the following steps were performed to ensure proper QC of each file before meta-analyze: (**a**) remove all structural variants and INDELs; (**b**) filter out variants with missing or unusual values (P-value < 0 or > 1, effect size > 10, effect allele frequency < 0 or > 1, imputation quality <0 or > 1); (**c**) filter out variants with effective allele count (EAC, 2 x minor allele frequency x N x imputation quality) < 10; (d) filter out variants with low imputation quality (e.g., INFO scores reported by the imputation software); (e) filter out variant with MAF < 5%; (f) align variants to the main reference panel (HRC for EA, and ethnic-specific 1KG for others); remove variants with absolute difference between its allele frequencies in the cohort and reference panel greater than 0.2. All variants were assigned a unique identifier as a combination of chromosome, position, reference, alternative alleles separated by semi-colon (CHR:POS:REF:ALT) to avoid issue with chromosomal positions mapping to multiple markers IDs. The above steps were repeated until satisfying results were obtained after visual inspection of the different diagnostic QC plots (AF, P-Z, Q-Q, and SE plots) generated by EasyQC as explained in Winkler et al.^106^.

### 3 Meta-analysis of GWAS results

#### 3 – 1 Ancestry-specific meta-analysis

The meta-analyses were conducted using the fixed-effect sample size weighted method implemented in METAL.^107^ Post-analysis results were filtered to retain only variants present in more than 40% of the overall cohorts and the effective sample size greater than 40% of the study sample size. We evaluated the heterogeneity across cohorts using the I^2^ statistic provided by METAL, which represents the percentage of variation across studies that is due to heterogeneity rather than chance. Variants with heterogeneity I^2^ statistic > 95% were filtered out. We used the standard p-value thresholds for genome-wide significance, p-value <=5×10^−8^; and suggestive, p-value <= 1×10^−6^. Since there was no evidence of genomic inflation in cohort’s summary statistics (lambda 0.98 – 1.06), no genomic control was applied during the meta-analysis. Genomic loci were defined as the region +-500kb around the SNP with the lowest p-value, considered as the index-SNP. We assessed the heterogeneity across studies using the I^2^ statistics of METAL (HetPVal output), which represents the percentage of variation across cohorts that is due to genetic heterogeneity rather than chance. Except for the APOE region (Chr 19, position 45xxxx) for which the HetPVal was > 1e-8, significant SNPs were selected with HetPVal > 0.01). We conducted ancestry-specific meta-analyses of VaD and ACD for European Ancestry (EA), African Ancestry (AA), Asian Ancestry (SA), and Hispanic ancestry (HA). In addition, we used LD score regression to quantify the contribution of true polygenicity and biases such as cryptic relatedness and population stratification of the meta-analysis results.

#### 3 – 2 Trans-ancestry meta-analysis

We performed trans-ancestry meta-analyses to assess whether the increase in sample size could lead to adequate power to identify additional genome-wide significant loci associated with ACD and VaD. To this end, we used MR-MEGA (Meta-Regression of Multi-Ethnic Genetic Association) software, which has been proven to be more efficient than others when dealing with genetic heterogeneity^108^. MR-MEGA uses a matrix of mean pairwise allele frequency differences to quantify genetic similarity between studies, and estimate the effect of each SNP after adjusting for ancestry principal components. We applied study-specific filters, as previously described in the QC section, with Effective Allele Count (EAC) > 20, for studies with small sample size, to reduce the amount of noise in the results driven rare SNPs in small cohorts. We fitted three principal components, as suggested by MR-MEGA authors, which proved sufficient to separate the cohorts into self-reported ancestry groups (**Supplementary File SF2 [Figure 33]**). As in the ancestry-specific meta-analysis, we retained only SNPs that were present in > 40% cohorts, with > 40% total sample size. Genome-wide significant SNPs had p-value < 5e^-8^ and showed evidence. Of allelic heterogeneity across population (MR-MEGA P-Het > 1e^-5^).

### 4 Shared genetic susceptibility with complex disease traits

Gene-based association test was conducted using MAGMA,^109^ with P<2.8×10−6 as a gene-wide significance threshold. Gene regions with SNPs not reaching GW significance for ACD or VaD in the primary GWAS analysis and additionally not in LD (r2<0.10) with the lead SNP were considered as novel.

We first explored the association of lead risk variants with related vascular, neurological traits and metabolic traits, excluding the APOE region. For each related trait, association statistics of SNPs falling in a window of ±250kb around each lead SNP were queried^110^ and SNPs satisfying the GW significance threshold in the original study were retained. Leveraging the polygenicity of ACD (mean chi2 = 1.1) and VaD (mean chi2 =1.06), we systematically explored the genetic overlap of ACD and VaD (in the European-only analysis) with (i) neurological and neurodegenerative traits (any stroke-AS, ischemic stroke-IS, small vessel stroke-SVS, large artery stroke-LAS, cardioembolic stroke-CES; general cognitive function-GCF and Alzheimer type dementia-AD); (ii) common MRI-imaging marker of cerebral small vessel disease (white matter hyperintensities-WMH)^9^ and (iii) vascular risk factors [systolic, diastolic and pulse pressure (SBP, DBP, PP); high-density, low-density lipoprotein (HDL, LDL)].^111^ We acquired summary statistics of the largest European-only GWAS for these traits.

Using LDSR^112^, genetic correlation estimates between ACD/VaD and the aforementioned complex traits were obtained. A p-value < 8.3×10^−3^ correcting for 6 independent phenotypes was considered significant. In the absence of confounding and sample overlap, the genetic covariance intercept is close to zero. As genome-wide correlation estimates may miss significant correlations at the regional level (balancing-effect),^113^ a Bayesian pairwise GWAS approach (GWAS-PW) was applied.^114^ GWAS-PW identifies trait pairs with high posterior probability of association (PPA) with a shared genetic variant (model 3, PPA3≥ 0.90). To ensure that PPA3 is unbiased by sample overlap, fgwas v.0.3.6 was run on each pair of traits and the correlation estimated from regions with null association evidence (PPA<0.20) was used as a correction factor.^114^ We then calculated Spearman’s rank correlation for regions showing PPA3>0.90, approximating the direction of effect.

Finally, using a Bayesian method - *ashR*^115^, we studied the effect-size distribution for ACD and VaD and related risk factors. Briefly, *ashr* tests the probability of non-zero effect conferred by SNPs as a function of linkage disequilibrium (LD) score, measuring the true effect size that is not zero and the underlying polygenic background. Using MTAG^116^, traits falling in similar polygenic profile to ACD or VAD are jointly analyzed in a bivariate scheme leveraging the pairwise trait genetic correlation to boost power to discover new loci. The significance threshold in the MTAG analysis is determined based on the number of traits sharing a similar polygenic profile, additionally was restricted to SNPs that also had nominal significance (p<0.05) for each phenotype separately in the pre-existing univariate GWAS.

### Transcriptome-wide association study and colocalization

We performed transcriptome-wide association studies (TWAS) using the association statistics from the ACD and VaD (European-only) and weights from 22 publicly available gene expression reference panels from blood (Netherlands Twin Registry, NTR; Young Finns Study, YFS), arterial (Genotype-Tissue Expression, GTEx), brain (GTEx, CommonMind Consortium, CMC) and peripheral nerve tissues (GTEx). For each gene in the reference panel, precomputed SNP-expression weights in the 1-Mb window were obtained, including the highly-tissue specific splicing QTL (sQTL) information on gene isoforms in the dorsolateral prefrontal cortex (DLPFC) derived from the CMC. TWAS-Fusion^117^ was used to estimate the TWAS Z score (association statistic between predicted expression and ACD or VaD), derived from the SNP-expression weights, SNP-trait effect estimates and the SNP correlation matrix. Transcriptome-wide (TW) significant genes (eGenes) and the corresponding QTLs (eQTLs) were determined using Bonferroni correction in each reference panel, based on the average number of features (4,235 genes) tested across all the reference panels^117^. eGene regions with eQTLs not reaching genome-wide significance in association with ACD or VaD, and not in LD (r^2^<0.01) with the lead SNP for genome-wide significant risk loci, were considered as novel. Finally, a colocalization analysis (COLOC)^118^ was carried out at each locus to estimate the posterior probability of a shared causal variant (PP4≥0.75) between the gene expression and trait association, using a prior probability of 1.1×10^−5^. Furthermore, functional validation of the eGenes was performed by testing for positional overlap of the best eQTLs from TWAS with enhancer (H3K4me1, H3K27ac) and/or promoter (H3K4me3/H3K9ac) elements across a broad category of relevant tissue types (blood-BLD, brain/neurological-BRN) using Haploreg V4.1^119^.

### Identification of independent case-case loci with CC-GWAS

Leveraging summary statistics from our GWAS of ACD and VaD, as well as from publicly available existing GWAS of AD^120^, we examined genetic uniqueness between these highly correlated whereas distinct disorders using case-case GWAS, a method that test for differences in allele frequency between cases of two disorders without individual-level data.^121^ By allowing for sample overlap between the two case-control GWAS, case-case GWAS has the potential to increase power to detect signals otherwise missed in case-control GWAS.

## Data availability

AD GWAS Summary data for CC-GWAS: https://ctg.cncr.nl/software/summary_statistics

## Code availability

CC-GWAS: https://github.com/wouterpeyrot/CCGWAS

## Definition

GWAS: Genome-Wide Association Analysis Study
ACD: All-cause dementia
VaD: Vascular dementia
VCID: Vascular Cognitive Impairment and Dementia
CHARGE: Cohorts for Heart and Aging Research in Genomic Epidemiology
ADGC: Alzheimer’s Disease Genetics *Consortium*
UKBB: UK Biobank
EADB: European Alzheimer’s Disease DNA BioBank

## Notes

### Competing Interest Statement

The authors have declared no competing interest.

