## Supplementary File SF2 for "A meta-analysis of genome-wide association studies identifies new genetic loci associated with all-cause and vascular dementia"

**(Figures and tables)**

### Introduction

### 1 - Meta-analysis plots

#### 1 – 1 All-cause dementia of European ancestry (ACD)

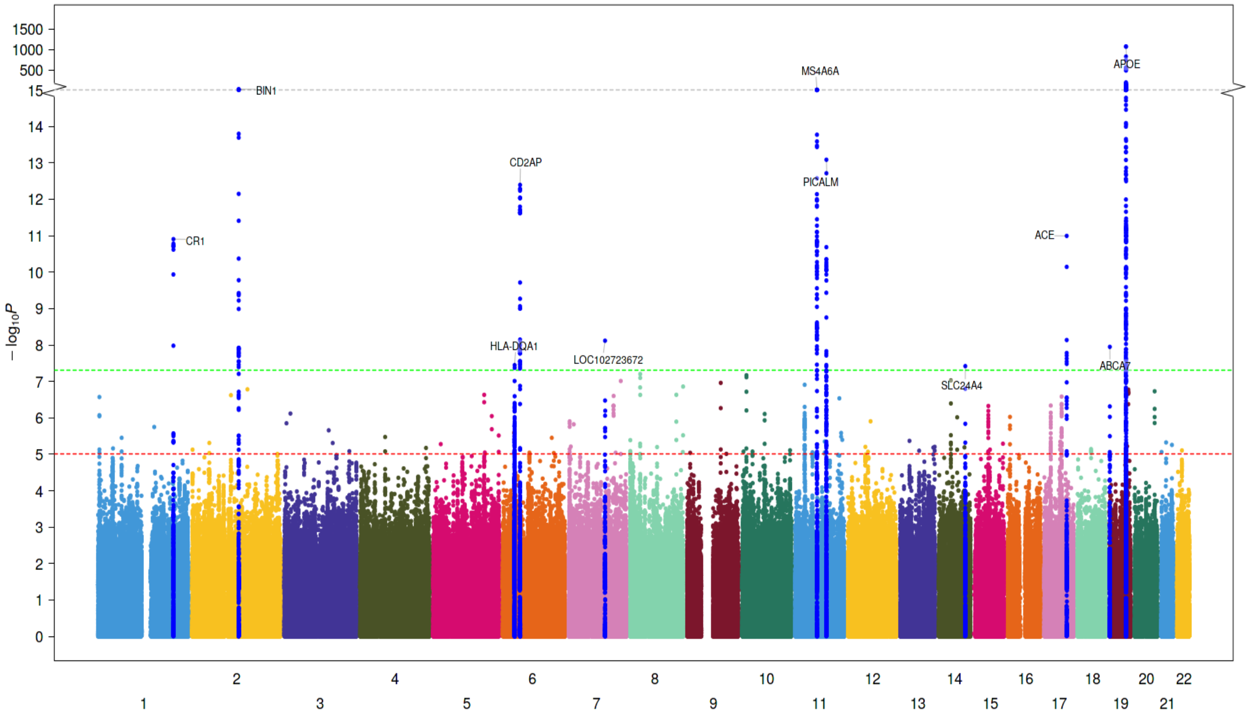

**Figure 1: Manhattan plot of the ACD GWAS.** In addition to variants in the APOE region, we identified five new genetic loci associated with VaD. Blue and red lines correspond to the p-value of 5e$^{-7}$ and 5e$^{-8}$ for genome-wide suggestive and significant SNPs, respectively. Manhattan plots for the cross-ancestry meta-analysis. Each dot represents a SNP, the X-axis shows the chromosomes where each SNP is located, and the Y-axis shows -log10 P-value of the association of each SNP with POAG in the cross-ancestry meta-analysis. The red horizontal line shows the genome-wide significant threshold (P-value=5e-8; -log10 P-value=7.30). The nearest gene to the most significant SNP in each locus has been labeled.

#### 1 – 2 Vascular dementia European ancestry (VaD)

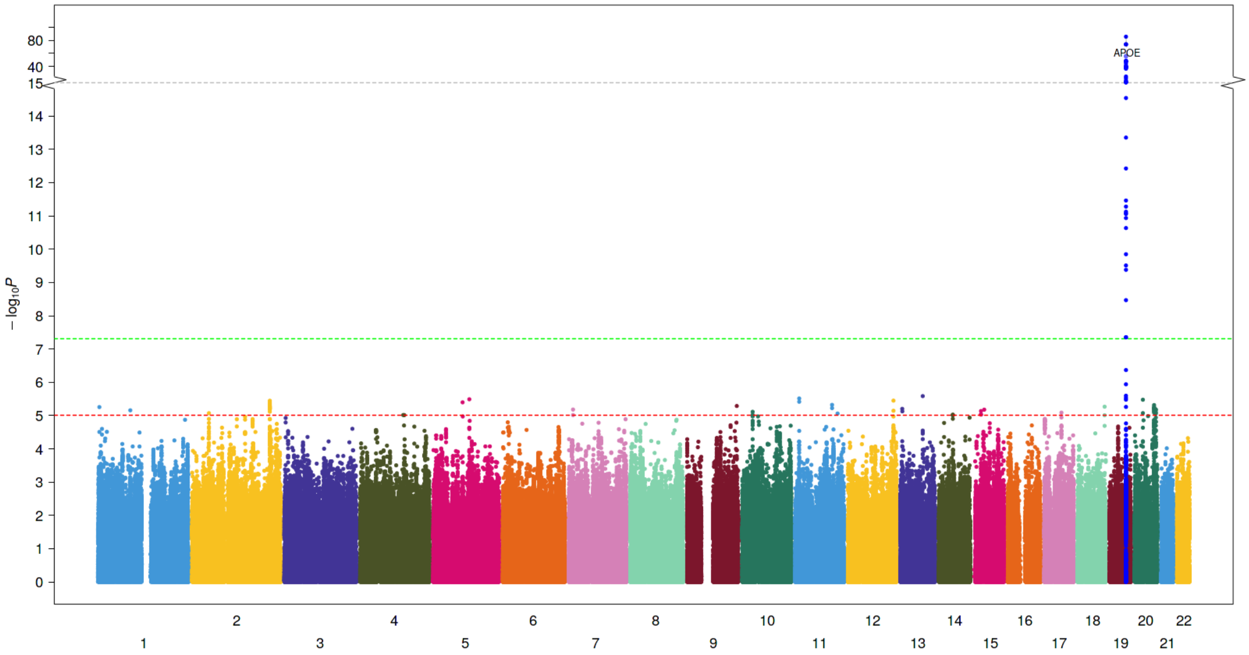

**Figure 2: Manhattan plot of the VaD GWAS.** In addition to variants in the APOE region, we identified five new genetic loci associated with VaD. Blue and red lines correspond to the p-value of 5e$^{-7}$ and 5e$^{-8}$ for genome-wide suggestive and significant SNPs, respectively. Manhattan plots for the cross-ancestry meta-analysis. Each dot represents a SNP, the X-axis shows the chromosomes where each SNP is located, and the Y-axis shows -log10 P-value of the association of each SNP with POAG in the cross-ancestry meta-analysis. The red horizontal line shows the genome-wide significant threshold (P-value=5e-8; -log10 P-value=7.30). The nearest gene to the most significant SNP in each locus has been labeled.

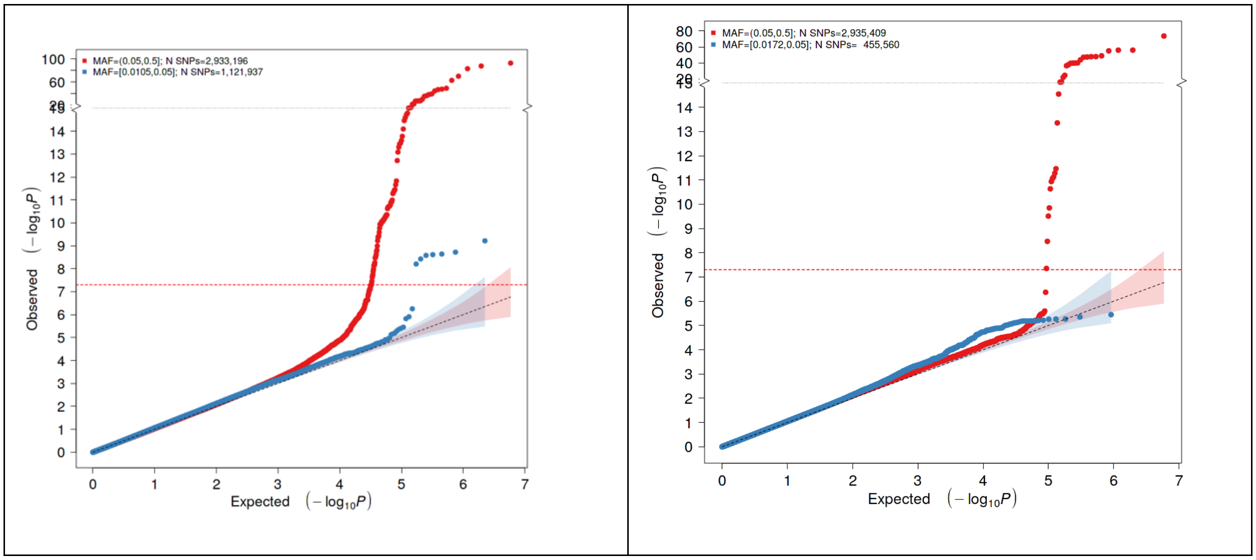

**Figure 3: Q-Q plots of the ACD (left) and VaD (right) GWASs.** The expected P values (X-axis) are plotted against the observed P-values (Y-axis). The units of the axes are the –log10 of the P-value. The red and blue curves represent the plots with MAF >= 0.05 and 0.01 respectively. The diagonal line of the null hypothesis and its 95% confidence interval were plotted in grey based on the P-values without the previously reported SNPs. The red dot line represents the cutoff for genome-wide significance.

#### 1 – 3 Trans-ancestral meta-analysis of all-cause dementia (ACD) GWASs

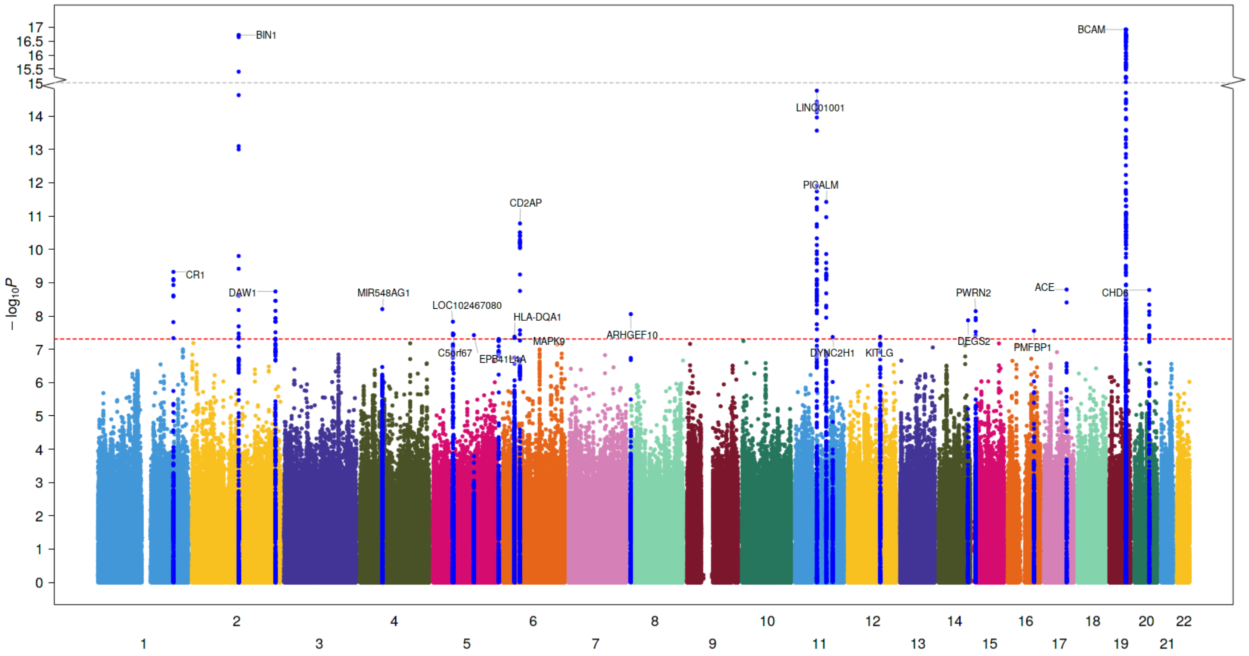

**<u>Figure 4:</u> Manhattan plot of the trans-ancestral meta-analysis of ACD GWASs.** In addition to variants in the APOE region, we identified five new genetic loci associated with VaD. Blue and red lines correspond to the p-value of 5e$^{-7}$ and 5e$^{-8}$ for genome-wide suggestive and significant SNPs, respectively. Manhattan plots for the cross-ancestry meta-analysis. Each dot represents a SNP, the X-axis shows the chromosomes where each SNP is located, and the Y-axis shows -log10 P-value of the association of each SNP with POAG in the cross-ancestry meta-analysis. The red horizontal line shows the genome-wide significant threshold (P-value=5e-8; -log10 P-value=7.30). The nearest gene to the most significant SNP in each locus has been labeled.

#### 1 – 4 Trans-ancestral meta-analysis of vascular dementia (VaD) GWASs

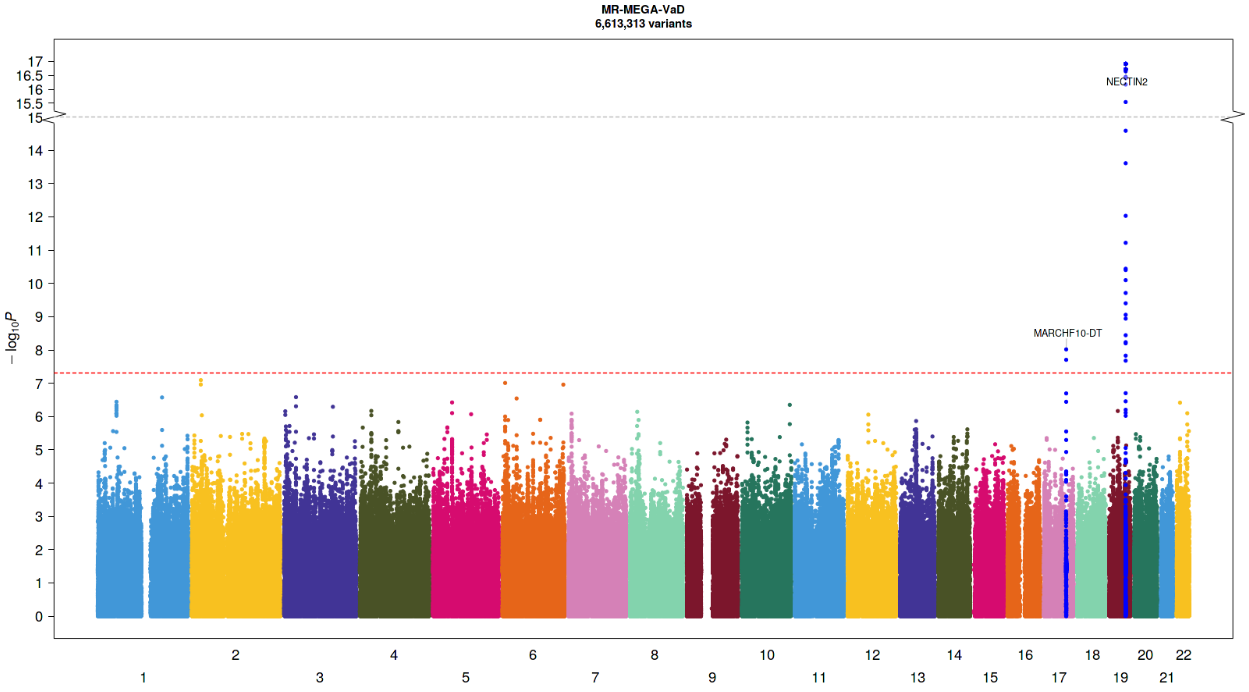

**Figure 5:** **Manhattan plot of the trans-ancestral meta-analysis of VaD GWASs.** In addition to variants in the APOE region, we identified five new genetic loci associated with VaD. Blue and red lines correspond to the p-value of $5e^{-7}$ and $5e^{-8}$ for genome-wide suggestive and significant SNPs, respectively. Manhattan plots for the cross-ancestry meta-analysis. Each dot represents a SNP, the X-axis shows the chromosomes where each SNP is located, and the Y-axis shows -log10 P-value of the association of each SNP with POAG in the cross-ancestry meta-analysis. The red horizontal line shows the genome-wide significant threshold (P-value=5e-8; -log10 P-value=7.30). The nearest gene to the most significant SNP in each locus has been labeled.

### 2 - Meta-analysis tables

#### 2 – 1 Meta-analysis of ACD GWASs in European population

**Table 1.** Genome-wide significant ($P<5x10^{-8}$) and suggestive ($P<1x10^{-6}$) variants associated with all-cause dementia in European population.

| rsID | Nearest Gene | Chr | Pos | EA/NEA | Pval | MAF | Beta |
|---|---|---|---|---|---|---|---|
| rs429358 | APOE | 19q13.32 | 45411941 | T/C | 1.E-1180 | 0.8485 | -1.17 |
| rs4663105 | BIN1 | 2q14.3 | 127891427 | A/C | 5.8E-34 | 0.4036 | -0.15 |
| rs12453 | MS4A6A | 11q12.2 | 59945745 | T/C | 1.1E-16 | 0.4046 | 0.09 |
| rs10792832 | PICALM | 11q14.2 | 85867875 | A/G | 8.2E-14 | 0.3718 | -0.08 |
| rs10948367 | CD2AP | 6p12.3 | 47585615 | A/G | 4.0E-13 | 0.2555 | -0.08 |
| rs4295 | ACE | 17q23.3 | 61556298 | C/G | 1.0E-11 | 0.3718 | -0.07 |
| rs4844610 | CR1 | 1q32.2 | 207802552 | A/C | 1.2E-11 | 0.172 | 0.09 |
| rs2906644 | PILRB | 7q22.1 | 99956290 | C/G | 7.6E-09 | 0.1113 | 0.10 |
| rs3764650 | ABCA7 | 19p13.3 | 1046520 | T/G | 1.1E-08 | 0.1054 | -0.11 |
| rs9323877 | SLC24A4 | 14q32.12 | 92934269 | A/G | 3.8E-08 | 0.2296 | -0.07 |
| rs1532278 | CLU | 8p21.1 | 27466315 | T/C | 6.2E-08 | 0.3857 | -0.06 |
| rs7912495 | USP6NL | 10p14 | 11718713 | A/G | 6.7E-08 | 0.4543 | -0.06 |
| rs17125924 | FERMT2 | 14q22.1 | 53391680 | A/G | 9.3E-08 | 0.07952 | -0.10 |
| rs11767557 | EPHA1 | 7q34 | 143109139 | T/C | 9.7E-08 | 0.2177 | 0.07 |
| rs1854554 | SEMA4D | 9q22.2 | 92155871 | A/G | 1.0E-07 | 0.4493 | 0.06 |
| rs7118826 | ANO3 | 11p14.2 | 26195535 | C/G | 1.2E-07 | 0.4732 | -0.07 |
| rs79832570 | SPATC1 | 8q24.3 | 145097720 | T/C | 1.3E-07 | 0.09344 | -0.14 |
| rs13010870 | RBM43 | 2q23.3 | 151765163 | T/C | 1.6E-07 | 0.2485 | 0.07 |
| rs1354106 | CD33 | 19q13.41 | 51737991 | T/G | 1.6E-07 | 0.338 | 0.06 |
| rs6014724 | CASS4 | 20q13.2 | 54998544 | A/G | 1.8E-07 | 0.08946 | 0.11 |
| rs897150 | TRIB1 | 8q24.13 | 126576702 | A/G | 2.3E-07 | 0.2813 | -0.06 |
| rs11168036 | HBEGF | 5q31.3 | 139707439 | T/G | 2.3E-07 | 0.4831 | 0.06 |
| rs677649 | RNU6-11P | 7 | 123439244 | T/G | 2.5E-07 | 0.1581 | -0.08 |
| rs8081878 | ZNF652 | 17q21.32 | 47436812 | A/T | 2.6E-07 | 0.4553 | -0.06 |
| rs4654450 | RP1-37J18.2 | 1 | 4667378 | A/G | 2.6E-07 | 0.336 | -0.07 |
| rs11218343 | SORL1 | 11q24.1 | 121435587 | T/C | 2.9E-07 | 0.04274 | 0.17 |
| rs2297508 | SREBF1 | 17p11.2 | 17715317 | C/G | 4.6E-07 | 0.4076 | 0.06 |
| rs442495 | ADAM10 | 15q21.3 | 59022615 | T/C | 4.7E-07 | 0.3549 | 0.06 |
| rs13316744 | AC091493.2 | 3 | 16742711 | C/G | 7.6E-07 | 0.4543 | -0.05 |
| rs7068231 | ANK3 | 10q21.2 | 61784928 | T/G | 7.8E-07 | 0.4006 | -0.06 |
| rs834398 | GABRB2 | 5q34 | 160528276 | A/G | 8.9E-07 | 0.1899 | -0.08 |
| rs62013908 | RBFOX1 | 16p13.3 | 5991314 | C/G | 9.4E-07 | 0.2336 | 0.07 |

#### 2 – 2 Meta-analysis of VaD GWASs in European population

| rsID | Nearest gene | CHR | POS | EA/NEA | Pval inCHARGE | Pval in EADB | Combined Pval | Direction |
|---|---|---|---|---|---|---|---|---|
| rs429358 | APOE | 19q13.32 | 45411941 | T/C | 2.67E-86 | 5.66E-113 | 2.90E-196 | -- |
| rs11911 | SPRY2 | 13q31.1 | 80910851 | A/C | 2.60E-06 | 6.53E-02 | 3.35E-05 | ++ |
| rs7101996 | MTND5P21 | 11 | 11259298 | T/C | 3.06E-06 | 4.80E-01 | 4.52E-03 | -+ |
| rs2845990 | GALNT18 | 5 | 98907502 | T/C | 3.24E-06 | 8.72E-01 | 1.31E-03 | +- |
| rs117904289 | FOXA2 | 20p11.21 | 22782154 | A/G | 3.35E-06 | 9.87E-01 | 1.12E-03 | -+ |
| rs838941 | SCARB1 | 12q24.31 | 125183316 | A/G | 3.56E-06 | 3.30E-01 | 6.54E-05 | -- |
| rs17418160 | ERBB4 | 2q34 | 213119022 | T/C | 3.58E-06 | 8.82E-01 | 1.71E-03 | -+ |
| rs77542509 | TRPC6 | 11q22.1 | 101415824 | T/C | 4.77E-06 | 1.37E-01 | 2.13E-05 | ++ |
| rs6127311 | DOK5 | 20q13.2 | 53501017 | T/C | 4.83E-06 | 2.94E-01 | 1.44E-02 | -+ |
| rs35945091 | LCN1P2 | 9 | 136185411 | T/C | 5.15E-06 | 8.55E-01 | 1.76E-03 | +- |
| rs17059857 | ZNF236 | 18q23 | 74469493 | T/C | 5.41E-06 | 4.17E-01 | 9.34E-03 | -+ |
| rs143750890 | AJAP1 | 1p36.32 | 4602505 | T/C | 5.56E-06 | 1.63E-01 | 3.34E-05 | -- |
| rs9510987 | SPATA13 | 13 | 24575243 | T/G | 6.28E-06 | 2.91E-01 | 7.44E-05 | ++ |
| rs55709546 | PHACTR3 | 20q13.32 | 58261107 | A/C | 6.53E-06 | 6.45E-01 | 4.96E-04 | ++ |
| rs12667855 | TMEM106B | 7p21.3 | 12124166 | T/G | 6.59E-06 | 5.42E-01 | 2.99E-04 | ++ |
| rs281219 | SEMA6D | 15q21.1 | 47711652 | A/G | 6.65E-06 | 3.16E-01 | 1.29E-02 | -+ |
| rs138352554 | GBP1 | 1p22.2 | 89517105 | A/G | 6.95E-06 | 5.73E-01 | 4.06E-04 | ++ |
| rs16967121 | RASGRP1 | 15q14 | 38923007 | A/G | 7.31E-06 | 9.16E-01 | 1.26E-03 | ++ |
| rs11007123 | WAC | 10p12.1 | 28763005 | T/C | 7.70E-06 | 2.92E-01 | 1.39E-02 | -+ |
| rs4794009 | GIP | 17q21.32 | 47051955 | A/G | 8.17E-06 | 7.97E-01 | 7.46E-04 | ++ |
| rs35448830 | PRKCE | 2p21 | 46080762 | T/C | 8.48E-06 | 6.31E-01 | 5.79E-04 | ++ |
| rs2233754 | PSMA3 | 14q23.1 | 58755574 | A/C | 9.39E-06 | 6.01E-01 | 6.04E-03 | -+ |

**Table 2. Genome-wide significant ($P<5x10^{-8}$) and suggestive ($P<1x10^{-6}$) variants associated with vascular dementia in European population**. The meta-analysis includes 11 cohorts from the CHARGE consortium, and the UK Biobank (UKBB) GWAS. Direction denotes the direction of association in CHARGE and EADB

#### 2 – 3 Trans-ancestral meta-analysis of ACD GWASs

| rsID | Nearest Gene | Chr | Pos | EA/NEA | Pval | MAF | Beta | SE |
|---|---|---|---|---|---|---|---|---|
| rs10402524 | BCAM | 19p11 | 45329344 | T/C | 1.21E-17 | 0.2336 | -0.168 | 0.045 |
| rs744373 | BIN1 | 2q14.3 | 127894615 | A/G | 1.90E-17 | 0.358 | -0.139 | 0.031 |
| rs2278867 | MS4A6A | 11q13.1 | 59943109 | A/T | 1.72E-15 | 0.2897 | 0.113 | 0.020 |
| rs10792832 | PICALM | 11q13.1 | 85867875 | A/G | 3.77E-12 | 0.3135 | -0.074 | 0.036 |
| rs10948367 | CD2AP | 6q14.3 | 47585615 | A/G | 1.67E-11 | 0.2328 | -0.042 | 0.017 |
| rs1408077 | CR1 | 1q11.1 | 207804141 | A/C | 4.75E-10 | 0.1412 | 0.088 | 0.055 |
| rs4295 | ACE | 17q21.1 | 61556298 | C/G | 1.60E-09 | 0.3666 | -0.066 | 0.018 |
| rs2208524 | CHD6 | 20q11.21 | 40423299 | T/C | 1.66E-09 | 0.1268 | -0.103 | 0.027 |
| rs11691153 | DAW1 | 2q14.1 | 228780072 | T/C | 1.83E-09 | 0.1536 | 0.099 | 0.025 |
| rs6853262 | LPHN3 | 4q22.1 | 61221892 | C/T | 6.22E-09 | 0.06989 | 0.208 | 0.112 |
| rs2677386 | PWRN2 | 15q15.1 | 24432053 | T/C | 7.18E-09 | 0.3612 | -0.083 | 0.017 |
| rs7006786 | ARHGEF10 | 8q13.2 | 1792639 | G/A | 8.81E-09 | 0.08986 | 0.097 | 0.045 |
| rs35483531 | DEGS2 | 14q21.3 | 100653772 | C/T | 1.35E-08 | 0.2993 | -0.004 | 0.024 |

| | | | | | | | | |
|---|---|---|---|---|---|---|---|---|
| rs170084 | PMFBP1 | 16q11.2 | 72178483 | T/A | 2.79E-08 | 0.107 | -0.068 | 0.029 |
| rs10940421 | SNX18 | 5q14.3 | 54036059 | A/G | 3.34E-08 | 0.372 | 0.040 | 0.017 |
| rs138908633 | EPB41L4A | 5q14.3 | 111649017 | G/A | 3.76E-08 | 0.03095 | -0.029 | 0.050 |
| rs74435987 | DUSP6 | 12q14.1 | 89152253 | G/T | 4.20E-08 | 0.08766 | 0.078 | 0.130 |
| rs11225924 | DDI1 | 11q13.1 | 103493165 | C/T | 4.27E-08 | 0.1034 | 0.152 | 0.105 |
| rs113747850 | MAPK9 | 5q14.3 | 179710663 | T/C | 4.93E-08 | 0.123 | 0.071 | 0.024 |

**Table 3. Genome-wide significant ($P<5x10^{-8}$) and suggestive ($P<1x10^{-6}$) variants associated with all-cause dementia in trans-ancestral meta-analysis**. The meta-analysis includes European, African, Asian, and Hispanic/Latino ancestries. Three new variants at 20q11.21, 2q14.1, and 15q15.1 reached genome-wide significance.

#### 2 – 4 Trans-ancestral meta-analysis of VaD GWASs

| rsID | Nearest Gene | Chr | Pos | EA/NEA | Pval | MAF | Beta | SE |
|---|---|---|---|---|---|---|---|---|
| **rs10119** | TOMM40 | 19 | 45406673 | G/A | 1.21E-17 | 0.2476 | -0.327 | 0.054 |
| **rs4380108** | MARCHF10 | 17 | 60893485 | C/T | 9.59E-09 | 0.3127 | -0.172 | 0.031 |
| **rs55747619** | ITSN2 | 2 | 24530447 | C/G | 8.05E-08 | 0.08706 | -0.384 | 1.969 |
| **rs9379092** | CAGE1 | 6 | 7344531 | G/A | 9.80E-08 | 0.1172 | -0.336 | 0.077 |
| **rs3757193** | RPS6KA2 | 6 | 166923463 | C/T | 1.09E-07 | 0.08347 | 2.151 | 0.636 |
| **rs3871399** | CMTM7 | 3 | 32496413 | C/G | 2.61E-07 | 0.124 | 0.550 | 0.640 |
| **rs17315346** | BRINP2 | 1 | 177282235 | C/T | 2.67E-07 | 0.01538 | -2.412 | 5.17 |
| **rs1738249** | DNAH8 | 6 | 38753960 | C/T | 2.86E-07 | 0.3013 | -0.050 | 0.040 |
| **rs12095469** | OSBPL9 | 1 | 52206082 | G/A | 3.60E-07 | 0.05292 | 3.654 | 3.513 |
| **rs4820650** | ADRBK2 | 22 | 25925358 | T/C | 3.82E-07 | 0.2468 | 0.050 | 0.054 |
| **rs61859886** | MGMT | 10 | 131353192 | T/G | 4.45E-07 | 0.1528 | -0.274 | 0.057 |
| **rs9857196** | RYK | 3 | 133830660 | T/A | 5.09E-07 | 0.01997 | 3.438 | 2.670 |
| **rs637924** | PCDH7 | 4 | 31465610 | T/C | 6.77E-07 | 0.2564 | -0.056 | 0.051 |
| **rs35810115** | ZNF675 | 19 | 23780763 | C/T | 6.81E-07 | 0.04992 | -1.169 | 2.339 |
| **rs115331896** | CRBN | 3 | 3204942 | T/G | 6.95E-07 | 0.01218 | 3.139 | 2.706 |
| **rs4401880** | SLC18A1 | 8 | 19946066 | C/T | 7.20E-07 | 0.3249 | 0.019 | 0.051 |
| **rs4823298** | FBLN1 | 22 | 45915987 | T/C | 7.96E-07 | 0.4581 | 0.029 | 0.0439 |
| **rs17335455** | NXPH1 | 7 | 8853946 | T/G | 8.20E-07 | 0.1633 | -0.096 | 0.042 |
| **rs517484** | RP11-6N13.1 | 5 | 104490130 | T/C | 8.51E-07 | 0.1965 | -0.120 | 0.046 |
| **rs12814413** | RBMS2 | 12 | 56916614 | T/C | 8.77E-07 | 0.3514 | 0.050 | 0.051 |
| **rs4665372** | CGREF1 | 2 | 27325837 | T/A | 9.19E-07 | 0.3948 | -0.104 | 0.043 |

**Table 4. Genome-wide significant ($P<5x10^{-8}$) and suggestive ($P<1x10^{-6}$) variants associated with vascular dementia in trans-ancestral meta-analysis**. The meta-analysis includes European, African, Asian, and Hispanic/Latino ancestries. One new variant at

### 3 – Variant plots

#### 3 – 1 All-cause dementia suggestive variants

#### 3 – 1 – 1 ANO3 locus

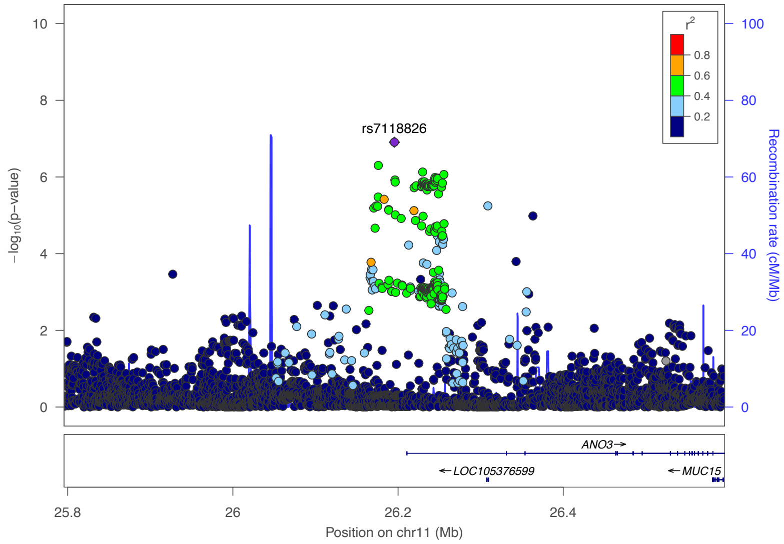

**Figure 7: Regional association plot showing the genomic region containing ANO3.** For each SNP, the P-value (log10 scale) of the association with ACD is represented (y-axis, left). The recombination rates (y-axis right), which reflects the local linkage disequilibrium structure is also plotted.

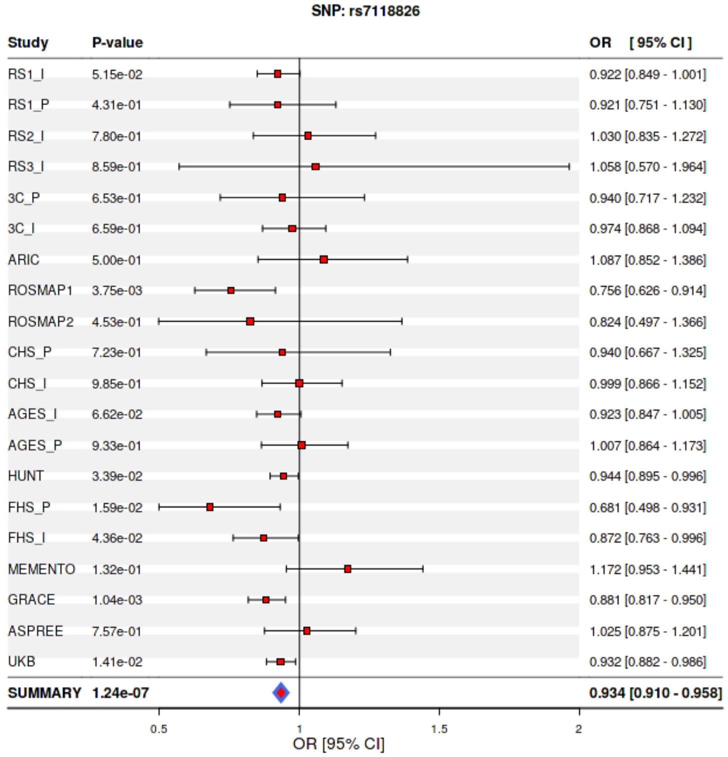

**Figure 8: Forest plot of the locus near ANO3 in ACD GWAS of European ancestry.** For cohort

#### 3 – 1 – 2 SEMA4D locus

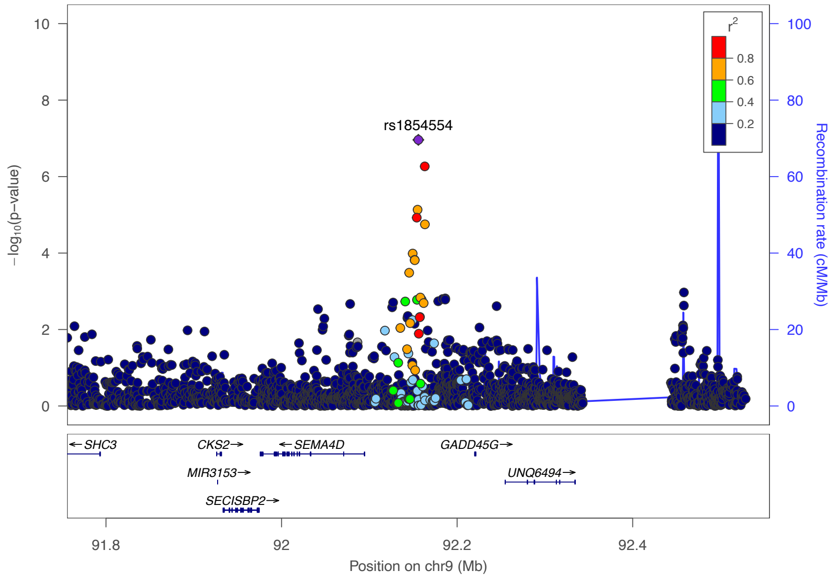

**Figure 9: Regional association plot showing the genomic region containing SEMA4D.** For each SNP, the P-value (log10 scale) of the association with ACD is represented (y-axis, left). The recombination rates (y-axis right), which reflects the local linkage disequilibrium structure is also plotted.

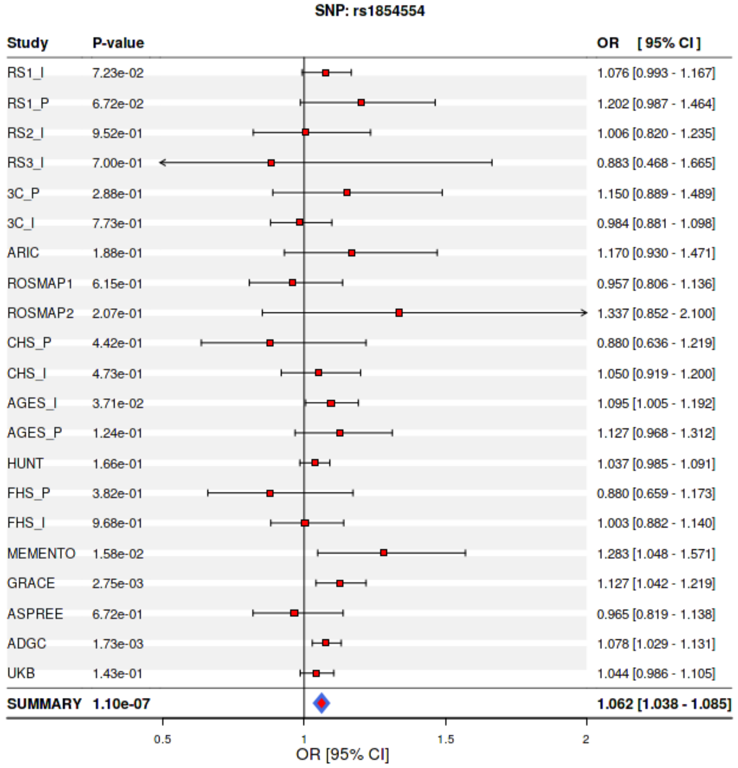

**Figure 10: Forest plot of the locus near SEMA4D in ACD GWAS of European ancestry.** For cohort

#### 3 – 1 – 3 RBFOX1 locus

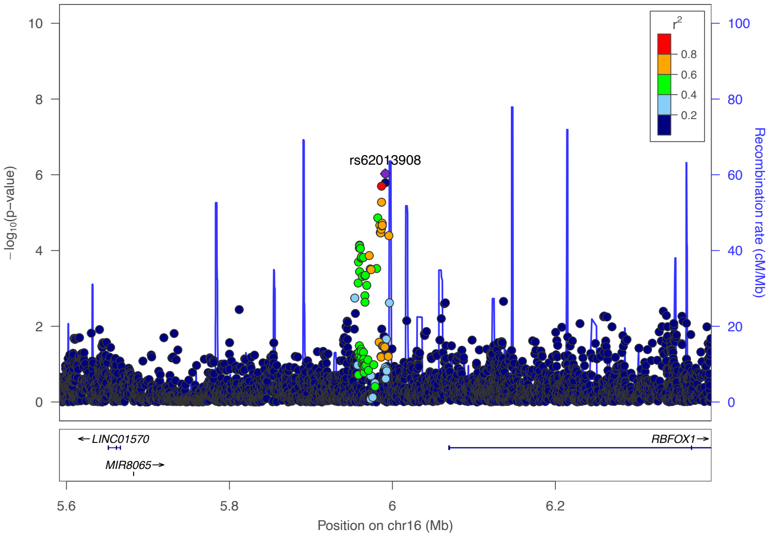

**<u>Figure 11:</u> Regional association plot showing the genomic region containing RBFOX1.** For each SNP, the P-value (log10 scale) of the association with ACD is represented (y-axis, left). The recombination rates (y-axis right), which reflects the local linkage disequilibrium structure is also plotted

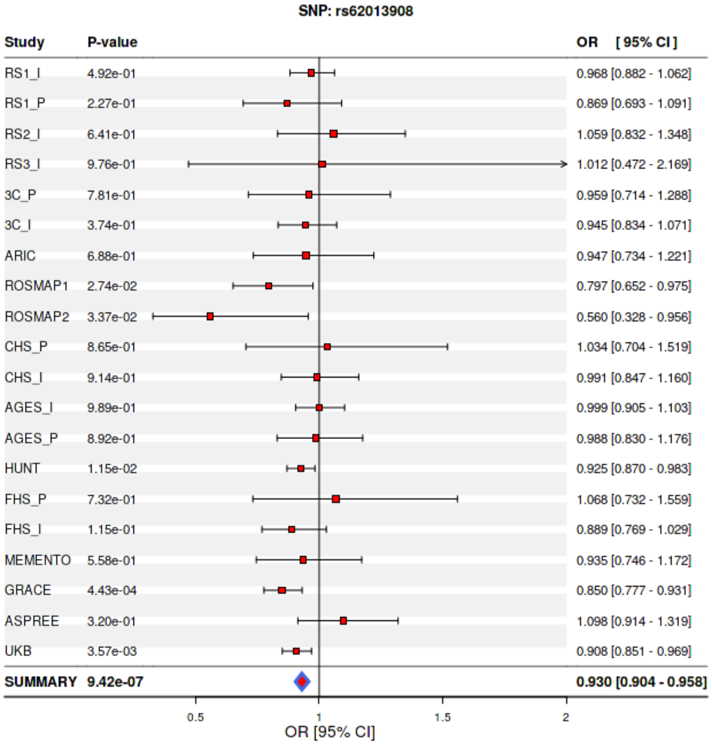

**<u>Figure 12:</u> Forest plot of the locus near RBFOX1 in ACD GWAS of European ancestry.** For cohort

#### 3 – 1 – 4 TRIB1 locus

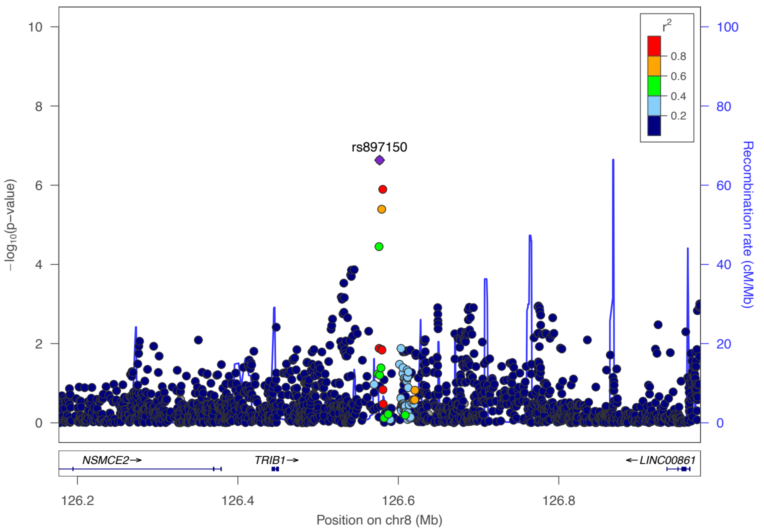

**Figure 13: Regional association plot showing the genomic region containing TRIB1.** For each SNP, the P-value (log10 scale) of the association with ACD is represented (y-axis, left). The recombination rates (y-axis right), which reflects the local linkage disequilibrium structure is also plotted

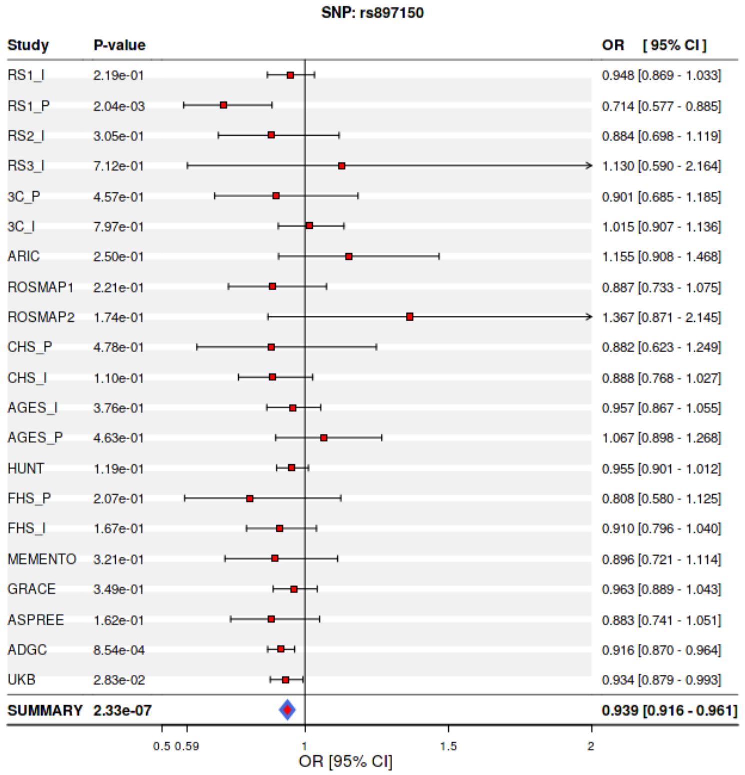

**Figure 12: Forest plot of the locus near TRIB1 in ACD GWAS of European ancestry.** For cohort

#### 3 – 1 – 5 HBEGF locus

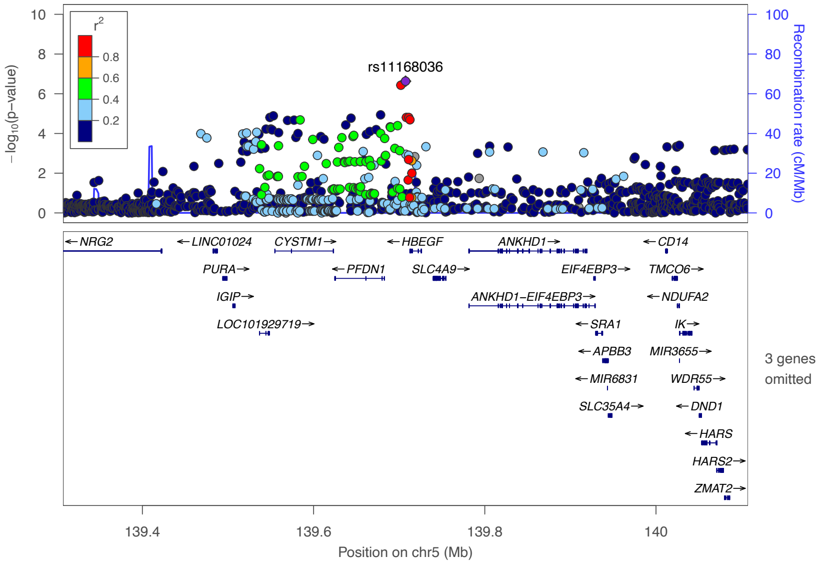

**<u>Figure 15:</u> Regional association plot showing the genomic region containing HBEGF.** For each SNP, the P-value (log10 scale) of the association with ACD is represented (y-axis, left). The recombination rates (y-axis right), which reflects the local linkage disequilibrium structure is also plotted

**<u>Figure 12:</u> Forest plot of the locus near TRIB1 in ACD GWAS of European ancestry.** For cohort

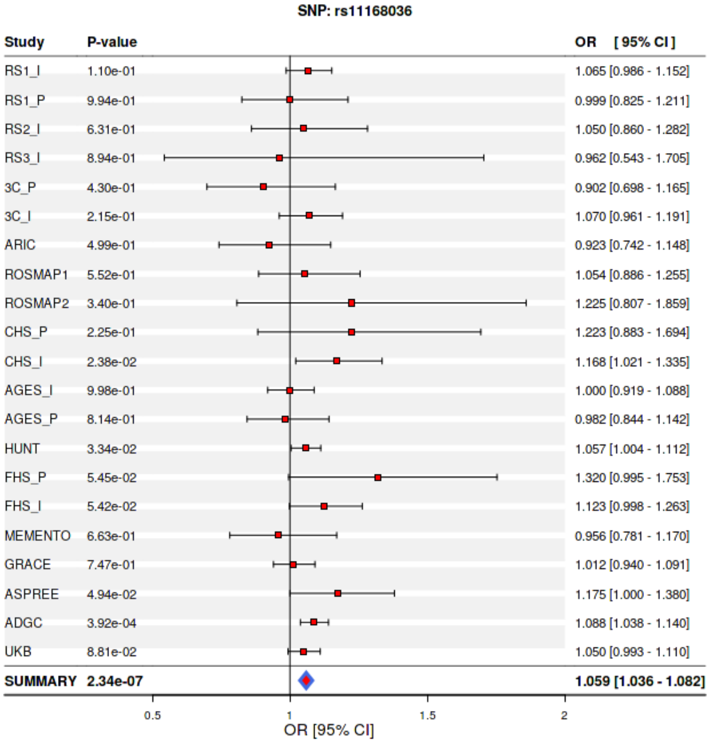

**<u>Figure 16:</u> Forest plot of the locus near HBEGF in ACD GWAS of European ancestry.** For cohort

#### 3 – 1 – 5 ZNF652 locus

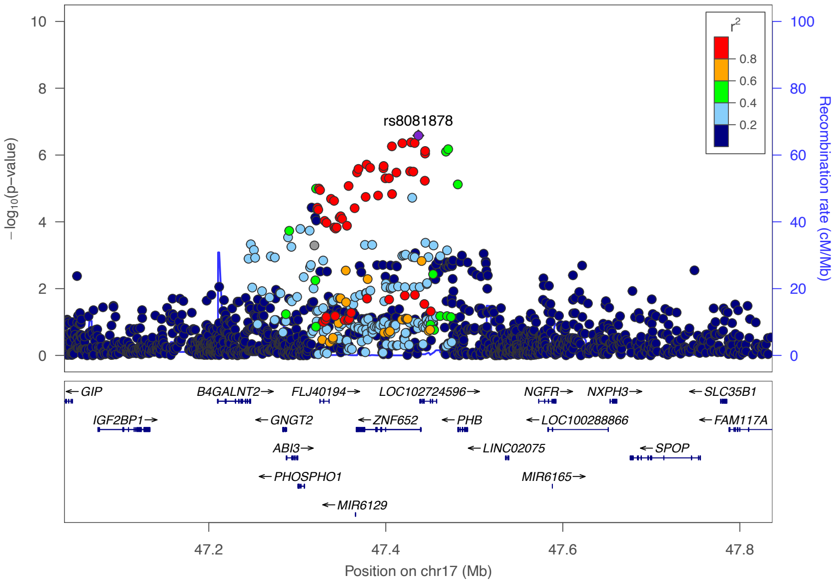

**<u>Figure 17:</u> Regional association plot showing the genomic region containing ZNF652.** For each SNP, the P-value (log10 scale) of the association with ACD is represented (y-axis, left). The recombination rates (y-axis right), which reflects the local linkage disequilibrium structure is also plotted

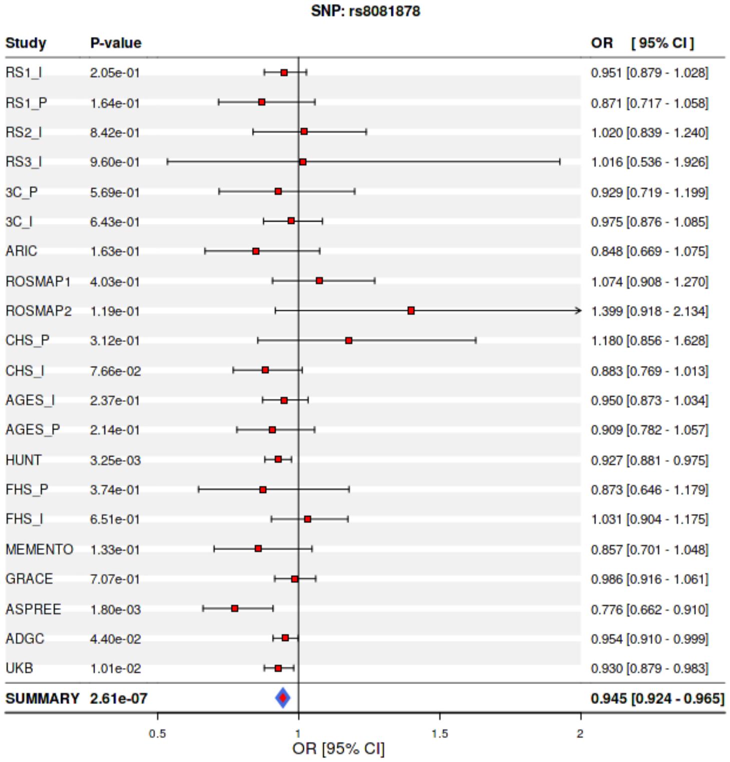

**<u>Figure 18:</u> Forest plot of the locus near ZNF652 in ACD GWAS of European ancestry.** For cohort

#### 3 – 1 – 6 SREBF1 locus

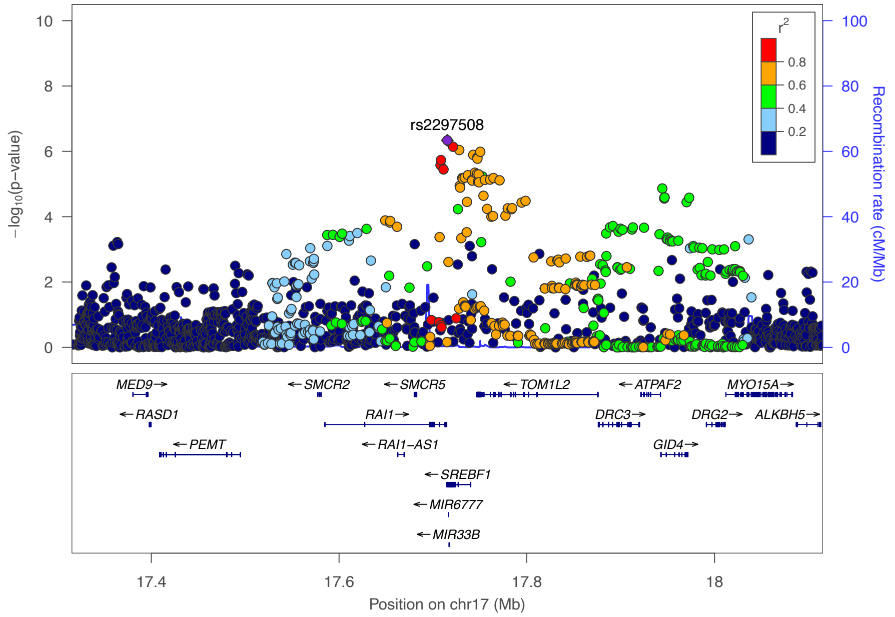

**<u>Figure 19:</u> Regional association plot showing the genomic region containing SREBF1.** For each SNP, the P-value (log10 scale) of the association with ACD is represented (y-axis, left). The recombination rates (y-axis right), which reflects the local linkage disequilibrium structure is also plotted

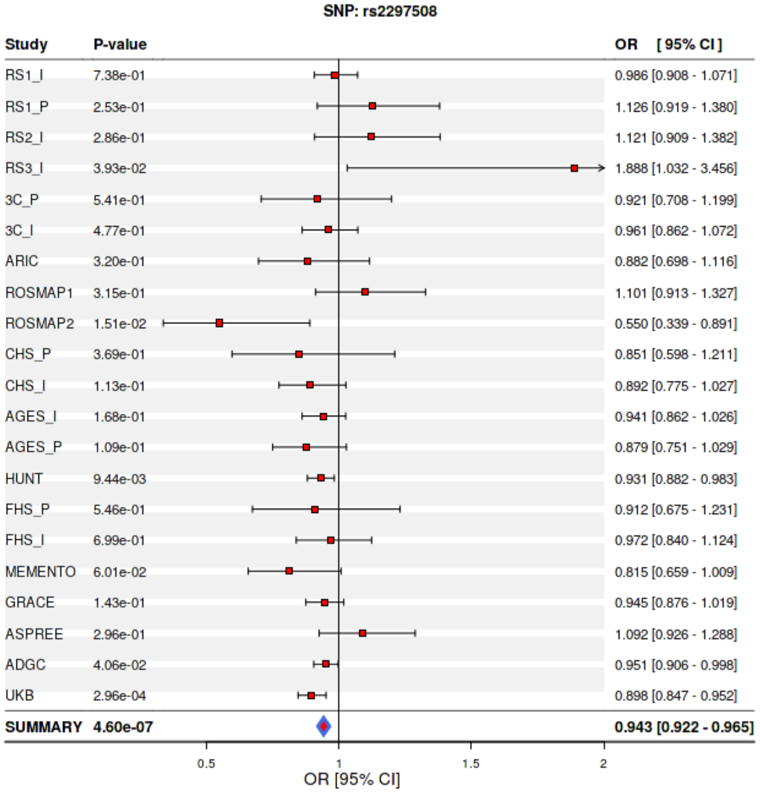

**<u>Figure 20:</u> Forest plot of the locus near SREBF1 in ACD GWAS of European ancestry.** For cohort

#### 3 – 1 – 7 INPP5D locus

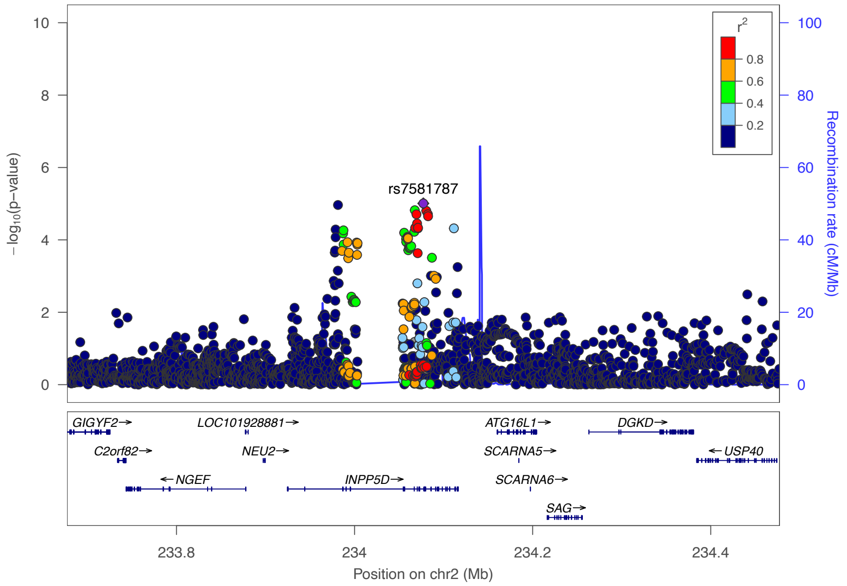

**Figure 21: Regional association plot showing the genomic region containing INPP5D.** For each SNP, the P-value (log10 scale) of the association with ACD is represented (y-axis, left). The recombination rates (y-axis right), which reflects the local linkage disequilibrium structure is also plotted

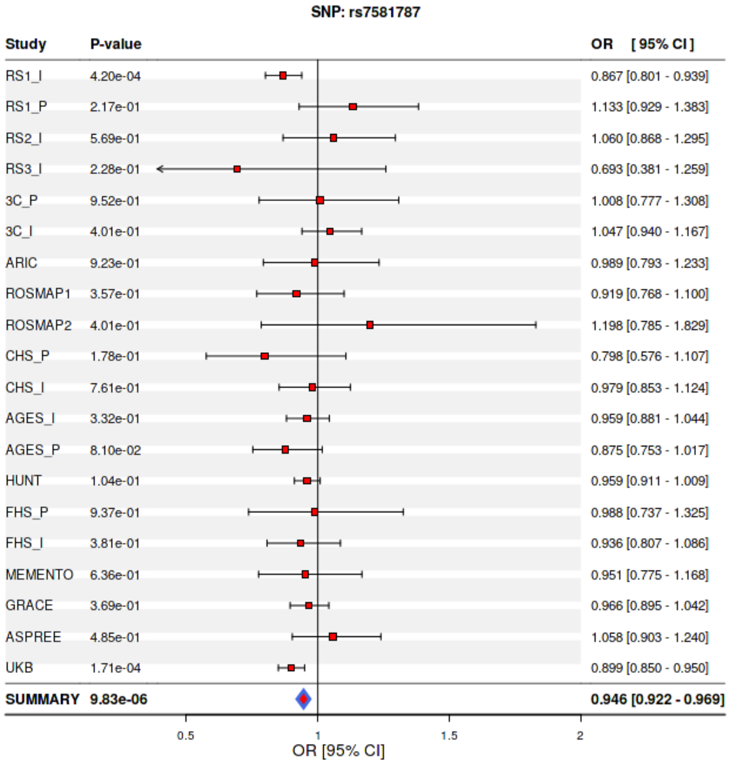

**Figure 22: Forest plot of the locus near INPP5D in ACD GWAS of European ancestry.** For cohort

#### 3 – 2 Vascular dementia suggestive variants

##### 3 – 2 – 1 SPRY2 locus

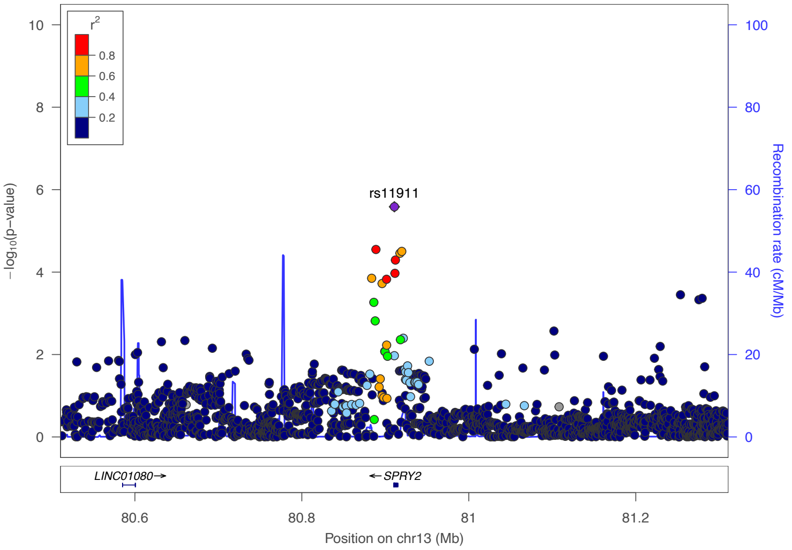

**Figure 23: Regional association plot showing the genomic region containing SPRY2.** For each SNP, the P-value (log10 scale) of the association with VaD is represented (y-axis, left). The recombination rates (y-axis right), which reflects the local linkage disequilibrium structure is also plotted

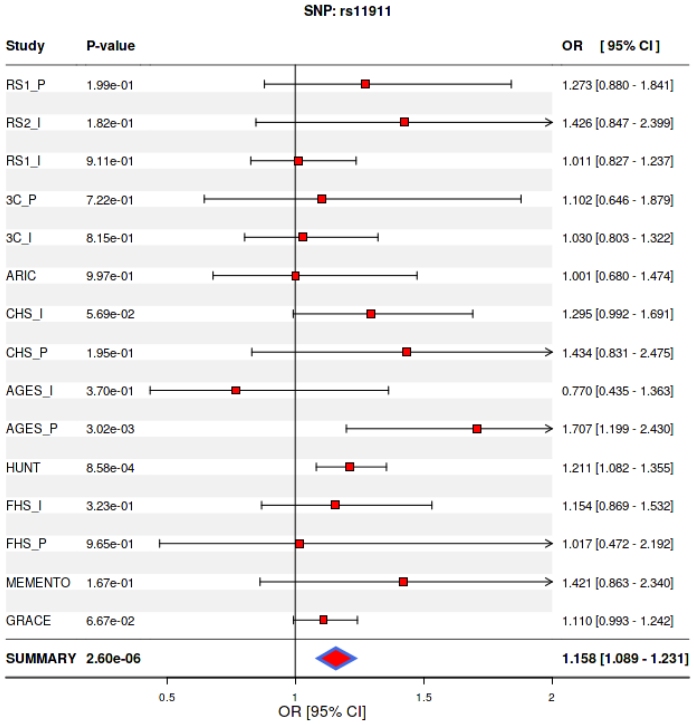

**Figure 24: Forest plot of the locus near SPRY2 in ACD GWAS of European ancestry.** For cohort

#### 3 – 2 – 1 SEMA6D locus

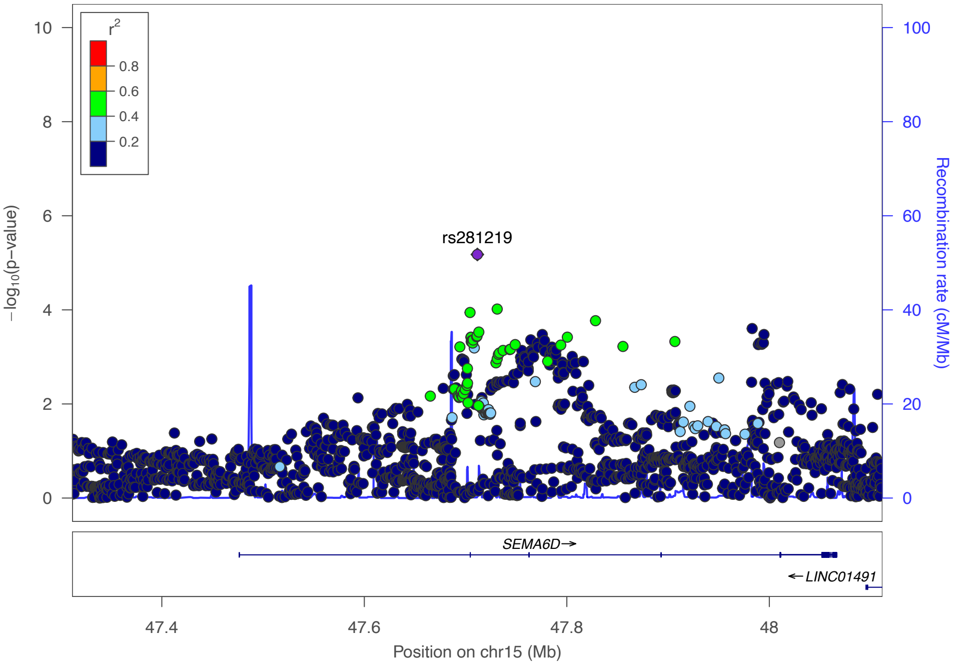

**<u>Figure 23:</u> Regional association plot showing the genomic region containing SPRY2.** For each SNP, the P-value (log10 scale) of the association with VaD is represented (y-axis, left). The recombination rates (y-axis right), which reflects the local linkage disequilibrium structure is also plotted

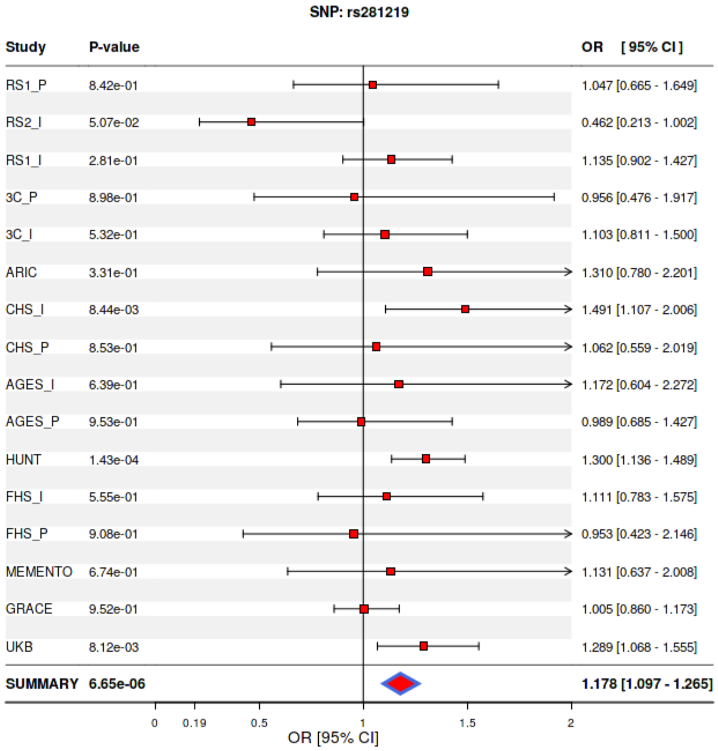

**<u>Figure 24:</u> Forest plot of the locus near SPRY2 in ACD GWAS of European ancestry.** For cohort

#### 3 – 2 – 1 SCARB1 locus

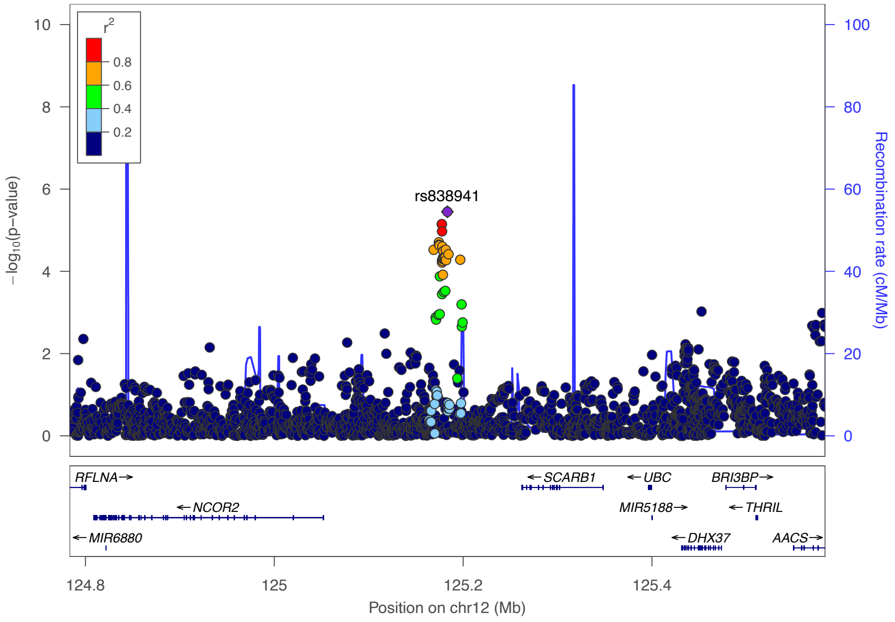

**Figure 23:** **Regional association plot showing the genomic region containing SPRY2.** For each SNP, the P-value (log10 scale) of the association with VaD is represented (y-axis, left). The recombination rates (y-axis right), which reflects the local linkage disequilibrium structure is also plotted

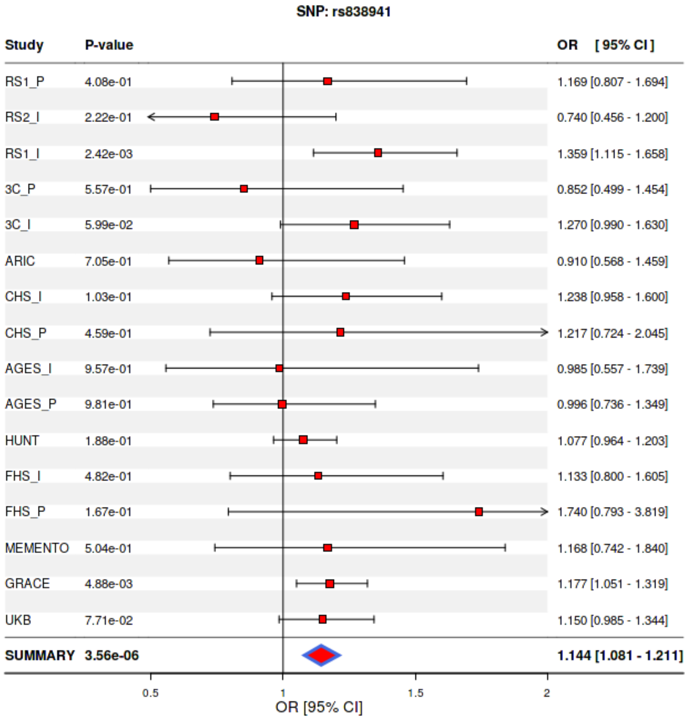

**Figure 24:** **Forest plot of the locus near SPRY2 in ACD GWAS of European ancestry.** For cohort

#### 3 – 2 – 1 PSMA3 locus

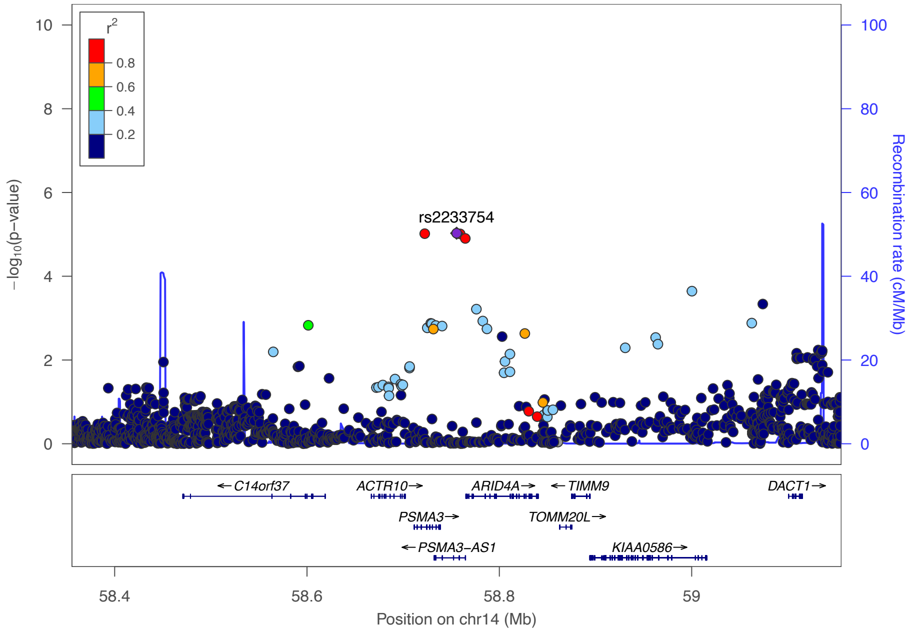

**Figure 23: Regional association plot showing the genomic region containing SPRY2.** For each SNP, the P-value (log10 scale) of the association with VaD is represented (y-axis, left). The recombination rates (y-axis right), which reflects the local linkage disequilibrium structure is also plotted

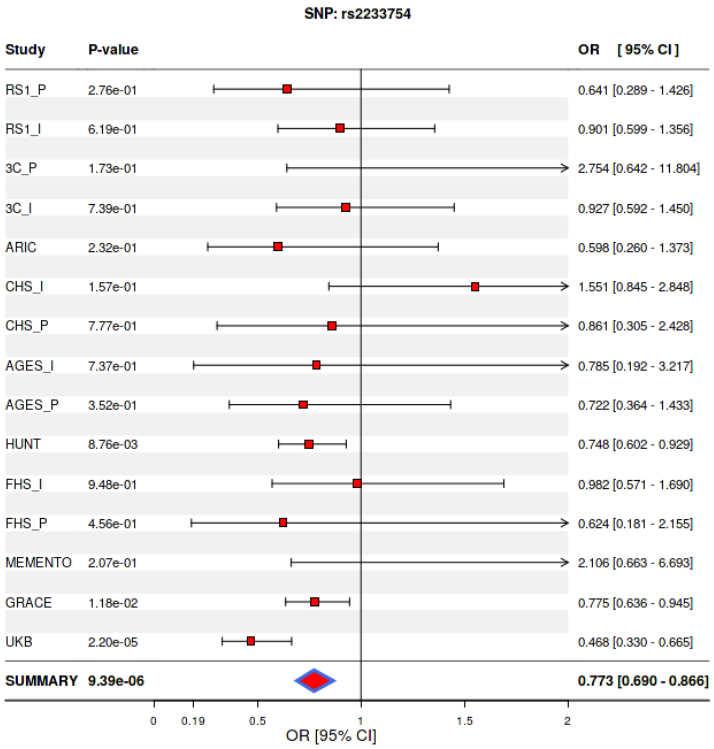

**Figure 24: Forest plot of the locus near SPRY2 in ACD GWAS of European ancestry.** For cohort

#### 3 – 2 – 1 LINC02113 locus

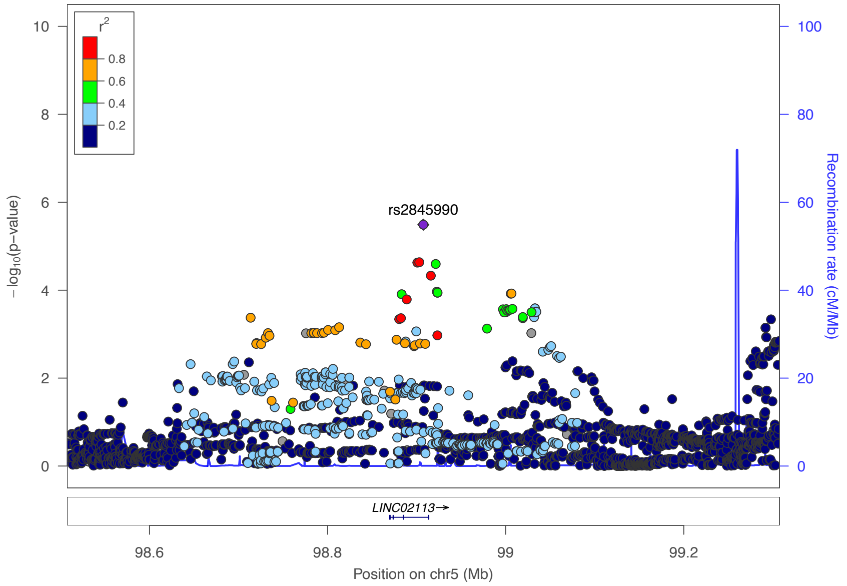

**<u>Figure 23:</u> Regional association plot showing the genomic region containing SPRY2.** For each SNP, the P-value (log10 scale) of the association with VaD is represented (y-axis, left). The recombination rates (y-axis right), which reflects the local linkage disequilibrium structure is also plotted

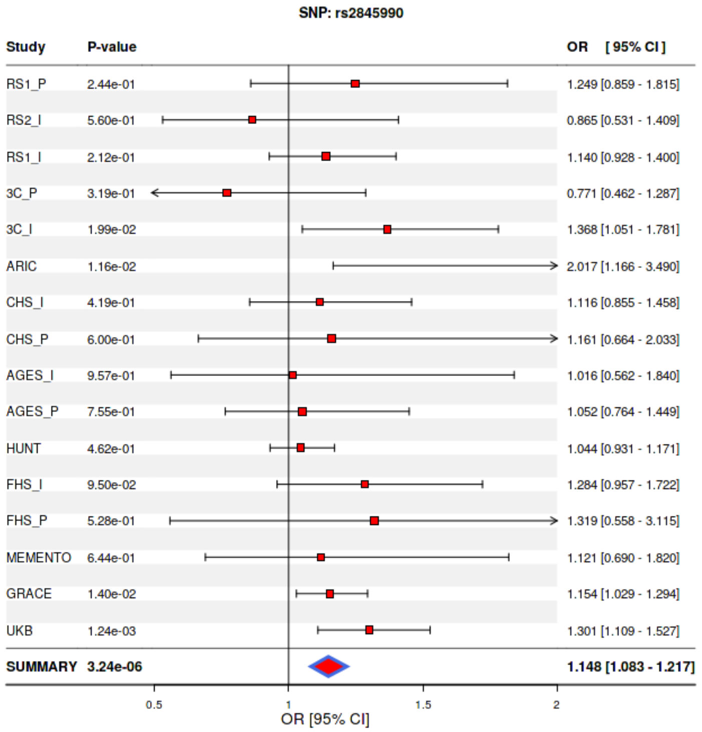

**<u>Figure 24:</u> Forest plot of the locus near SPRY2 in ACD GWAS of European ancestry.** For cohort

#### 3 – 2 – 1 GIP locus

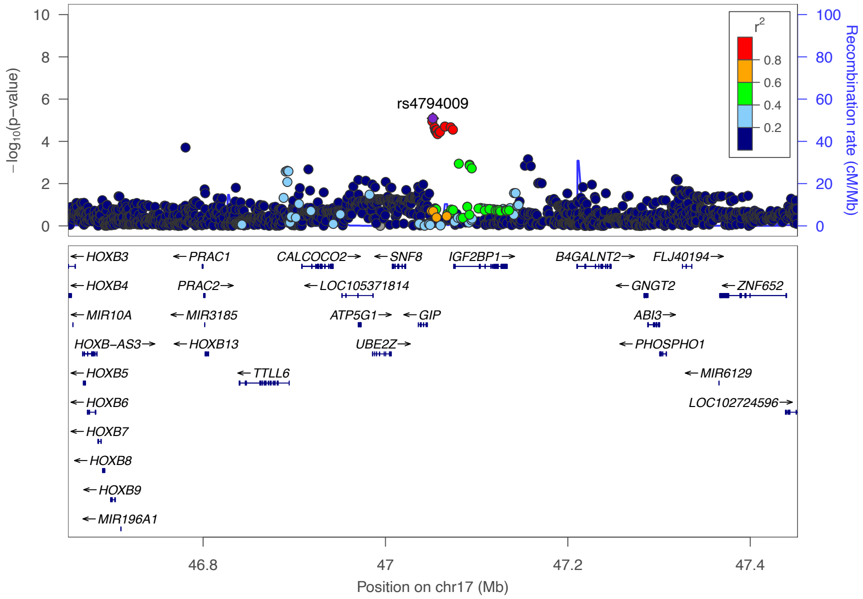

**Figure 23: Regional association plot showing the genomic region containing SPRY2.** For each SNP, the P-value (log10 scale) of the association with VaD is represented (y-axis, left). The recombination rates (y-axis right), which reflects the local linkage disequilibrium structure is also plotted

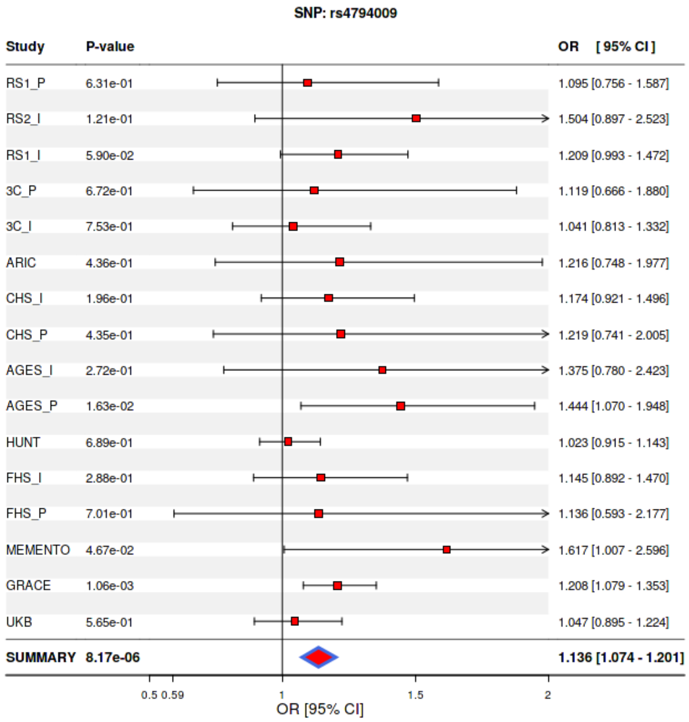

**Figure 24: Forest plot of the locus near SPRY2 in ACD GWAS of European ancestry.** For cohort

#### 3 – 2 – 1 DOK5 locus

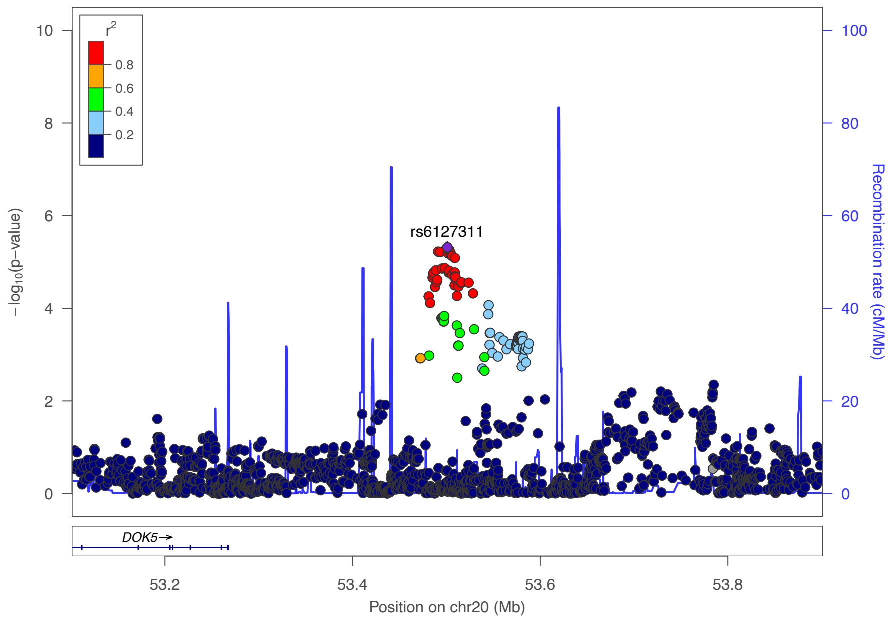

**Figure 23: Regional association plot showing the genomic region containing SPRY2.** For each SNP, the P-value (log10 scale) of the association with VaD is represented (y-axis, left). The recombination rates (y-axis right), which reflects the local linkage disequilibrium structure is also plotted

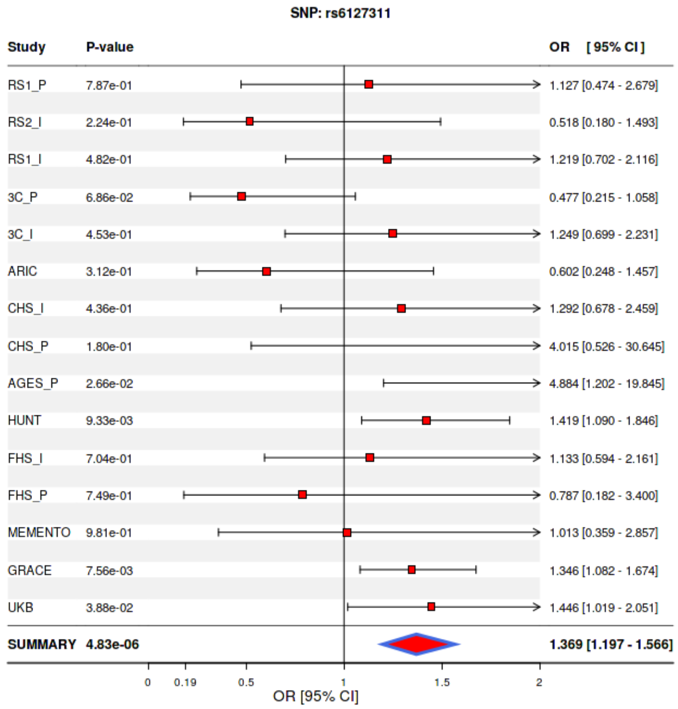

**Figure 24: Forest plot of the locus near SPRY2 in ACD GWAS of European ancestry.** For cohort

#### 3 – 2 – 1 GALNT18 locus

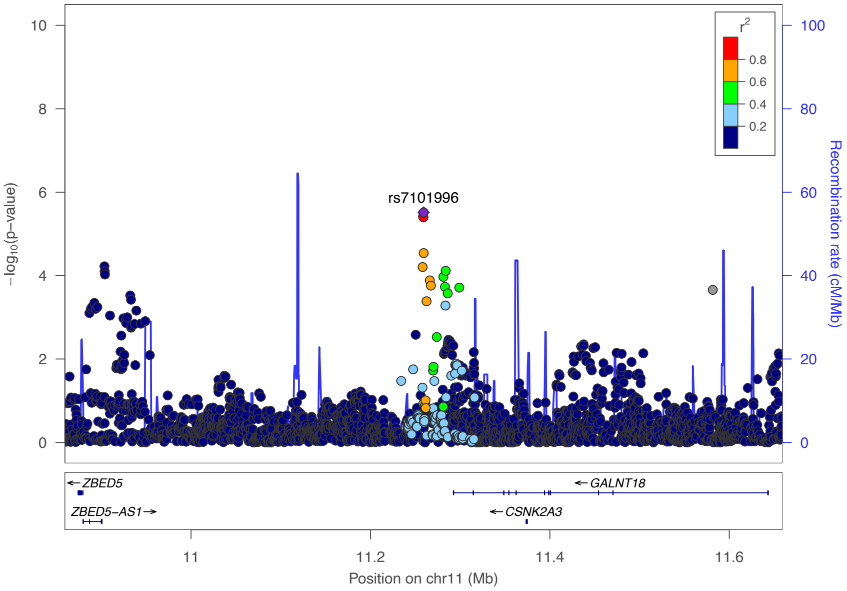

**<u>Figure 23:</u> Regional association plot showing the genomic region containing SPRY2.** For each SNP, the P-value (log10 scale) of the association with VaD is represented (y-axis, left). The recombination rates (y-axis right), which reflects the local linkage disequilibrium structure is also plotted

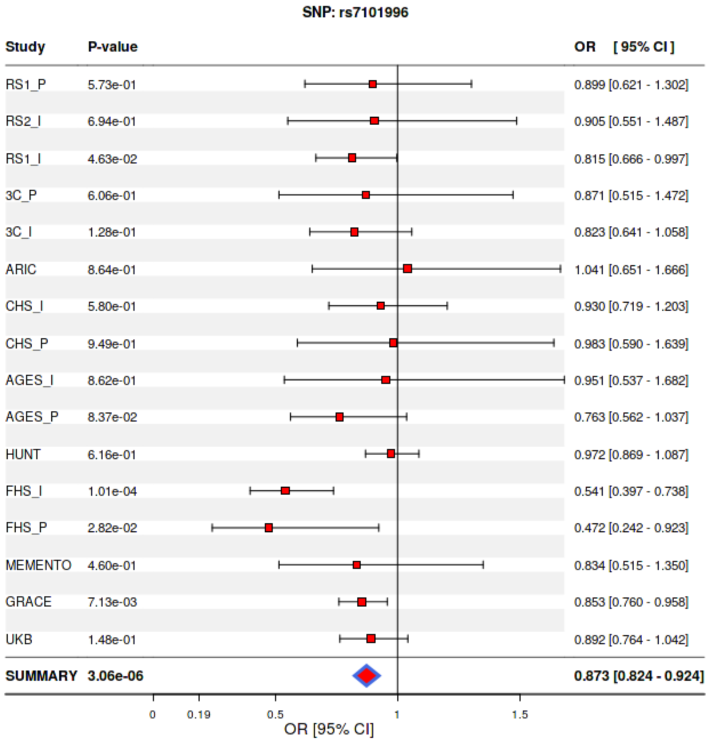

**<u>Figure 24:</u> Forest plot of the locus near SPRY2 in ACD GWAS of European ancestry.** For cohort

#### 3 – 2 – 1 WAC locus

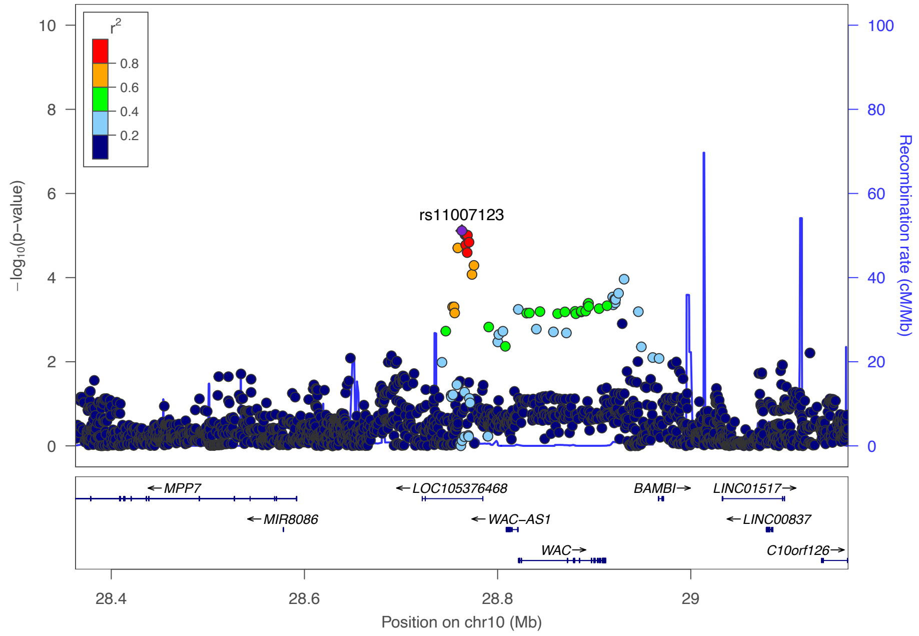

**<u>Figure 23:</u> Regional association plot showing the genomic region containing SPRY2.** For each SNP, the P-value (log10 scale) of the association with VaD is represented (y-axis, left). The recombination rates (y-axis right), which reflects the local linkage disequilibrium structure is also plotted

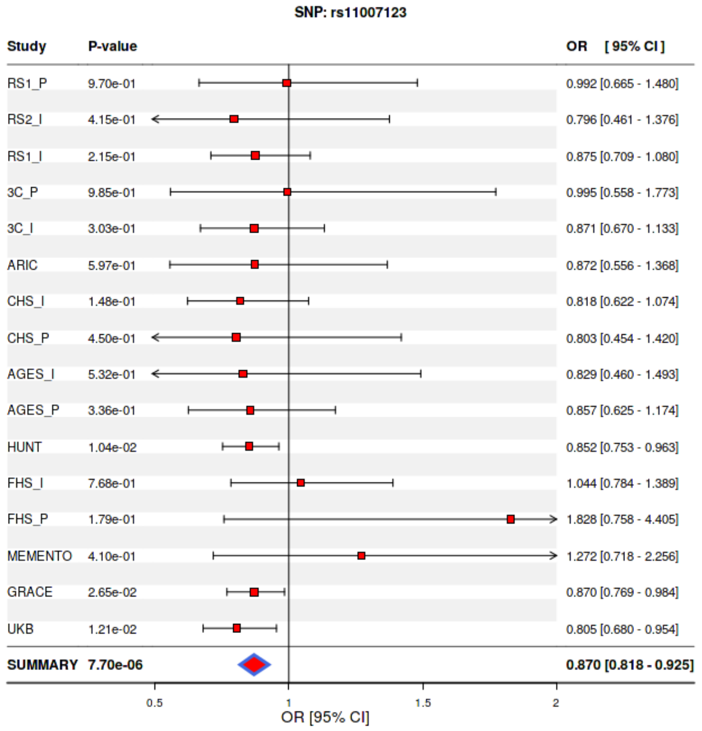

**<u>Figure 24:</u> Forest plot of the locus near SPRY2 in ACD GWAS of European ancestry.** For cohort

#### 3 – 2 – 1 ERBB4 locus

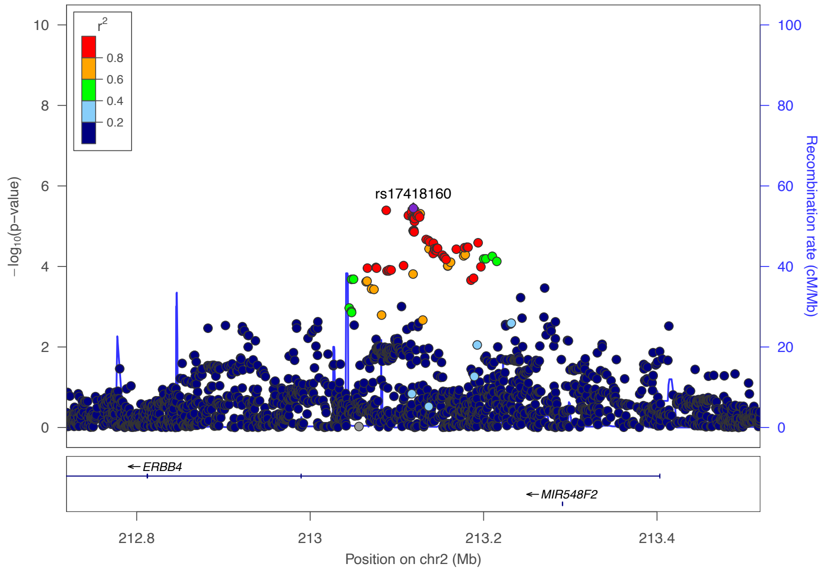

**<u>Figure 23:</u> Regional association plot showing the genomic region containing SPRY2.** For each SNP, the P-value (log10 scale) of the association with VaD is represented (y-axis, left). The recombination rates (y-axis right), which reflects the local linkage disequilibrium structure is also plotted

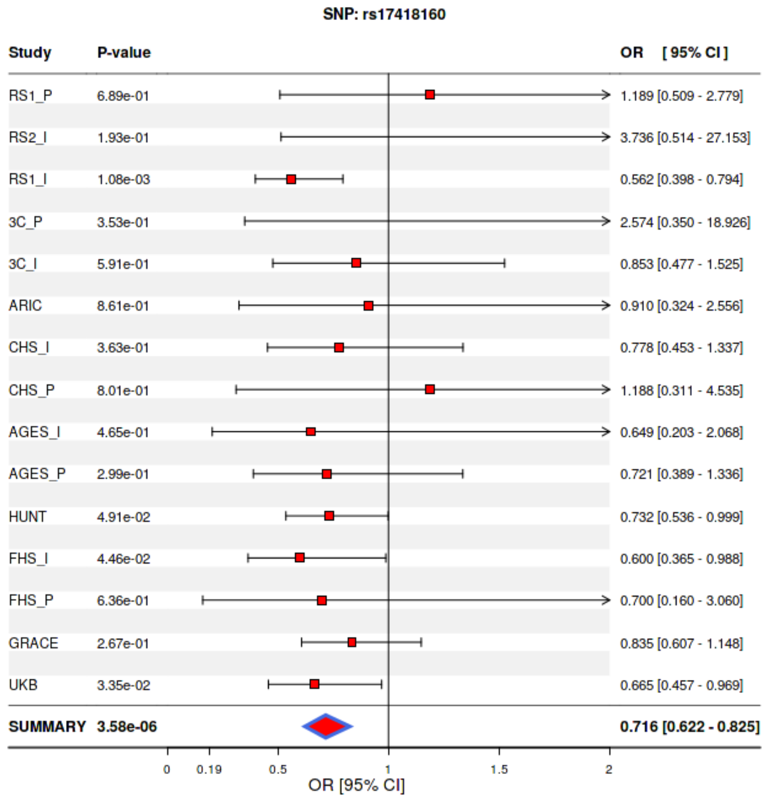

**<u>Figure 24:</u> Forest plot of the locus near SPRY2 in ACD GWAS of European ancestry.** For cohort

#### 3 – 2 – 1 PRKCE locus

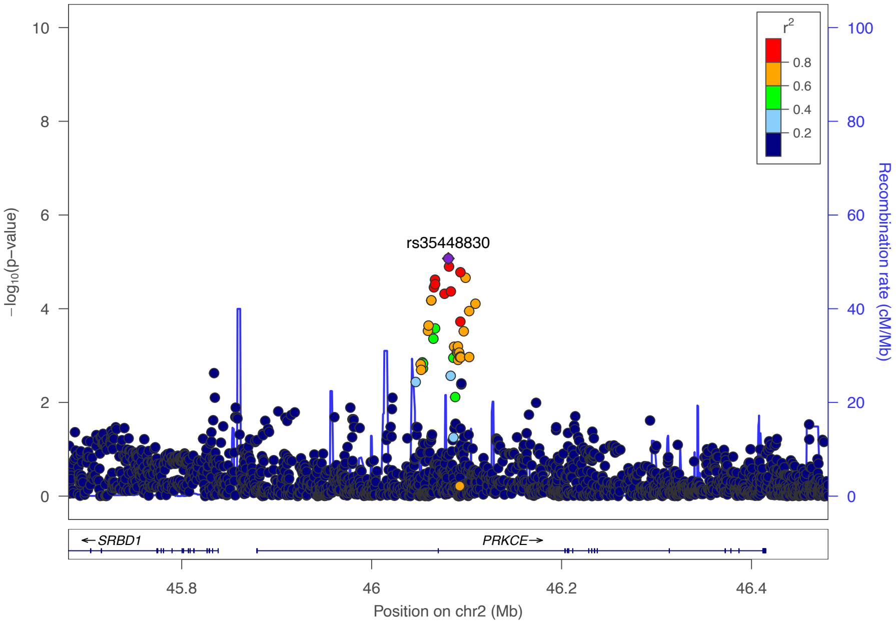

**Figure 23: Regional association plot showing the genomic region containing SPRY2.** For each SNP, the P-value (log10 scale) of the association with VaD is represented (y-axis, left). The recombination rates (y-axis right), which reflects the local linkage disequilibrium structure is also plotted

#### 3 – 2 – 1 PHACTR3 locus

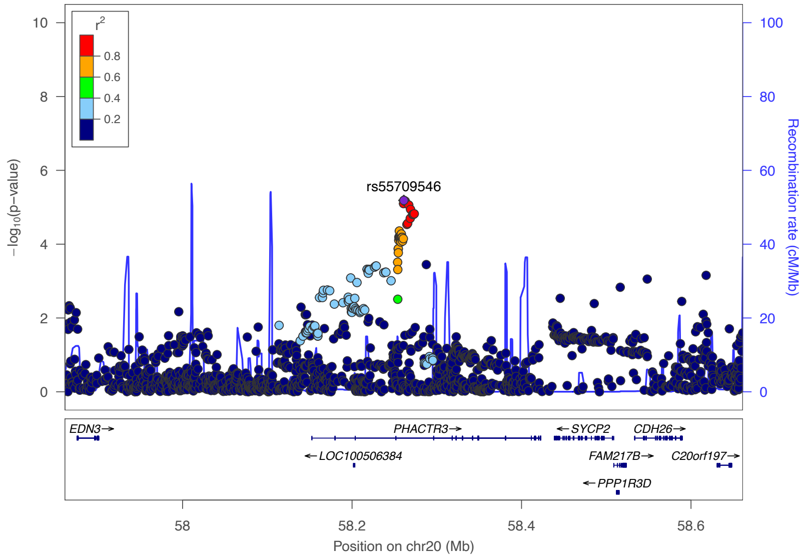

**Figure 23: Regional association plot showing the genomic region containing SPRY2.** For each SNP, the P-value (log10 scale) of the association with VaD is represented (y-axis, left). The recombination rates (y-axis right), which reflects the local linkage disequilibrium structure is also plotted

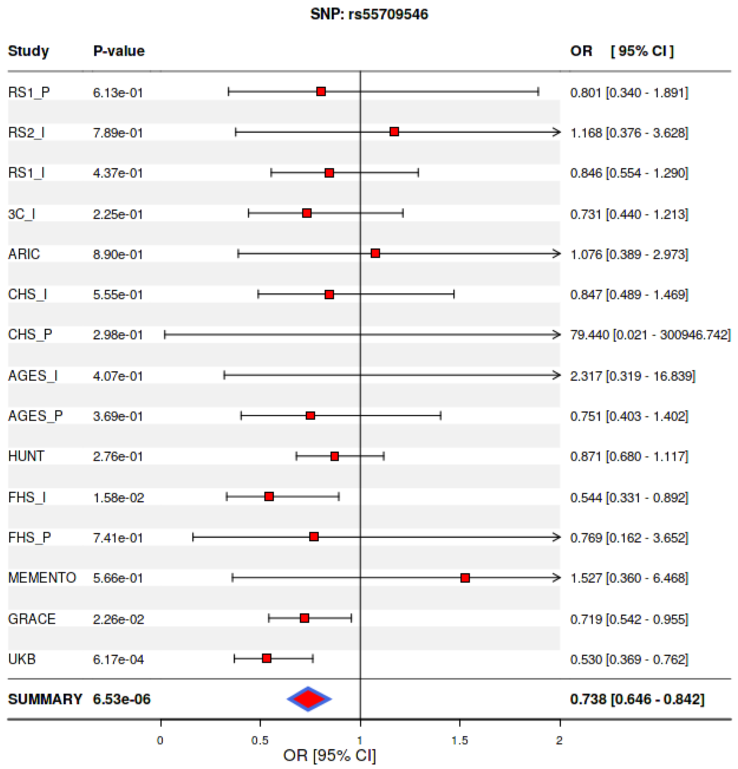

**<u>Figure 24:</u> Forest plot of the locus near SPRY2 in ACD GWAS of European ancestry.** For cohort

#### 3 – 2 – 1 AJAP1 locus

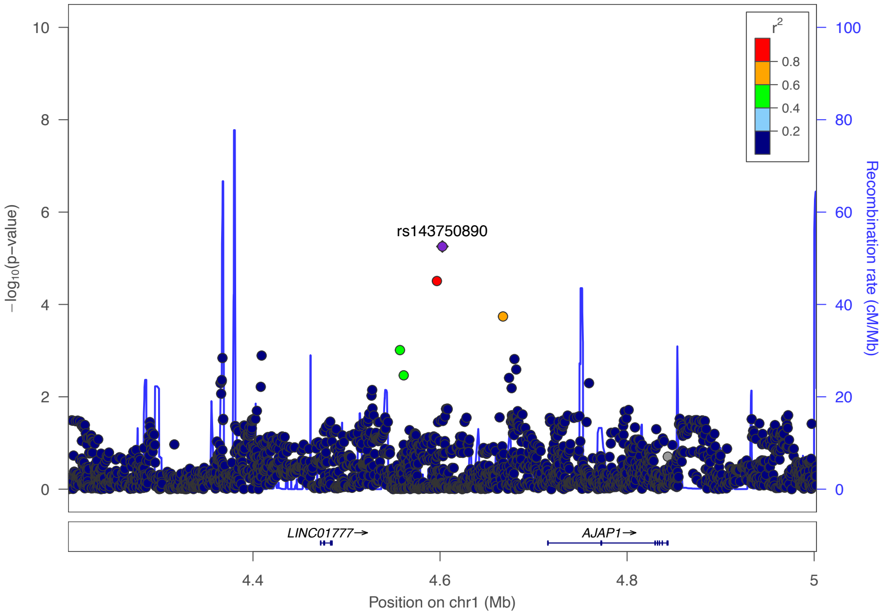

**<u>Figure 23:</u> Regional association plot showing the genomic region containing SPRY2.** For each SNP, the P-value (log10 scale) of the association with VaD is represented (y-axis, left). The recombination rates (y-axis right), which reflects the local linkage disequilibrium structure is also plotted

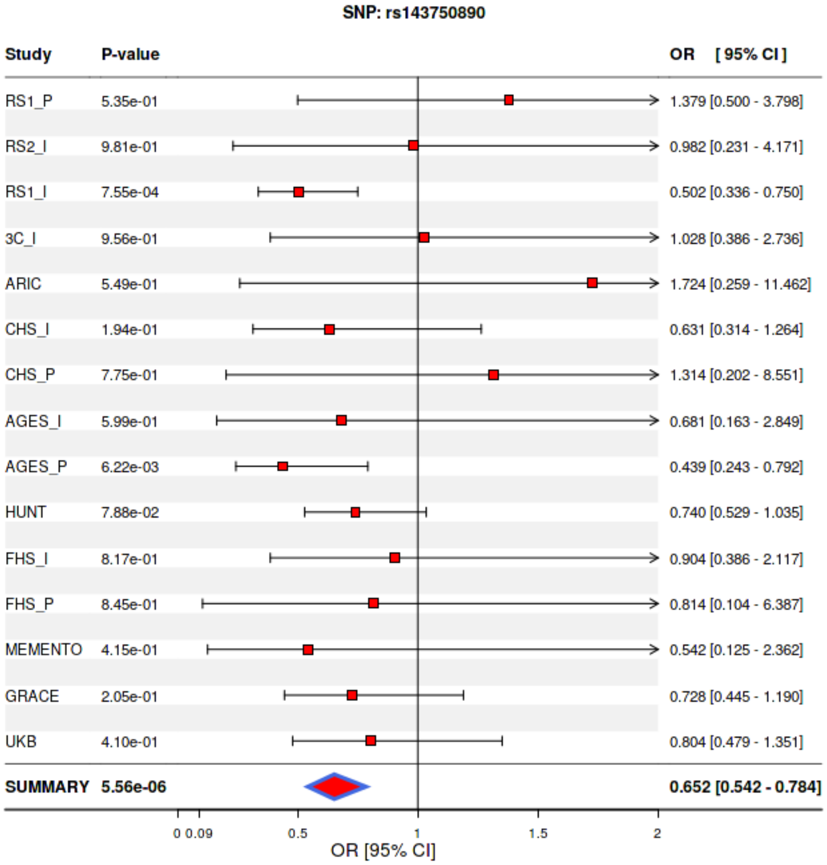

<u>**Figure 24:**</u> **Forest plot of the locus near SPRY2 in ACD GWAS of European ancestry.** For cohort

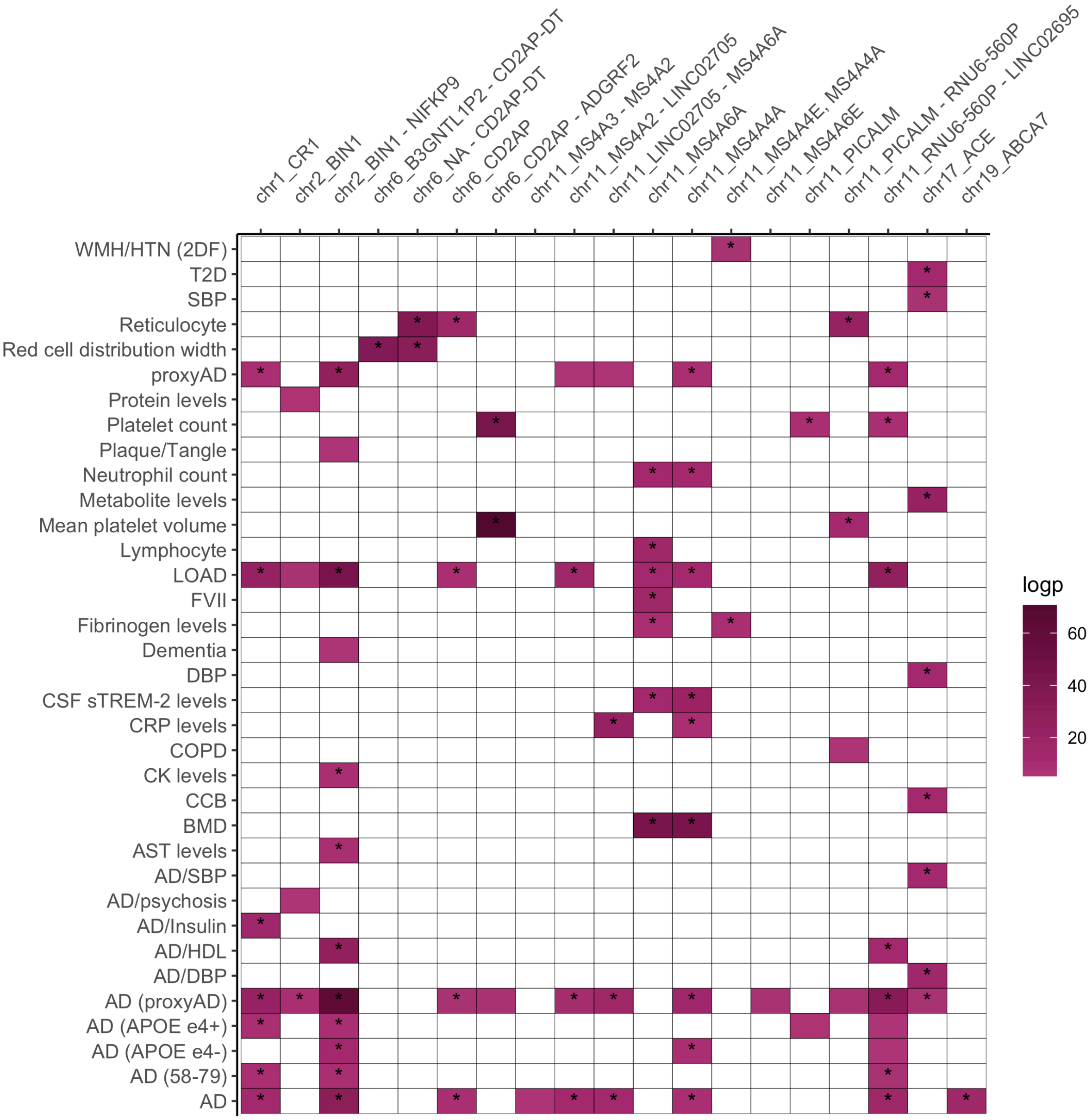

**Figure 25:** **Variant level overlap with complex disease traits**

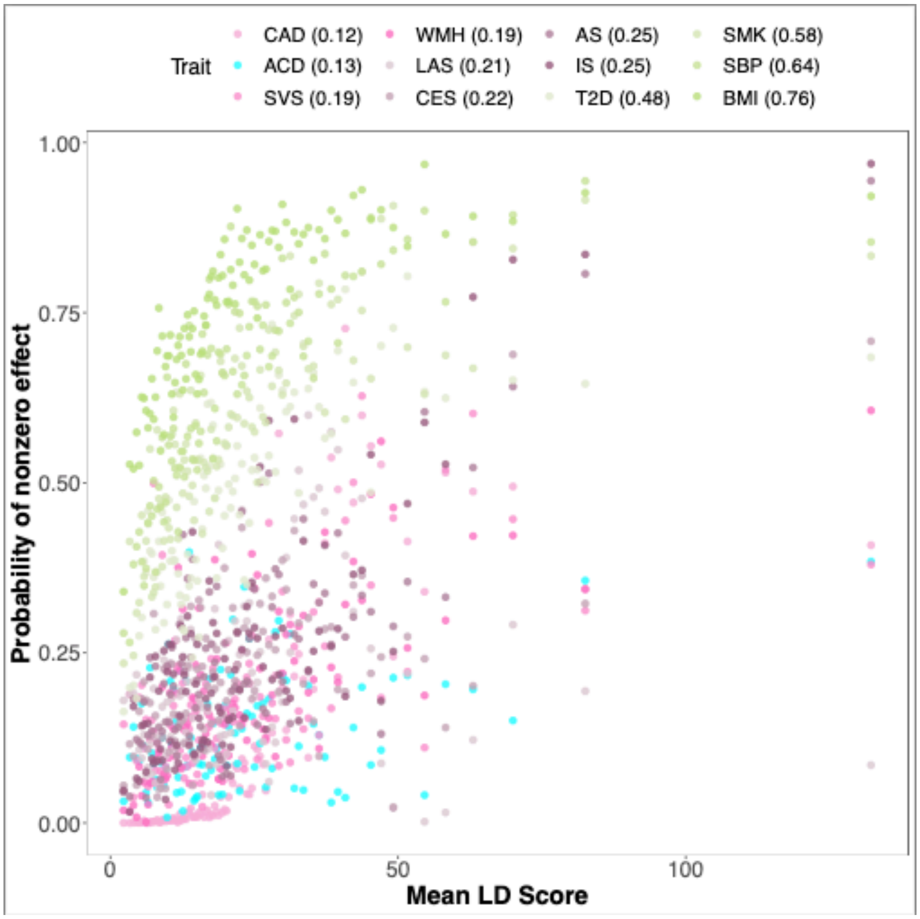

**Figure 26:** Non-zero effect distribution of ACD and other complex disease traits

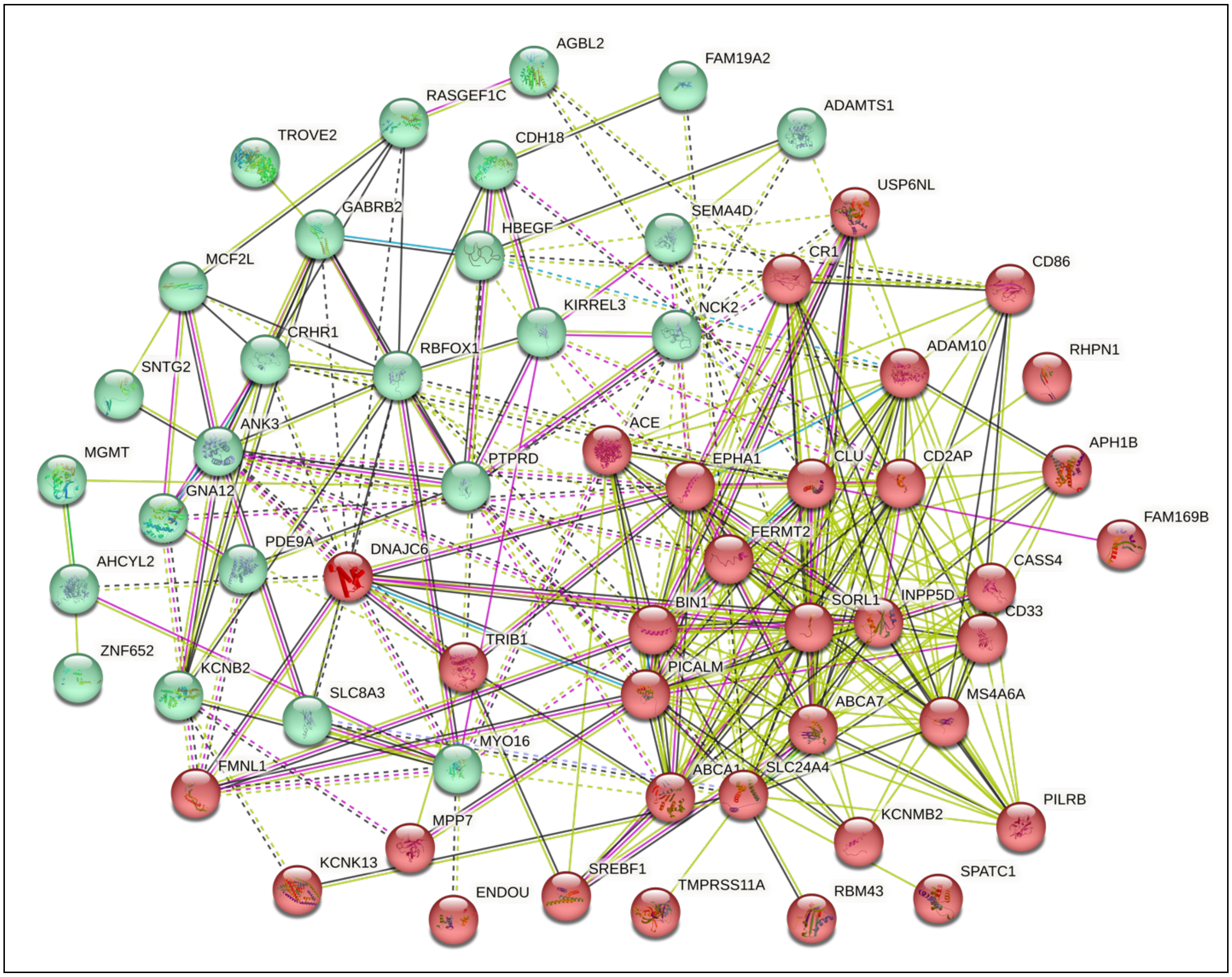

**Figure 27**: **STRING clustering of ACD genes.** Hierarchical clustering of ACD in two groups provides insights into the potential interactions between prominent genes. The red cluster is highly enriched in known AD-related genes while the green cluster mostly included our ACD suggestive genes.

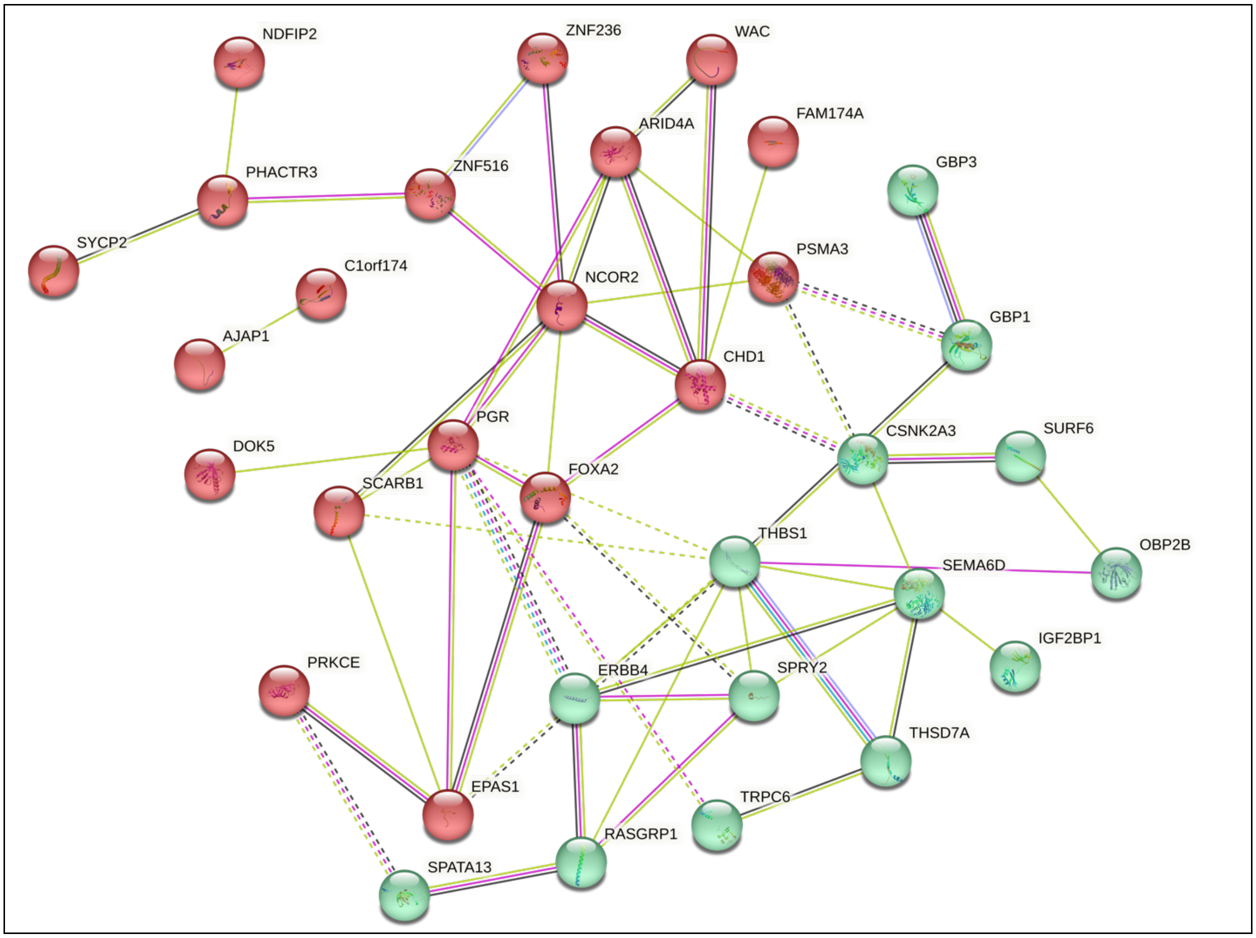

**Figure 28**: **STRING clustering of VaD genes.**

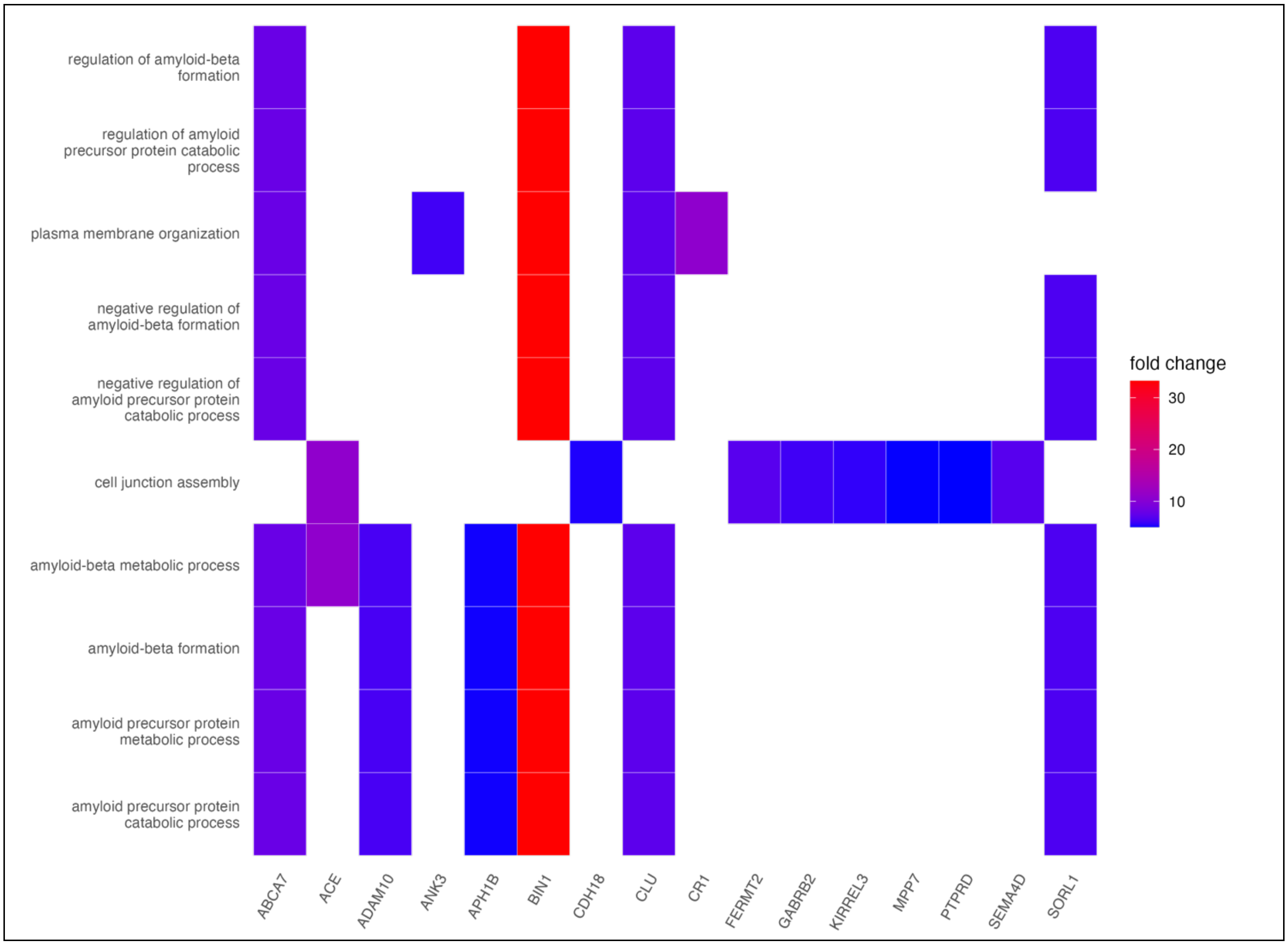

**Figure 29**: **ACD Gene Ontology (GO) analysis, Biological Pathways.**

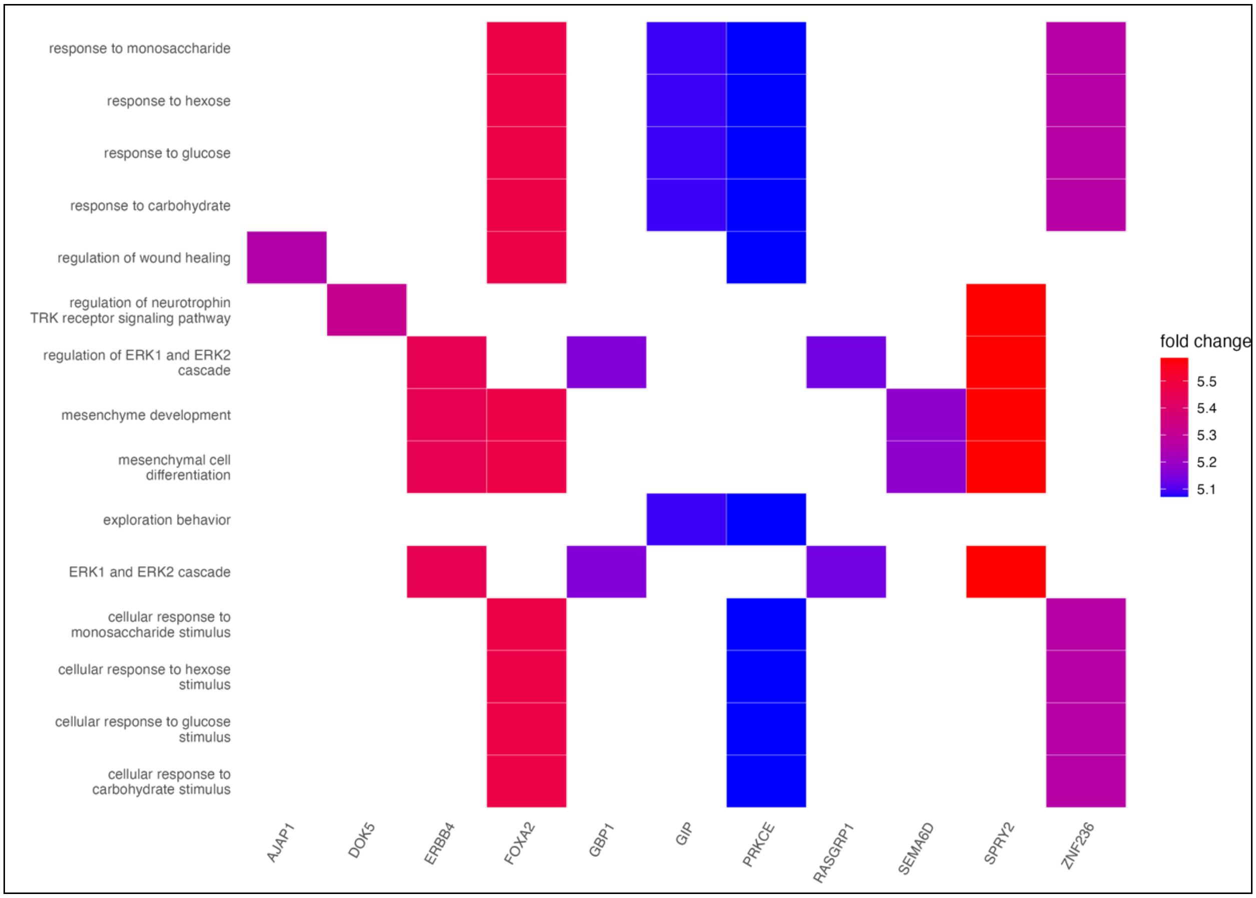

**Figure 30**: **VaD Gene Ontology (GO) analysis, Biological Pathways.**

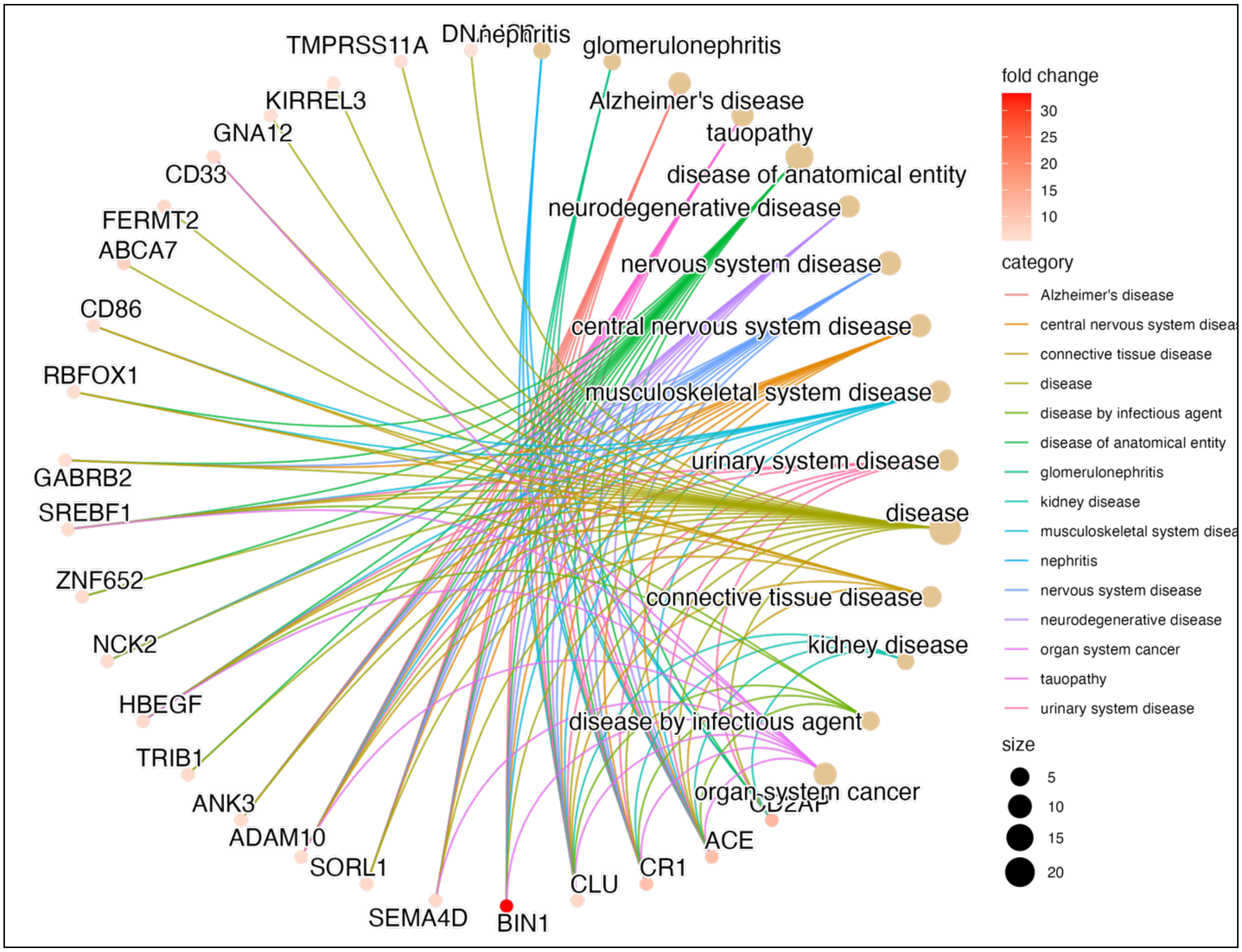

**Figure 31**: **ACD disease association analysis**

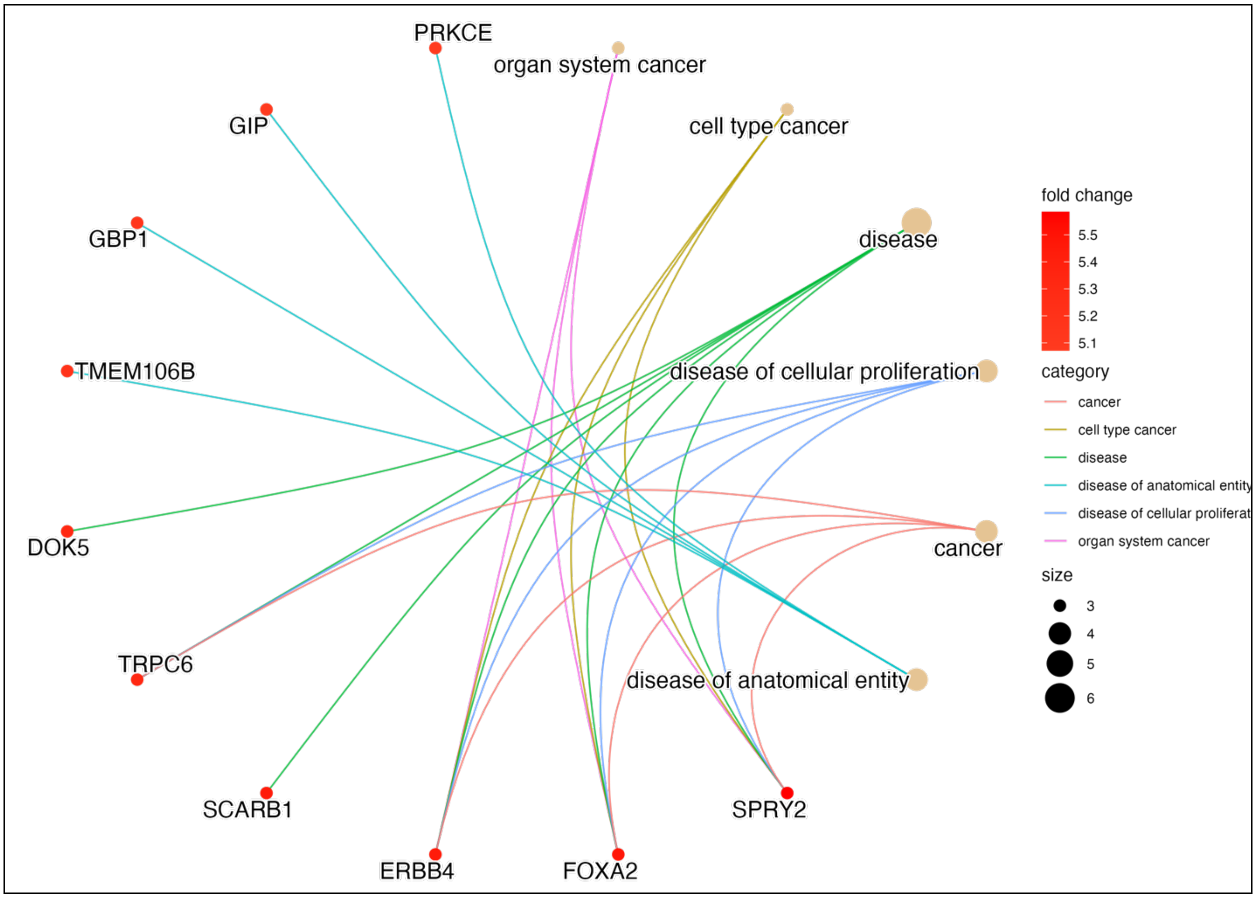

**Figure 32**: **VaD disease association analysis.**

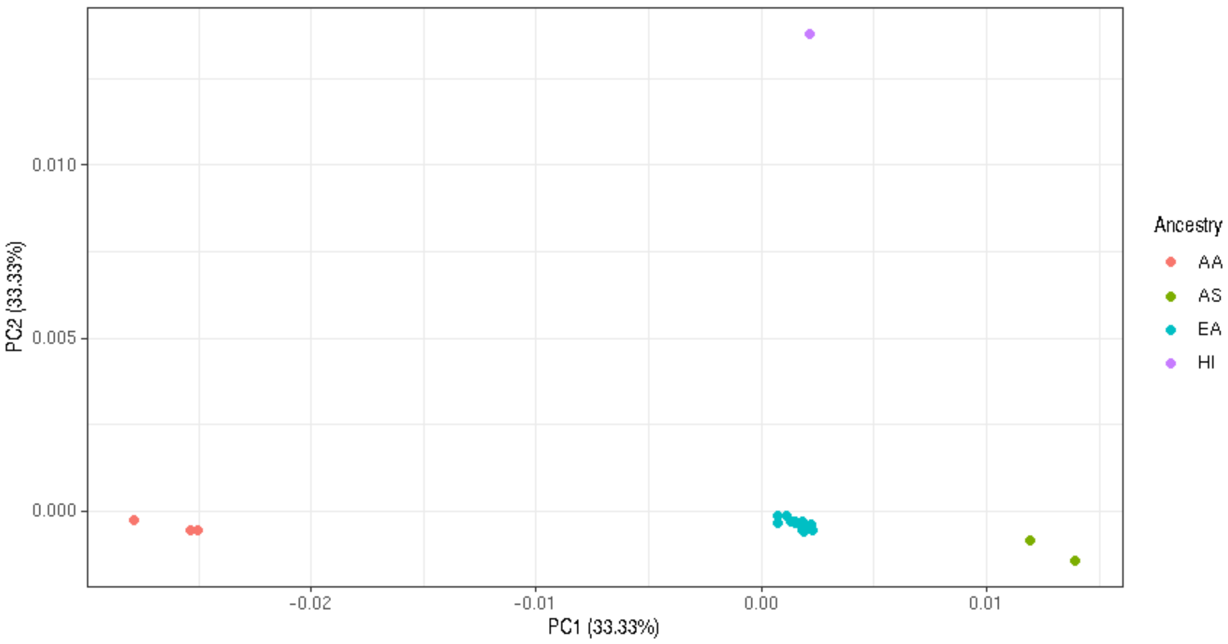

**Figure 33**: **Principal components generated using the mean allele frequency difference between studies.**
Principal components (PCs) are generated using MR-MEGA from a matrix of mean pairwise allele frequency differences between cohorts (total n=). Color denotes self-reported ancestry for each cohort. Selected outliers are labeled with cohort name. Three PCs were chosen as per author recommendations, and, as shown, are sufficient to separate self- reported ancestry groups.
